# Lineage recording in monoclonal gastruloids reveals heritable modes of early development

**DOI:** 10.1101/2025.05.23.655664

**Authors:** Samuel G. Regalado, Chengxiang Qiu, Sanjay Kottapalli, Beth K. Martin, Wei Chen, Hanna Liao, Haedong Kim, Xiaoyi Li, Jean-Benoît Lalanne, Nobuhiko Hamazaki, Silvia Domcke, Junhong Choi, Jay Shendure

## Abstract

Mammalian stem cells possess a remarkable capacity for self-organization, a property that underlies increasingly sophisticated *in vitro* models of early development. However, even under carefully controlled conditions, stem cell-derived models exhibit substantial “inter-individual” heterogeneity. Focusing on gastruloids, a powerful model of the early posterior embryo^1^, we sought to investigate the origins of this heterogeneity. To this end, we developed a scalable protocol for generating gastruloids that are monoclonal, *i.e.* derived from a single mouse embryonic stem cell (mESC). Single cell transcriptional profiling of monoclonal gastruloids revealed extensive inter-individual heterogeneity, with some hardly progressing, others resembling conventional gastruloids but biased towards mesodermal or neural lineages, and yet others bearing cell types rare or absent from conventional polyclonal gastruloids. To investigate this further, we leveraged DNA Typewriter^2^ to record the cell lineage relationships among the mESCs from which monoclonal gastruloids originate. Early in the expansion of “founder” mESCsーprior to induction of the resulting aggregates to form gastruloidsーwe observe clear examples of fate bias or fate restriction, *i.e.* sister clades that exhibit markedly different cell type compositions. In a separate experiment with DNA Typewriter, we reconstructed a monophyletic “tree of trees”, composed of ∼50,000 cells derived from ∼100 gastruloids, all descended from a single “founder of founders” stem cell. From these data, we find that founder mESCs that are more closely related are more likely to give rise to monoclonal gastruloids with similar cell type compositions. Our results suggest that fluctuations in the intrinsic states of mESCs are heritable, and shape their descendants’ fates across many cell divisions. Our study also showcases how DNA Typewriter can be used to reconstruct high-resolution, monophyletic cell lineage trees in stem cell models of early development.

## INTRODUCTION

Stem cell-derived models are powerful tools for studying cell fate decisions in early mammalian development and organogenesis, as they take otherwise inscrutable *in vivo* processes and render them accessible to both measurement and manipulation^3,4^. Conventionally, stem cell-derived models derive from a polyclonal aggregate of embryonic (ESC) or induced pluripotent (iPSC) stem cells^5,6^ subjected to specific regimens of external stimuli intended to mimic *in vivo* lineage specification. In recent decades, such protocols have been established to model the development of the brain^7,8^, gut^9^, retina^10^, kidney^11^, heart^12^ and other organs, as well as early structures such as the blastula^13^, gastrula^1,14–16^ or post-gastrulation embryo^17,18^ (<u>note</u>: as terminology is inconsistent across the literature, here we use “organoid” to refer to all stem cell-derived *in vitro* models, and “gastruloid” to refer to the specific mouse gastruloid protocol used here^14^).

Inter-organoid heterogeneity is a pervasive challenge in this field^19–21^, manifesting not only across experiments and labs, but also <u>within</u> batches of organoids induced under carefully controlled conditions. A potential explanation is that even among a seemingly uniform population of ESCs or iPSCs, stem cells’ internal states may fluctuate, leading to heterogeneous responses to identical stimuli during pre- or post-induction culturing. In this model, stem cell heterogeneity propagates to inter-aggregate heterogeneity, which in turn propagates to compositional and morphological heterogeneity among the resulting organoids. The possibility of internal state fluctuations in stem cells is consistent with the transcriptional heterogeneity revealed by single-cell RNA-seq (scRNA-seq) of ESCs or iPSCs^22,23^. Indeed, such fluctuations are hypothesized to contribute to the emergence of core-periphery differences in stem cell spheroids prior to induction^24,25^, and are not mutually exclusive with the possibility that cell-extrinsic factors (*e.g.* spheroid size, microenvironmental differences) also contribute.

If stem cell-intrinsic fluctuations contribute to inter-organoid heterogeneity, a question that follows is how heritable these fluctuations are. At one extreme, one can imagine these fluctuations occurring more quickly than the rate of cell division, such that there is no persistence or “memory” of them across stem cell divisions. In this scenario, the fate of any given stem cell aggregate, *i.e.* the morphology and cell type composition of the organoid that it will give rise to, would be stochastically determined <u>during or after</u> aggregation/induction. At the other extreme, one can imagine these fluctuations occurring slowly or infrequently, such that any given stem cell’s state is highly correlated with that of its ancestors and descendants. In this scenario, the intrinsic states of the cells composing a stem cell aggregate are inherited from their ancestors, such that the resulting organoid’s fate is effectively determined <u>prior to</u> aggregation/induction.

A challenge to experimentally distinguishing between these possibilities is that in most conventional organoid induction protocols, hundreds to thousands of stem cells are first aggregated, and the resulting polyclonal aggregate serves as starting material for the induction of an individual organoid. Thus, each organoid is founded by stem cells with potentially diverse internal states and unknown lineage relationships to one another, and these stem cells may furthermore contribute unequally to the organoid. As a consequence, any formulation of our question becomes entangled with additional complexities, *e.g.* how the fluctuating internal states of the unrelated or distantly related stem cells contributing to the polyclonal aggregate, whether heritable or not, interact to collectively shape the fate of the resulting organoid.

To simplify the question, we developed a general protocol for producing <u>monoclonal</u> stem cell aggregates^26^, such that the stem cells that give rise to each aggregate compose a “clone” whose members are as closely related as possible. We can induce these monoclonal aggregates to form conventional mouse gastruloids, which model symmetry breaking, axial elongation and germ layer specification^1^. Leveraging scRNA-seq, we find that monoclonal gastruloids exhibit extensive inter-individual heterogeneity. To probe the heritability of internal state fluctuations, we engineered the DNA Typewriter^2^ system into mESCs, and used it to measure the lineage relationships among cells and cell types in a series of monoclonal gastruloids. Early in the expansion of founder stem cells but prior to their induction to form monoclonal gastruloids, we observe evidence for fluctuations in the intrinsic states of mESCs that shape their descendants’ post-induction fates. Furthermore, within a small “family” of mESCs that subsequently serve as founders of ∼100 monoclonal gastruloids, we find that more closely related founder stem cells give rise to gastruloids that are correlated with respect to cell type composition. We conclude that the inter-individual heterogeneity across these gastruloids is shaped by internal cell state fluctuations in founder stem cells that are heritable across multiple cell divisions.

## RESULTS

### A protocol for generating monoclonal gastruloids

The process for generating monoclonal mouse gastruloids begins similarly to how one would generate a monoclonal, genetically engineered mESCs cell line (**Figure 1A-B**). We first dissociate mESCs into single cells and plate them onto a feeder layer of mouse embryonic fibroblasts (MEFs) at low density (200 to 600 cells per well of a 6-well tissue culture plate). Over several days on the MEF layer, each mESC expands into a dome-shaped colony with a sharp border. Five days after mESC seeding, the co-culture is treated with collagenase IV to gently dissociate the MEF layer while keeping mESC colonies intact. The mESC colonies are then collected and cultured for another day on a non-adherent plate, such that each colony gives rise to a spherical mESC aggregate. Each monoclonal stem cell aggregate (100-150 µm in diameter) is then transferred to an individual well of a 96-well culture plate containing NDiff 227 medium (equivalent to N2B27). Finally, we induce a mouse gastruloid in each well with 24-hour exposure to CHIR99021 (CHIR)^14^.

**Figure 1.**
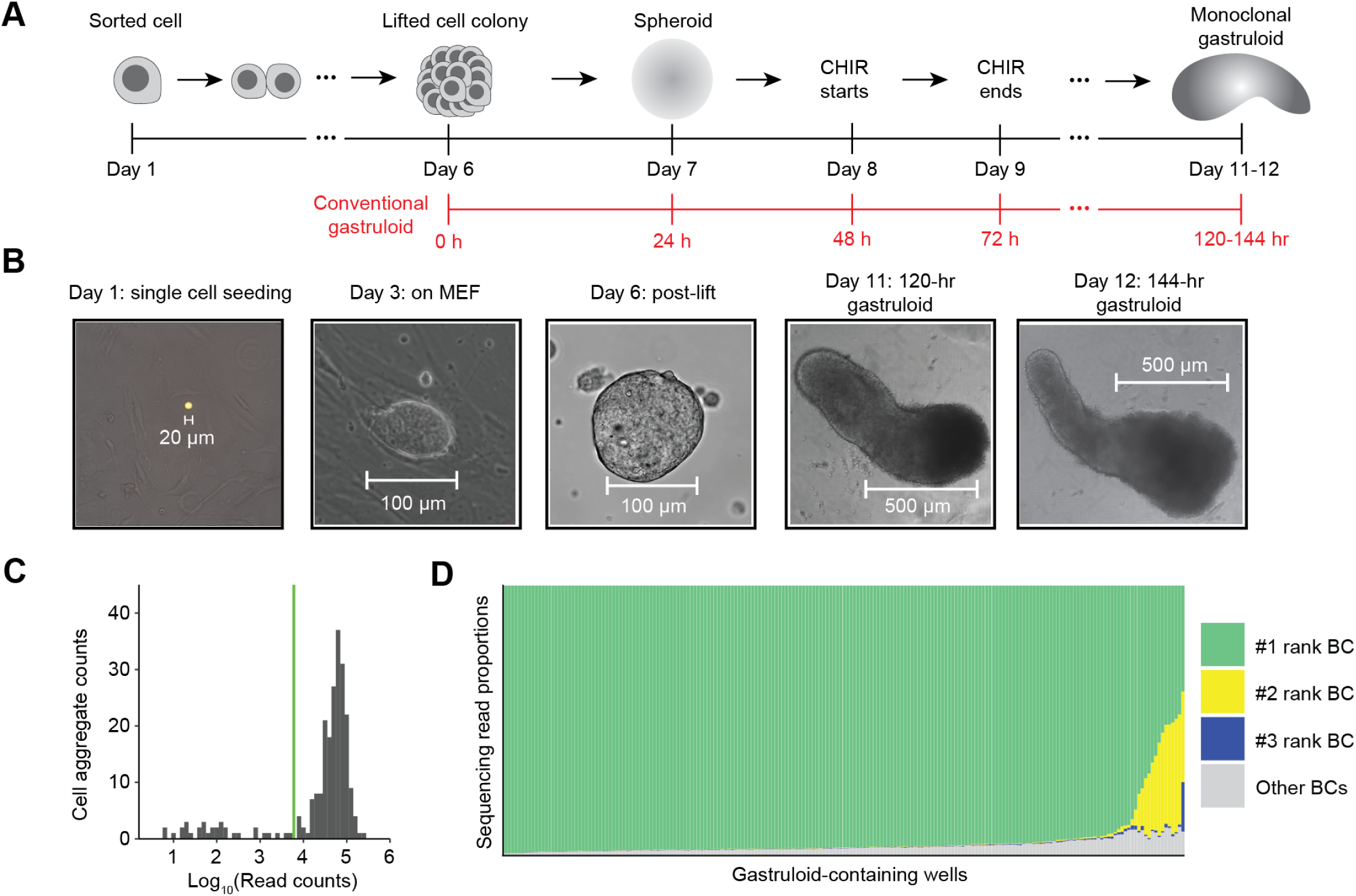
A protocol for generating monoclonal gastruloids. **(A)** Schematic of protocol for generating monoclonal mouse gastruloids. Dissociated mESCs are FACS-sorted followed by density controlled seeding onto a layer of MEF feeder cells. After single cells expand into monoclonal colonies, they are lifted off intact through collagenase IV treatment and gentle agitation, and transferred to a nonadherent plate where they give rise to spherical aggregates. The aggregates are then transferred again to an individual well where they undergo induction via 24-hr exposure to CHIR, a Wnt agonist. Timeline from the conventional mouse gastruloid protocol^14^ is displayed below in red text for a comparison. **(B)** Bright-field images of representative gastruloids from days 1, 3, 6, 11, and 12 are presented. For the day 1 image, GFP (green) and mCherry (red) fluorescence, also shown, were used to locate a GFP+/mCherry+ single cell on the MEF feeder cell layer. **(C)** Sequencing of static 8N DNA barcodes. Histogram of log-scaled read counts assigned to each of 234 wells. 201/234 wells were assigned ≥6,000 reads (green vertical line). **(D)** For each of the 201 wells with ≥6,000 assigned reads (x-axis), stacked bar plots of the proportions of reads corresponding to the top three most abundant DNA barcodes observed in that well (y-axis). In 93% of these wells, the top-ranked barcode accounted for ≥90% of the observed barcodes.

To formally test whether this protocol robustly gives rise to monoclonal gastruloids, we constructed a polyclonal mESC line bearing randomly integrated, static DNA barcodes. More specifically, static DNA barcodes (8N) were cloned to a puromycin and GFP-bearing lentiviral vector, and the resulting virus used to transduce mESCs at a low multiplicity of infection (MOI), such that the overwhelming majority of puromycin-selected mESCs should harbor only a single integrated DNA barcode. With these sequence-tagged mESCs, we continued the aforedescribed protocol to generate monoclonal mESC aggregates. Eight days after the initial plating of dissociated mESCs, we initiated a 24-hr pulse of 3 μM CHIR. Visualizing the aggregates 48 or 72 hrs after CHIR removal (which corresponds to the 120 or 144 hr time points in the conventional mouse gastruloid protocol), we observed elongating structures that resemble conventional polyclonal gastruloids (**Figure 1B**).

To query how many uniquely barcoded mESCs founded each gastruloid, we performed indexed PCR amplification and sequencing on genomic DNA from 234 individual wells to which aggregates had been transferred, with primers flanking the lentivirally integrated static DNA barcode. Amplicons were successfully sequenced from 201 of these wells (**Figure 1C**), of which 187 (93%) were dominated by a single barcode (≥90%). The result confirms that our protocol gives rise to mouse gastruloids that overwhelmingly derive from a single founder mESC. Of the 14 (7%) remaining wells, there were 13 for which the top two barcodes, and 1 for which the top three barcodes, accounted for ≥90% of sequences (**Figure 1D**). These instances could reflect di- or tri-clonal gastruloids (*i.e.* if two or three mESCs contributed to each aggregate that in turn gave rise to these gastruloids) or alternatively, monoclonal gastruloids whose founding mESC harbored 2-3 lentiviral integrants.

### Inter-individual heterogeneity among monoclonal gastruloids

We next sought to annotate cell types and characterize inter-individual heterogeneity among monoclonal gastruloids. To this end, we used piggyBac transposition to introduce two cassettes to mESCs, one bearing both DNA Typewriter Tape (DTT) and DTT-targeting enhanced prime editing guide RNAs (epegRNAs), and the other encoding a doxycycline-inducible prime editor (inducible PEmax or iPEmax) (**Figure S1**). Each cell in the resulting polyclonal mESC line was expected to harbor a unique complement of DTT/epegRNA integrants and at least 1 copy of iPEmax. Each DTT includes a static DNA barcode (12N; “TapeBC’’) upstream of the monomeric tape array that serves as substrate for ordered editing by DNA Typewriter. The TapeBC and DTT reside downstream of a T7 promoter sequence and constitutive Pol II promoter, providing two means of enhancing their recovery in scRNA-seq profiles. We note that for the experiment described in this section, the purpose of activating DNA Typewriter was not to record cell lineage, but rather to uniquely mark clones by enhancing barcode diversity.

Monoclonal gastruloids were generated by the protocol outlined above (**Figure 1A-B**), except that a 24-hr pulse of doxycycline prior to FACS and single cell seeding onto the MEF layer was used to induce PEmax expression, edit DTT and thereby boost barcode diversity. On day 12 (*i.e.* 72 hrs after the end of the 24-hr CHIR pulse), we collected individual, putatively monoclonal gastruloids from 144 wells. We pooled these gastruloids, dissociated them to single nuclei, and performed three-level single cell combinatorial indexing RNA sequencing (sci-RNA-seq3)^27^ (**Figure 2A**). A modified version of this protocol was used in which *in situ* T7 transcription, together with enrichment PCR, boosts recovery of DTT^28^. In parallel, in order to facilitate “gastruloid-to-well” assignments in our later analyses, we also isolated genomic DNA from cellular debris present in residual media from each well, followed by PCR amplification and sequencing of DTT (“Debris-seq”; **Figure 2A**).

**Figure 2.**
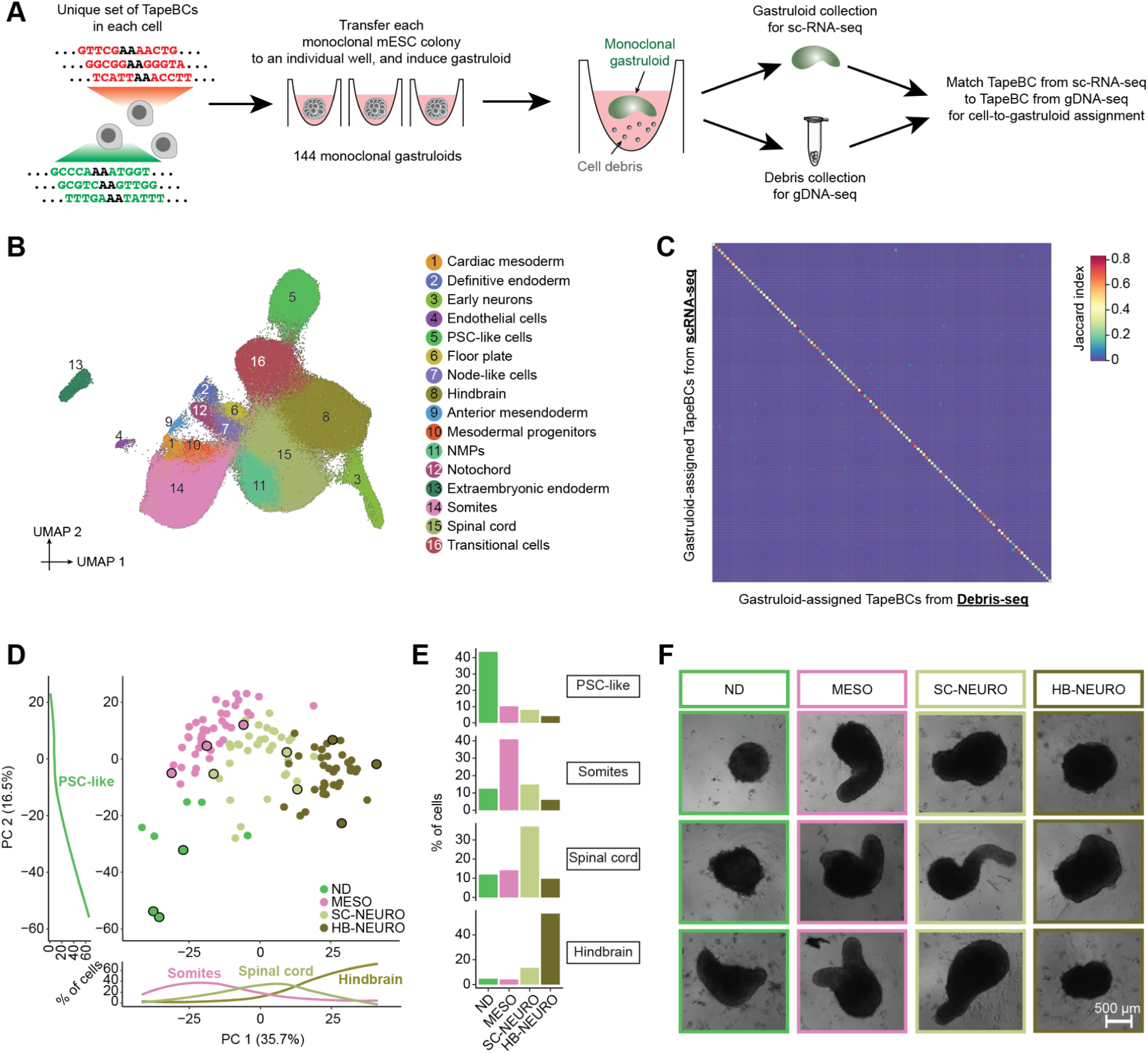
Quantifying phenotypic heterogeneity among mouse gastruloids derived from single cells. **(A)** Schematic of mapping cells to wells/gastruloids by intersecting TapeBCs recovered from sci-RNA-seq or Debris-seq. **(B)** 2D UMAP visualization of 247,064 cells derived from 144 mouse gastruloids. Colors and numbers correspond to 16 cell cluster annotations as listed on the right (**Table S1**). **(C)** After identifying the 121 gastruloids, Jaccard similarities were computed between the TapeBCs of cells assigned to each gastruloid (row; detected in ≥5% of the cells within that gastruloid) and those detected by Debris-seq of each gastruloid’s well (column). **(D)** Embeddings of pseudobulk RNA-seq profiles of 121 mouse gastruloids in PCA space with visualization of top two PCs. Briefly, single nucleus data from each gastruloid was aggregated to create 121 profiles, on which we performed dimensionality reduction via PCA. Gastruloids were clustered into four groups based on their cell type compositions using k-means clustering (ND: non-differentiating, MESO: somite-like, SC-NEURO: spinal cord-like, or HB-NEURO: hindbrain-like). The percentage of cells belonging to selected cell types is plotted along PC1 (bottom) and PC2 (right). The plotted line was generated using the geom_smooth function in ggplot2. The dots highlighted with black circles correspond to the gastruloids whose images are shown in panel **F**. **(E)** The proportions of cells in four selected cell types for individual gastruloids were calculated, and the average values within each of the four clusters are presented. **(F)** Representative images of gastruloids from each of the four groups are presented.

We performed cell type annotation on all scRNA-seq data, *i.e.* without parsing out the cells composing individual gastruloids. Following demultiplexing, alignment, and exclusion of low-quality cells and potential doublets, we obtained profiles for 247,064 cells (median 609 UMIs per cell). We then performed dimensionality reduction, clustering and manual annotation based on gene markers and integrative analyses with other *in vitro* and *in vivo* datasets^25,29–33^ (**Figure 2B**; **Figures S2-5**; **Table S1**). Reassuringly, many of the cell types observed in 144-hr monoclonal mouse gastruloids aligned with expectation as defined by four *in vitro* studies^25,29,32,33^ that profiled 120-hr conventional polyclonal mouse gastruloids with scRNA-seq, *e.g.* cell types resembling neuromesodermal progenitors (NMPs; *T+*, *Cdx2*+), spinal cord (*Hoxb4+*; *Hoxc6*+), mesodermal progenitors (*Tbx6*+, *Hes7*+), somites (*Tcf15*+, *Meox1*+), cardiac mesoderm (*Hand1+*; *Tbx20*+), definitive endoderm (*Sox17+*, *Trh*+), and endothelial cells (*Kdr*+, *Cdh5+*).

However, in contrast with 120-hr conventional polyclonal mouse gastruloids, these 144-hr monoclonal mouse gastruloids also contained an abundance of hindbrain-like (*Egr2*+) cells, a subset of which expressed midbrain-hindbrain boundary markers (*En1+*, *En2+*, *Pax5*+), as well as cells resembling early differentiating neurons (*Neurod1+*, *Ebf2+*, *Ebf3*+) that were transcriptionally contiguous with both spinal cord and hindbrain-like cells (**Figure 2B**; **Figures S2-5**; **Table S1**). Additional abundant cell types included undifferentiated PSC-like cells, expressing both epiblast (*Nanog+*, *Utf1+*) and primordial germ cell (*Dppa3*+; *Dppa5a*+) markers, and “transitional” cells (*Bhlhe41*+; *Atp6v0b*+), seemingly intermediate between PSCs and germ layers but failing to express primitive streak markers. Rarer, unexpected cell types resembled the node (*Foxa2*+; *Kcnip4*+), notochord (*T+, Noto+*), floor plate (*Shh+*; *Foxa2*+), anterior mesendoderm (*Eomes*+; *Lhx1*+), and extraembryonic endoderm (*Ttr4*+, *Gata4*+, *Sparc*+). Of note, some of these have correlates in a subset of the conventional gastruloid studies^25,29,32,33^ but at lower proportions (*e.g.* neurons in Rosen *et al.* (2022)^32^), while others are entirely absent (*e.g.* extraembryonic endoderm).

Given that there are also clear differences between the conventional gastruloid studies (**Figure S3**; *e.g.* the paucity of neural lineage cells in Suppinger *et al.* (2023)^25^), there are trivial but potentially valid explanations for the differences we observe between 144-hr monoclonal vs. 120-hr polyclonal mouse gastruloids, *e.g.* subtle protocol differences between studies conducted by different groups, 144 vs. 120-hr collection, etc. However, an additional possibility is that “outlier” states within a conventional polyclonal aggregate are non-autonomously “buffered” by other clones, and thus unlikely to dominate the fate of a given gastruloid. In contrast, with the monoclonal gastruloid protocol, the cells composing the stem cell aggregate are highly related, and free from such buffering might be more likely to support the emergence of unexpected cell types or cell type compositions.

As gastruloids’ cell type compositions might reveal some of this latent heterogeneity, we sought to assign each scRNA-seq profile to one of the 144 wells in which gastruloids were grown (**Figure 2A**), with the following heuristic: 1) We defined a white list of 4,995 unique, well-specific DTTs observed by Debris-seq, and aligned DTTs observed by scRNA-seq to this white list. After excluding cells with <3 DTTs detected, we obtained a matrix of 4,609 DTTs-by-154,988 cells. 3) We quantified overlap between DTTs seen in cells vs. wells, and assigned 129,853 cells for whom the largest overlap exceeded the second largest overlap by >50%. 4) For additional assignments, we performed dimensionality reduction (PCA) of scRNA-seq profiles based solely on the DTT-by-cell matrix (**Figure S6A**). For still-unassigned cells, we identified the wells of their top 10 nearest already-assigned neighbor cells in PCA space. We then assigned 19,363 cells for whom the largest overlap exceeded the second largest overlap by >50%. 5) We excluded 23 wells with no unique DTTs or <100 assigned cells. Cells assigned the remaining 121 wells were assumed to derive from those wells’ gastruloids.

Altogether, 148,724 cells were assigned to 121 wells/gastruloids (1,229 ± 953). Focusing only on DTTs detected in ≥5% of cells assigned to a given gastruloid, there were 25 ± 16 DTT/epegRNA integrants per founder mESCs (range 5-93). For quality control, we assessed the Jaccard similarity between DTTs observed in each of the 121 cell groups (by scRNA-seq) vs. those observed in each of the 121 wells (by Debris-seq), which confirmed a strong one-to-one coherence (**Figure 2C**).

To assess inter-individual heterogeneity, we performed PCA on “pseudo-bulked” transcriptional profiles of individual monoclonal gastruloids. PC1 (35.7% of variance) separated gastruloids dominated by somite-like, spinal cord-like or hindbrain-like cells, while PC2 (16.5% of variance) correlated with the proportion of cells remaining in an undifferentiated PSC-like state (**Figure 2D**). Based on their cell type compositions, we clustered the 121 gastruloids into four groups, which were dominated by PSC-like (non-differentiating or ND), somite-like (MESO), spinal cord-like (SC-NEURO), or hindbrain-like (HB-NEURO) cells (**Figure 2E**; **Figure S6B**). Many other cell types were strongly associated with specific groups, *e.g.* mesodermal cell types such as anterior mesendoderm with MESO gastruloids, and early neurons with HB-NEURO gastruloids. Furthermore, some cell types were overwhelmingly accounted for by a handful of monoclonal gastruloids (**Figure S6C**). For example, just 3 of the 121 gastruloids accounted for 59% of cells annotated as anterior mesoderm, 45% of cells annotated as mesodermal progenitors, and 46% of cells annotated as cardiac mesoderm. Of note, recovery, which presumably correlates with the total number of cells in each monoclonal gastruloid, did not significantly differ across the four groups (**Figure S6D**). However, from images of each well prior to harvesting, composition-based group identity did correlate with gastruloids’ morphological appearances, with MESO and SC-NEURO gastruloids more likely to exhibit elongation (**Figure 2F**).

### Phylogenetic reconstruction of the cell lineage histories of monoclonal gastruloids

With a protocol for generating monoclonal gastruloids in hand, together with results suggesting that their heterogeneity is at least on par with, and may even exceed, that of polyclonal gastruloids, we sought to establish a framework for reconstructing their cell lineage histories. For this, we once again relied on mESCs equipped with DNA Typewriter^2^ (**Figure S1**), but first established three monoclonal cell lines (Clone-05, Clone-25, Clone-32). Based on amplicon sequencing of TapeBC barcodes from genomic DNA, these lines respectively bear at least 76, 116 and 84 DTT/epegRNA integrants (lower bounds, as some integrants are duplicated by piggyBac excision and re-integration events^28,34^). Thus, with six ordered units per DTT, each line harbors at least 400 sites that could potentially be written to by DNA Typewriter (**Figure S7**).

We mixed these cell lines and then generated monoclonal gastruloids as outlined above, the main differences being: 1) the inclusion of 100 ng/mL doxycycline beginning 24 hrs prior to sorting and seeding of single cells and continuing throughout, in order to induce PEmax; and 2) the collection of replaced media on Day 10, for Debris-seq (**Figure 3A**). Rapid analysis of Debris-seq data allowed us to identify and focus on harvesting wells containing gastruloids with high rates of editing at DTT sites. On Day 11, which corresponds to the 120-hr time point of the conventional protocol, we selected 8 clonal gastruloids for their elongated morphology and high DTT editing rates, harvested them, and performed droplet-based scRNA-seq on the 10X Genomics platform with PCR-based feature enrichment to enhance recovery of DTT.

**Figure 3.**
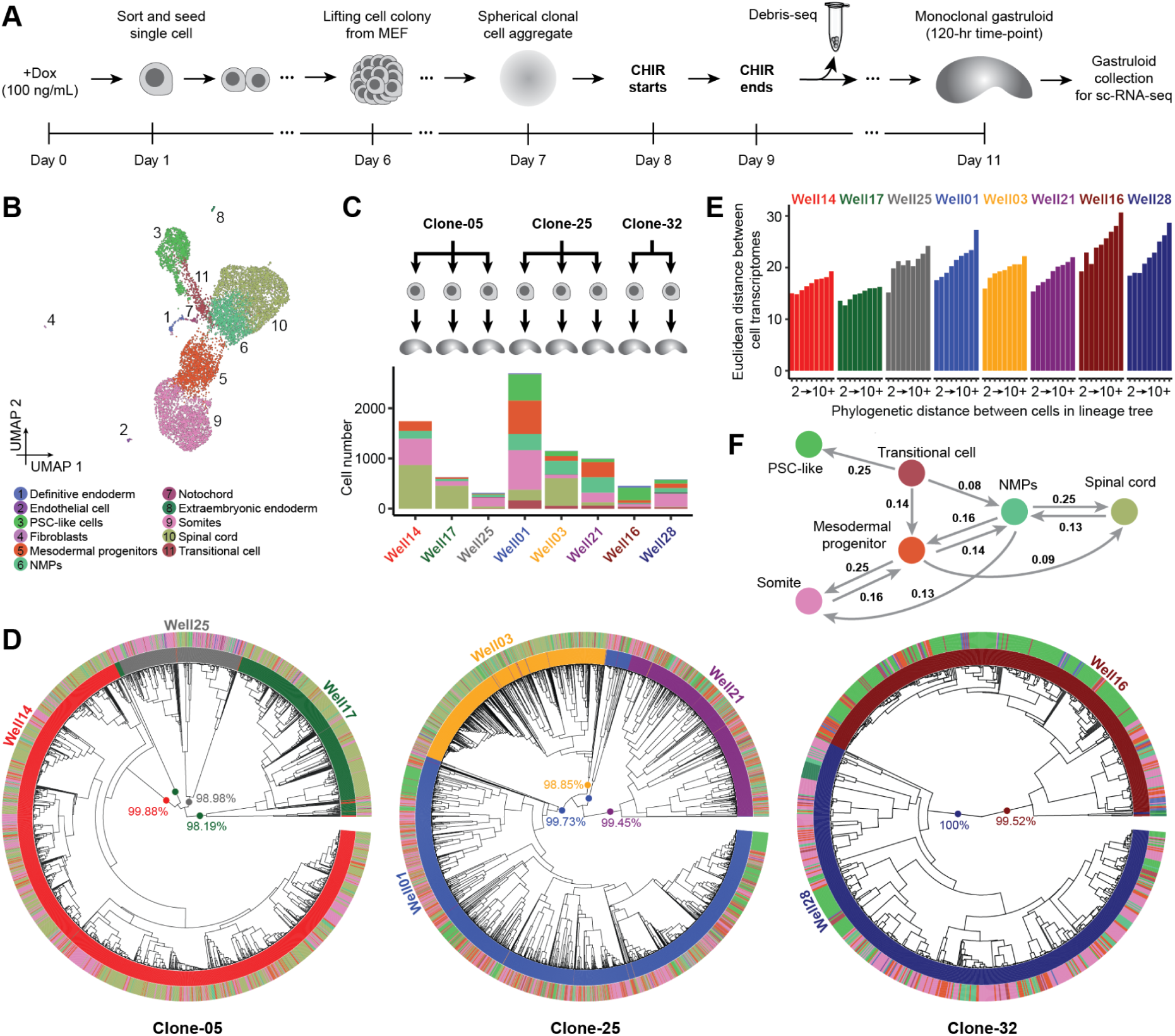
DNA Typewriter-based reconstruction of the cell lineage histories of eight monoclonal gastruloids. **(A)** Summary of experiment performed on mESCs engineered to bear all DNA Typewriter components (Dox-inducible PEmax, epegRNAs, DTT). Dox induction of PEmax was initiated 24 hrs prior to seeding single-cell dissociated mESCs onto the MEF layer. Eight wells were selected based on Debris-seq results and morphology, and these gastruloids harvested for scRNA-seq. **(B)** 2D UMAP visualization of the resulting scRNA-seq data for 9,929 cells, annotated by cell type. **(C)** Through comparison of TapeBCs detected by Debris-seq vs. scRNA-seq, 8,545 cells profiled by scRNA-seq could be assigned to a well-of-origin. **(D)** Phylogenetic reconstruction of cell lineage relationships of 2,438 (Clone-05), 4,418 (Clone-25), and 960 (Clone-32) higher-quality, gastruloid-assigned cells per clone, constructed by first splitting some TapeBCs to account for piggyBac excision and re-integration events, and then applying UPGMA with a custom DTT-based distance metric. The inner circle shows the phylogenetic tree based on cell-cell distances, the colors of the middle circle correspond to 8 gastruloids to which each cell was assigned, and the colors of the outer circle correspond to the 11 annotated cell types. Key for gastruloid colors is in panel **C**. Key for cell type colors is in panel **B**. The center of each tree highlights ten major clades, labeled by the % of its cells assigned to a single gastruloid. **(E)** For cells assigned to each gastruloid, the mean Euclidean distances between all possible pairs of cells, grouped into bins by phylogenetic distances (calculated from a tree reconstructed using cells from that gastruloid) ranging from 2 to 10+, are shown. Euclidean distances were calculated following dimensionality reduction (30 principal components) of the cell × gene matrix. **(F)** A directional graph of the inferred lineage relationships, with inferred transition probabilities >0.08, among the six most abundant cell types (collectively >98% of cells), based on PATH^37^.

Following conventional filtering, we obtained scRNA-seq profiles for 9,929 cells (median UMIs detected: 10,881; median genes detected: 3,611; **Figure 3B**; **Figure S8A**). By matching the DTT sequences associated with each single cell transcriptome to those observed in Debris-seq, we could confidently determine the “well of origin” of 86% of cells (8,545 of 9,929), *i.e.* assign them to one of the eight monoclonal gastruloids. The numbers of scRNA-seq profiles recovered per gastruloid ranged from 312 to 2,696 (mean 1,068) (**Figure 3C**; **Figure S8B-C**).

Once again, we observed a continuous range of cell states that includes PSC-like cells differentiating and then bifurcating to both mesodermal and neural states of varying maturity (**Figure 3B**). Following procedures described above, we annotated eleven cell types, but six of these accounted for >98% of cells (PSC-like, transitional, NMPs, mesodermal progenitors, somites, spinal cord) (**Figure 3C**; **Figure S8D**; **Table S1**). Furthermore, presumably due to time point, technical and/or sampling differences (**Figure 2A**: [144 x 144-hr gastruloids; sci-RNA-seq3; 247,064 nuclei; median UMI 609] vs. **Figure 3A**: [8 x 120-hr gastruloids; 10X Genomics; 9,929 cells; median UMI 10,881]), some previously observed cell types were missing among these eight gastruloids (cardiac mesoderm, early neurons, floor plate, node-like cells, hindbrain, anterior mesoderm), while one was only detected here (fibroblasts).

For reconstructing cell lineage trees, we focused on 7,816 higher-quality, gastruloid-assigned cells (Clone-05: 2,438; Clone-25: 4,418; Clone-32: 960; **Figure S8E**). For these cells, we recovered 45 ± 13 DTTs bearing at least one edit as part of their scRNA-seq profiles (3.0 ± 1.5 edits per edited DTT; mean 138 ± 47 edits per cell). Interestingly, the cumulative activity of DNA Typewriter modestly varied between cells from different gastruloids, even those derived from the same clone, but was relatively consistent within cell types of an individual gastruloid (**Figure S9**).

We constructed phylogenetic trees for each clone by first splitting some TapeBCs to account for piggyBac excision and re-integration events (**Figure S10; Supplementary Note**) and then applying UPGMA (Unweighted Pair Group Method with Arithmetic Mean) with a custom DTT-based distance metric (**Figure 3D**; **Figures S11-13**). A bootstrapping analysis, based on 100 iterations of DTT sampling with replacement, found that 48% (Clone-05), 77%, (Clone-25) and 60% (Clone-32) of nodes had moderate to strong support (transfer bootstrap expectation (TBE) >70%)^35^. Poorly supported nodes were overwhelmingly shallow, while deep nodes were strongly supported (**Figure S8F**). The deepest nodes corresponded to 10 major clades, each of whose membership overwhelmingly (∼99%) derived from just one of the eight monoclonal gastruloids (**Figure 3D**; **Figure S8G**). These clades were defined by numerous edits always or nearly always present when a given DTT was sampled from a given gastruloid, but hardly ever found in any other gastruloid (**Figure S14**). We infer that these “gastruloid-defining edits” are likely to have occurred in the 24 hrs following the induction of PEmax but prior to the seeding of single founder cells.

What is the relationship between cell lineage and cell state? In each monoclonal gastruloid, we observe a positive correlation between the phylogenetic distances vs. “state distances” of pairs of cells^36^, with the latter defined here as Euclidean distance in a reduced dimensionality transformation of the cell × gene matrix (**Figure 3E**). To investigate this further, we sought to infer the lineage relationships among observed cell types from the phylogenetic tree. For this, we calculated two metrics for each possible pair of cell types (including self-pairs): 1) the normalized, log-transformed frequency at which the cell types appear in pairs of cells that are nearest neighbors in the terminal leaves of the phylogenetic tree (**Figure S15A-B**); and 2) phylogenetic cross-correlation with the PATH package^37^ (**Figure S15C-D**). Focusing on the six abundant cell types (collectively >98% of cells), a directional graph derived from PATH-based state-transition probabilities was broadly consistent with expectation, except for the directionality of the transition between PSC-like cells and transitional cells (**Figure 3F**). We suspect that this may be because the PSC-like cells that have persisted through to harvesting at 120-hrs are self-renewing and refractory to differentiation signals.

### Strong fate biases within a few cell divisions of monoclonal stem cell seeding

As with the 144 x 144-hr gastruloids, our 8 x 120-hr gastruloids exhibit substantial heterogeneity with respect to cell type composition, but our examples now include multiple gastruloids derived from the same clone. Most strikingly, the monoclonal gastruloids originating from Clone-05 include one that is well-balanced (Well14), one that is neurally biased (Well17) and one that is mesodermally biased (Well25) (**Figure S16**). However, there is also clear structure <u>within</u> the phylogenetic trees of individual gastruloids, evident in the trees themselves (**Figure 4A**) and quantified by the extent of phylogenetic auto-correlation of individual cell types (**Figure S15C**).

**Figure 4.**
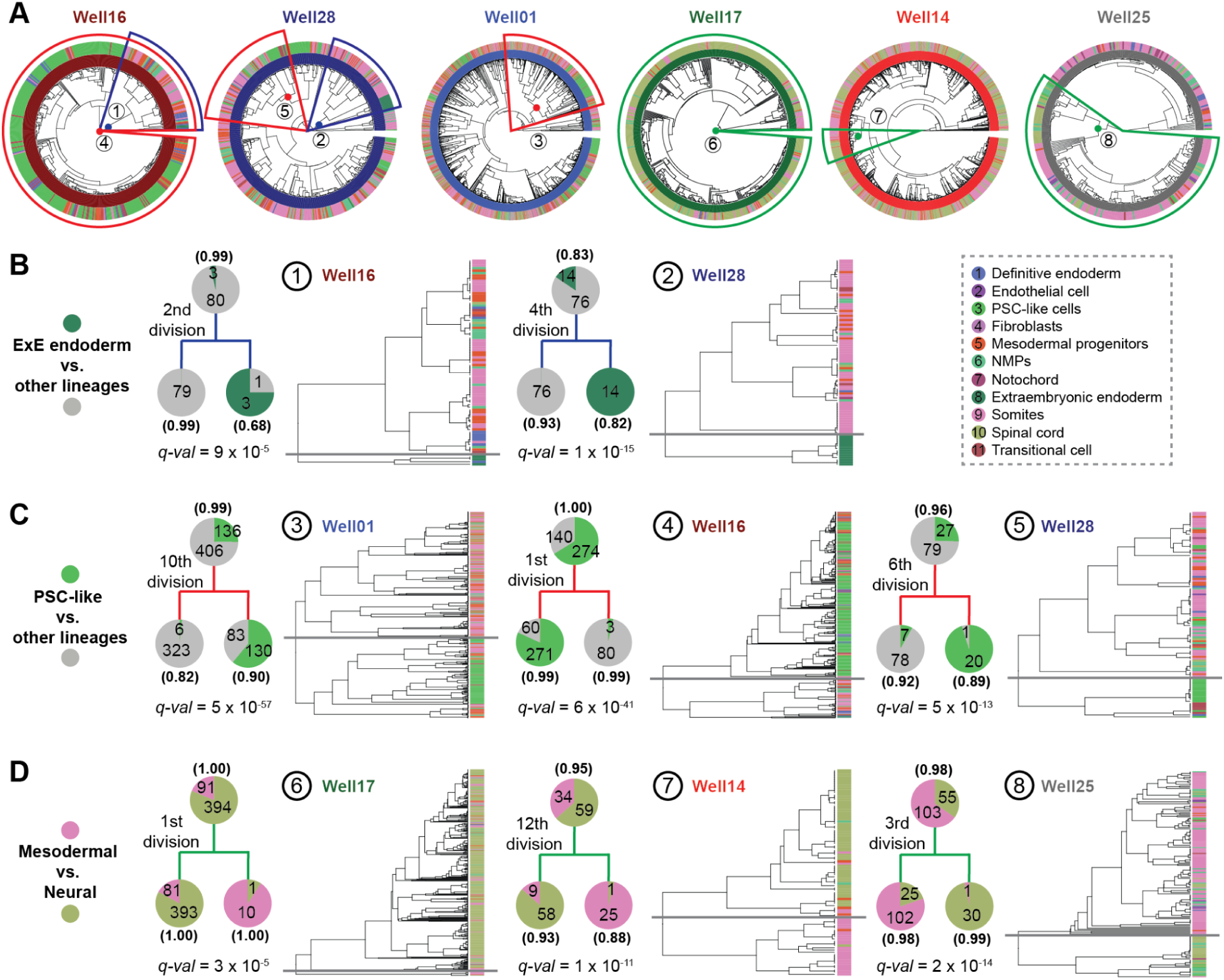
Cell lineage histories of individual monoclonal gastruloids suggest fate biases and fate restrictions in embryonic stem cells prior to induction. **(A)** For each monoclonal gastruloid, phylogenetic reconstruction of cell lineage relationships among higher-quality, gastruloid-assigned cells. The inner circle shows the phylogenetic tree based on cell-cell distances, the colors of the middle circle correspond to the gastruloids to which each cell was assigned, and the colors of the outer circle correspond to the 11 annotated cell types. We manually selected eight clades to highlight, corresponding to inferred cell type divisions for which daughter clades appear to be fate-restricted or fate-biased in specific ways, including: extraembryonic endoderm vs. other (green, detailed in panel **B**), undifferentiated vs. differentiated (blue, detailed in panel **C**), and mesodermal vs. neural (red, detailed in panel **D**). **(B)** Two inferred cell divisions whose daughter clades appear to be differentially fate-restricted with respect to extraembryonic endoderm (dark green) vs. other (gray) cell types. **(C)** Three inferred cell divisions whose daughter clades appear to be differentially fate-biased with respect to undifferentiated (PSC-like & translational; olive green) vs. differentiated (all other; gray) cell types. **(D)** Three inferred cell divisions whose daughter clades appear to be differentially fate-biased with respect to mesodermal (somites & mesodermal progenitors; pink) vs. neural (spinal cord & NMPs; light yellow) cell types. In each subpanel of panels **B-D**, pie charts at the ancestral node and its two child nodes indicate cell numbers by category. The number of inferred cell divisions from the root to the highlighted division is indicated to the left. Transfer bootstrap expectation (TBE) values^35^, shown in bold/parentheses for the ancestral and child nodes, were calculated based on 100 bootstraps. Fisher’s exact test was used to compare categorical counts between child nodes, with *p*-values corrected using Benjamini-Hochberg (FDR) method across all ancestor nodes within the same gastruloid tree to obtain *q-*values. A subtree view of the clade is shown on the right, with tip cells colored by cell type. The gray horizontal line marks the highlighted cell division.

The strongest auto-correlation is observed for extraembryonic endoderm, a cell type not typically observed in monoclonal gastruloids (**Figure S5**). Remarkably, 94% (17/18) of cells bearing this annotation appear in just two homogenous clades, both from Clone-32 gastruloids, predicted to have derived from the second (Well16 gastruloid) or 4th (Well28 gastruloid) cell division (**Figure 4B**; **Figure 3C**). A plausible explanation is that during early clonal expansion from a single “founder” cell to a colony, but prior to dissociation from MEF feeder-cell layer, some mESCs spontaneously differentiated into extraembryonic endoderm^38^. These extraembryonic endoderm cells may have remained associated with other mESCs throughout the spherical aggregation and CHIR-treatment of these gastruloids.

This phenomenon of an internal, strongly supported node in a gastruloid’s phylogenetic tree, whose daughter clades exhibit markedly different distributions of cell type compositions (**Figure 4B**), was observed for more common cell types as well (**Figure 4C-D**). For example, there are clear examples of inferred ancestral cells for whom one daughter clade is dominated by undifferentiated (PSC-like, transitional) and the other by differentiated (mesodermal, neural lineages) cell types (**Figure 4C**). There are also clear examples of inferred ancestral cells for whom one daughter clade is dominated by mesodermal and the other by neural cell types (**Figure 4D**). We note that all aspects of the phylogenetic reconstruction algorithm are “blind” to cell type, and these distributions are highly unlikely to have occurred by chance (**Figure 4B-D**).

Are these fate restrictions/biases occurring before or after induction of the gastruloid? Of note, for each class of fate restriction/bias highlighted above and in **Figure 4**, at least one example occurs within a few cell divisions of the monoclonal gastruloid’s founding. For example, example #2 highlights the 4th cell division of the Well28 gastruloid, with one daughter clade composed exclusively of extraembryonic endoderm (14/14) and the other exclusively of other cell types (76/76) (**Figure 4B**; *q*-value = 1e-14). Example #4 highlights the 1st cell division of the Well16 gastruloid, with one daughter clade composed largely of PSC-like cells (271/331) and the other daughter clade overwhelmingly of differentiated cell types (80/83) (**Figure 4C**; *q*-value = 6e-41). Example #8 highlights the 3rd cell division of the Well25 gastruloid, with one daughter clade composed largely of mesodermal cell types (102/127) and the other almost entirely of neural cell types (30/31) (**Figure 4D**; *q*-value = 2e-14). As each founder cell was expanded to an aggregate of 400-700 mESCs over seven days prior to lifting, the cell divisions giving rise to these large clades are overwhelmingly likely to have occurred long prior to CHIR-mediated induction.

### The cell type compositions of monoclonal gastruloids are informed by relatedness of their founders

To summarize the last section, our analyses of the phylogenetic histories of eight monoclonal gastruloids identified numerous examples of well-supported “sister clades” whose fate biases were highly divergent from one another. Many of these examples appeared to have occurred within the first few cell divisions of the seeding of a founding mESC. Heritable fluctuations in the intrinsic states of these stem cells, *e.g.* in one daughter cell but not the other, are a plausible explanation for these strong fate biases, but there are other possibilities. For example, the clades defined by a pair of differentially fate-biased daughter cells might have different spatial distributions in the resulting stem cell spheroid, and these spatial distributions might in turn underlie the observed fate biases^25,39^.

As a more rigorous test of our hypothesis, we designed a two-epoch “tree-of-trees” experiment (**Figure 5A**). In the first epoch of this design, we seeded a single mESC to generate a colony of 500-1000 cells. In the second epoch, we isolated this monoclonal colony, dissociated it to single mESCs, and then followed the monoclonal gastruloid induction protocol. A key point is that we induce the activity of DNA Typewriter at the onset of the first epoch, such that we are recording the lineage relationships among founder mESCs, each of which will go on to seed an individual monoclonal gastruloid. We can then ask whether or not the founder mESCs that are more closely related (during the first epoch) tend to give rise to compositionally similar gastruloids (during the second epoch). Additional features of this experimental design include: 1) it constrains the lineage distances between founder mESCs; and 2) it tests our hypothesis on mESCs whose lineage relationships are not confounded by spatial correlation.

**Figure 5.**
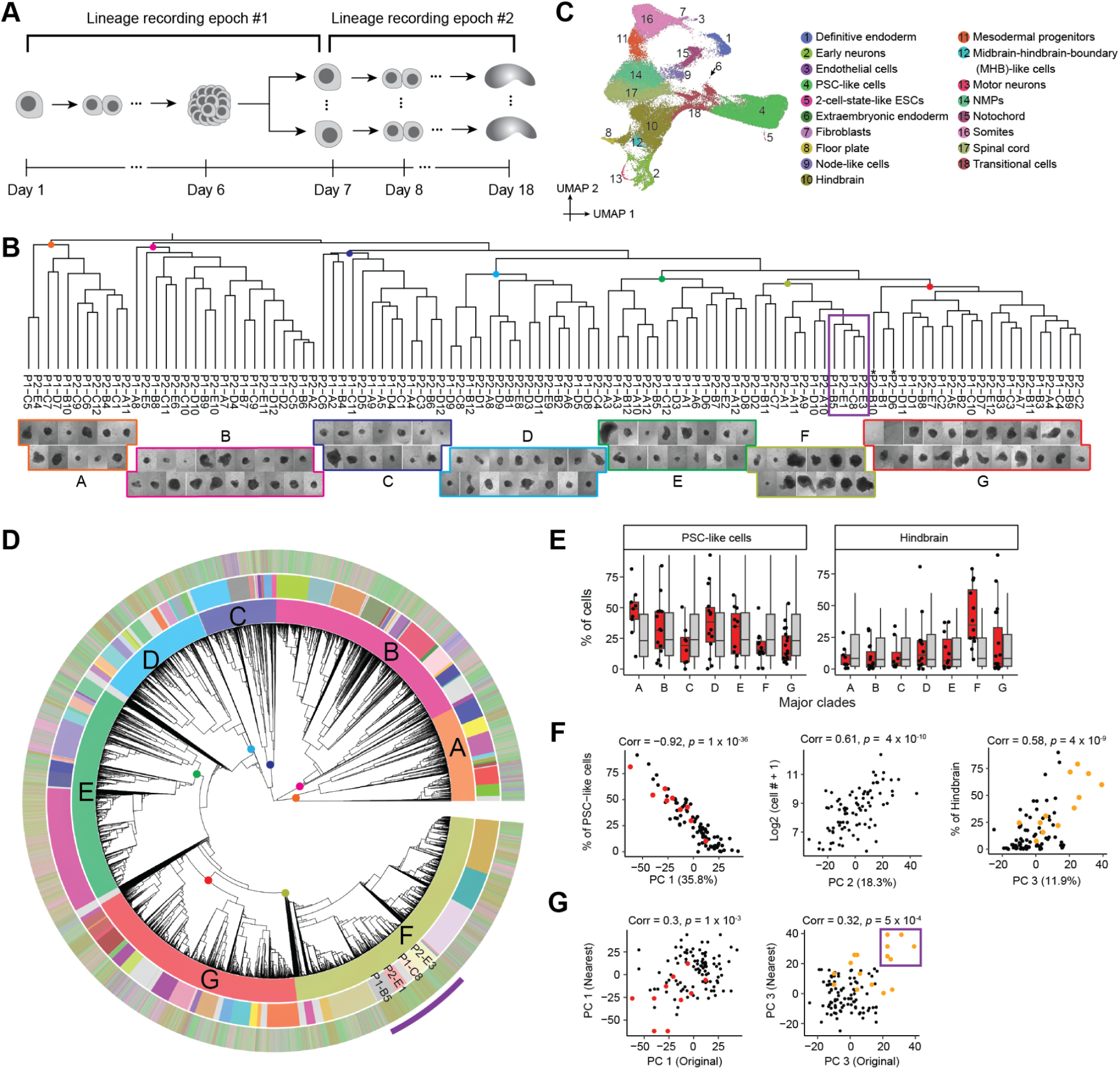
The cell type compositions of monoclonal gastruloids are informed by relatedness of their founders. **(A)** A two-epoch experiment was performed to record the lineage relationships among founder mESCs of monoclonal gastruloids. In the first epoch, a single “founder of founders” mESC was expanded to a colony of 500-1000 cells, which was then isolated, dissociated to single mESCs, and replated per the monoclonal gastruloid induction protocol. During the second epoch, 108 monoclonal cell aggregates were transferred to individual wells and induced into form gastruloids, which were harvested at Day 18 (equivalent of 144-hours in the conventional protocol). Two gastruloids (P2-D6 & P2-B10, names marked with asterisks, both in clade G) were lost during transfer. **(B)** A phylogenetic tree of 108 gastruloids was constructed based on gastruloid-gastruloid distances derived from monoclonal gastruloid-specific edits of DTTs captured by Debris-seq, with representative images of individual gastruloids shown below. To calculate the phylogenetic distance between two gastruloids, the following steps were performed: For each gastruloid, at each of the six sites across all 76 DTTs, the percentage of reads for each specific edit was determined, and combinations with at least 20% of reads were identified as dominant "TapeBC-Site-Edit" combinations. The Jaccard similarity between the two gastruloids’ sets of dominant combinations was calculated, with (1 - similarity) representing their distance. The images at the bottom for individual gastruloids correspond to seven major clades (A-F). Four clade F gastruloids which have a high percentage of hindbrain cells are highlighted by a purple rectangle (see text and purple arc in panel **D**). **(C)** 2D UMAP visualization of the resulting scRNA-seq data for 66,978 cells, annotated by cell type. **(D)** Phylogenetic reconstruction of cell lineage relationships of 58,283 higher-quality cells, constructed by first splitting some TapeBCs to account for piggyBac excision and re-integration events, and then applying UPGMA with a custom DTT-based distance metric. From inner to outer circle: the first circle shows the phylogenetic tree based on cell-cell distances, the colors of the second circle correspond to seven major clades to which each cell was assigned (with grey representing 702 unassigned cells), the colors of the third circle correspond to 96 wells/gastruloids to which each cell was assigned (with grey representing 6,320 unassigned cells), and the colors of the fourth circle correspond to the 18 annotated cell types. Key for cell type colors is in panel **C**. Four subclades within clade F, assigned to gastruloids with a high percentage of hindbrain cells, are labeled and highlighted with a purple arc at the periphery (see text and purple rectangle in panel **B**). **(E)** The proportions of the PSC-like cells (left) and hindbrain cells (right) for 88 individual gastruloids (≥50 cells assigned) were calculated, and are shown as box plots (red) for clades A-G. The group labels for each gastruloid were permuted 5,000 times, and cell-type proportions were recalculated each time. Boxplots, showing 88 replicates from real observations (red) and 5,000 × 88 replicates from permutation (grey) in each subpanel, represent the interquartile range (25th, 50th, 75th percentiles), with whiskers extending to 1.5× the IQR. Black dots represent replicates from the real observations. **(F)** Embeddings of pseudo-bulk RNA-seq profiles of 88 monoclonal gastruloids (≥50 cells assigned) were generated by aggregating single-nucleus data and performing PCA for dimensionality reduction. The percentage of PSC-like cells is plotted along PC1 (left), the log2-transformed cell number along PC2 (middle), and the percentage of hindbrain cells along PC3 (right). Pearson correlation coefficients and p-values are indicated above each plot. Gastruloids from major clade A (red) and clade F (orange) are colored. **(G)** For each of the 88 gastruloids, their nearest neighbors were identified from the phylogenetic lineage tree, resulting in 112 pairs. Of note, some gastruloids had more than one nearest neighbor, and some pairs were redundant (in which case were retained). For two of the top three PCs, PC1 and PC3, the values for each gastruloid (x-axis) are plotted against those of their nearest neighbors (y-axis). Pearson correlation coefficients and p-values are indicated above each plot. Gastruloids from major clade A (red) and clade F (orange) are colored. Four gastruloids in clade F, forming six pairs and exhibiting a high percentage of hindbrain cells as shown in panels **B** & **D**, are highlighted within a purple rectangle.

To implement this design, we seeded a single mESC derived from Clone-05 to a well, harvested the resulting “first epoch” aggregate, dissociated and plated the resulting mESCs, and then picked and seeded 108 of the resulting “second phase” aggregates to individual wells, where they were induced to form monoclonal gastruloids (**Figure 5A**). Doxycycline was present throughout in order to induce PEmax expression and DNA Typewriter activity during both epochs. We harvested the gastruloids at Day 18, which corresponds to the 144-hr time point in the conventional gastruloid induction protocol, concurrently collecting media from each well for Debris-seq. We pooled all gastruloids for scRNA-seq, collecting both transcriptomes and DTT, the latter via enrichment PCR.

A phylogenetic tree of the relationships of monoclonal gastruloids to one another, *i.e.* reflecting the relationships among mESCs during the first phase, was constructed using a distance metric based on Jaccard similarity between dominant edits observed by Debris-seq of each well. This resulting tree (**Figure 5B**, top) was robust to bootstrapping analysis, with 99% (106/107) ancestral nodes showing moderate to strong support (TBE>70%) (**Figure S17A**). Each gastruloid was assigned to one of seven major clades (labeled A-G in **Figure 5** and **Figures S17-19**), some of which subjectively exhibited different morphological characteristics.

From scRNA-seq, we recovered transcriptional profiles for 66,978 cells which we annotated to 18 cell types (**Figure 5C**). These annotations largely overlapped with our earlier experiments, with a few new annotations of rare cell types (2-cell-state-like ESCs, motor neurons, midbrain-hindbrain-boundary (MHB)-like cells). In aggregate, cell type composition was also consistent with prior experiments, although there were substantially more PSC-like cells (**Figure S17B**).

We then constructed a monophyletic cell lineage tree for 58,283 cells with sufficient DTT recovery, leveraging the same custom DTT-based distance metric as previously described (**Figure 5D**; **Figure S18A**). However, rather than assigning cells to wells, we sought to identify a specific pseudo-ancestor in this phylogenetic tree that specifically corresponds to the monoclonal founder of each gastruloid. To do this, we first manually identified seven pseudo-ancestors near the root of the tree that, based on “TapeBC-Site-Edit” combinations exclusive to its Debris-seq wells, correspond to the founders of the seven major gastruloid clades (**Figure 5B**; **Figure S18B-C**). We then repeated this heuristic on each major clade, focusing on subclades of the phylogenetic tree defined by a given clade’s pseudo-ancestor. Overall, we were able to identify 96 pseudo-ancestors that we posit correspond to the founders of 96 gastruloids. The subclades defined by these 96 pseudo-ancestors collectively contain 51,963 (89%) of cells in the tree (541 +/- 704 cells per gastruloid/pseudo-ancestor; range: 2 to 5,319; **Figure S18D-E**). Although each assignment is strongly supported by Debris-seq, a further validation lies in the strong correlation that we observe between the number of cells recovered for a given pseudo-ancestor and the physical size of the corresponding gastruloid as estimated from brightfield imaging (**Figure 5B**, bottom; **Figure S18F**).

Finally, we sought to ask whether the phenotypic variation among gastruloids (second epoch) is informed by the lineage relationships of their founder mESCs (first epoch) (**Figure 5A**). As a first approach, we simply asked whether the gastruloids of major clades A-G are enriched for specific cell types. Indeed, Clade A is enriched for gastruloids containing a high proportion of PSC-like cells (*p =* 8 x 10^-3^, Wilcoxon test compared to 5,000 permutations), and Clade F for gastruloids containing a high proportion of hindbrain cells (*p =* 2 x 10^-4^) (**Figure 5E**; **Figure S19A**).

As a second approach, we sought to more systematically define the principal components of phenotypic variation among these gastruloids. The first three PCs of “pseudo-bulked” transcriptional profiles of 88 gastruloids (those with ≥50 assigned cells) explained two-thirds of the variance. PC1 was very strongly correlated with the proportion of cells in an undifferentiated PSC-like state (35.8% of variance; *r =* -0.92; *p =* 1 x 10^-36^), PC2 with the number of cells, a proxy for gastruloid size (18.3% of variance; *r =* 0.61; *p =* 4 x 10^-10^) and PC3 with the proportion of hindbrain-like cells (11.9% of variance; *r =* 0.58; *p =* 4 x 10^-9^) (**Figure 5F**; **Figure S19B**).

Are these phenotypic PCs correlated with the lineage relationships of their founder PCs? We examined whether “nearest neighbor” gastruloids, as defined by the first epoch, were phenotypically correlated. Modestly strong correlations were observed for both PC1 (*r* = 0.30; *p* = 1 x 10^-3^) and PC3 (*r* = 0.32; *p* = 5 x 10^-4^), but not PC2 (**Figure 5G**; **Figure S19C**). These correlations appear to be driven in part by the aforementioned enrichment of Clade A for PSC-rich gastruloids, and Clade F for hindbrain-rich gastruloids (**Figure 5G**). Of particular note, four gastruloids that have among the highest proportion of hindbrain cells and correspondingly the highest PC3 values, P1-B5 (79%), P2-E3 (60%), P2-E1 (38%) and P1-C8 (70%), compose a single subclade on the phylogenetic tree (purple box in **Figure 5B & G**). The result suggests that the mESC at the root of this subclade was biased to give rise to hindbrain-rich gastruloids, a proclivity that heritably maintained through final cell divisions of the first epoch, the dissociation and replating of the first epoch mESC aggregate, and the expansion of the second epoch mESC aggregates (**Figure 5A**).

To more generally test the correlation between founder mESC relatedness and gastruloid phenotype, we devised two metrics of gastruloid relatedness across all 3,828 possible pairs of 88 tips/gastruloids. The first metric calculates the Euclidean distance between a pair of gastruloids in a 10-dimensional space defined by the aforementioned PCA of “pseudo-bulked” gastruloid transcriptional profiles. In this case, we observe a positive correlation, *i.e.* phylogenetically distant founder mESCs gave rise to gastruloids whose pseudo-bulk transcriptional profiles were more distant from another (*rho* = 0.052; *p =* 1 x 10^-3^; **Figure S20A**). The second metric relies on a 30-dimensional space defined by PCA of all single cell transcriptional data from these 88 gastruloids and calculates, for each a given pair of gastruloids, the average percentage of nearest neighbors (*k* = 15) of cells of one gastruloid that derive from the other gastruloid. Here we observe a negative correlation, *i.e.* phylogenetically close founder mESCs gave rise to gastruloids whose cells were more similar than expected by chance (*rho* = -0.094; *p =* 6 x 10^-9^; **Figure S20B**). Although these metrics are not entirely independent (**Figure S20C**), the pattern across a range of phylogenetic distances supports the conclusion that there are aspects of mESC heterogeneity that arise during the first epoch, transmit to the second epoch, and contribute to gastruloid heterogeneity. A caveat is that errors during phylogenetic tree reconstruction confound this analysis, *e.g.* the misassignment of some cells to sibling gastruloids would be expected to inflate these correlations. However, the trends and significance are robust to the exclusion of pairs of gastruloids that are nearest neighbors (**Figure S20D**).

## DISCUSSION

Our study had two primary goals: first, to explore the origins of intra-batch, inter-individual heterogeneity in organoids, and second, to establish the DNA Typewriter lineage tracing system, whose proof-of-concept was limited to HEK293T cells^2^, in mammalian stem cells. To these ends, we developed an *in vitro* system in which a single mouse embryonic stem cell (mESC) serves as the founder of a "monoclonal gastruloid."

For the first goal, our strategy was inspired by Luria and Delbrück’s classic Fluctuation Test experiment^40^ from 1943, as well as recent work by Shaffer, Raj, and colleagues on clonal memory in cancer cells^41,42^. Much like how Luria and Delbrück demonstrated that bacterial resistance to phages arises from pre-existing, heritable mutations rather than being induced by phage exposure, our findings suggest that heterogeneity in gastruloids arises from pre-existing, heritable fluctuations in founder mESCs—both within the first few cell divisions (intra-gastruloid heterogeneity; **Figure 4**) and across different founders (inter-gastruloid heterogeneity; **Figure 5**)—rather than being stochastically induced by Wnt agonist exposure solely.

Focusing first on intra-gastruloid fate biases of mESCs (**Figure 4**), our findings align with recent work by Liberali and colleagues^25^ and by McNamara, Toettcher, and colleagues^39^, who demonstrated how spatial positioning in spheroids, differential Wnt exposure, and cell sorting rearrangements, shape individual cell contributions to gastruloids. Relatedly, a recent study by Chan, Smith, Meissner, Veenvliet, and colleagues^43^ observed diverse intra-gastruloid fate trajectories in a Trunk-Like Structure (TLS) model that is similar to gastruloids. Collectively, these studies suggest that fluctuations in stem cell states, such as Nodal/BMP signaling, drive physical rearrangements in aggregates, which in turn influence differential exposure to Wnt signaling and ultimately shape the emergence of the anterior-posterior axis. However, because these studies relied on polyclonal mESC aggregates, they could not determine when such fluctuations arise or how heritable they are across cell divisions. Leveraging DNA Typewriter in monoclonal gastruloids, our results suggest that mESC state fluctuations can be both sharp and heritable. For instance, in the Well16 gastruloid (**Figure 4C**), we infer that the first cell division created two daughter lineages, one of which was enriched in the gastruloid core and maintained pluripotency, while the other localized to the periphery and underwent primitive streak-like differentiation. This suggests that fluctuations occurring in early cell divisions persist over multiple generations, influencing fate decisions long before induction. On the other hand, this experiment did not fully rule out the possibility that the lineage relationships of cells are simply reflecting their spatial positions.

Our two-epoch experiment was specifically designed to isolate lineage effects from spatial influences on monoclonal gastruloids (**Figure 5**). Although the observed effects were more subtle, the data support the notion that founder mESC states influence gastruloid outcomes. Interestingly, this experiment exhibited less overall heterogeneity than our earlier experiments, likely due to the tight lineage relationships among founder mESCs, which we controlled by limiting the number of cell divisions in the first epoch. Even with this constraint, principal components of variation among gastruloids remained apparent, particularly with respect to the representation of particular cell types (PSC-like cells, hindbrain) and gastruloid size. Gastruloids originating from closely related founder mESCs exhibited correlated cell type compositions, reinforcing the idea that lineage-dependent fate biases emerge long before differentiation cues are introduced.

Much like how Luria and Delbrück’s work demonstrated the pre-existence of mutations before selective pressure, our findings suggest that pre-existing heterogeneity among mESC founders influences gastruloid cell type composition. Our results broadly align with recent studies on clonal memory in cancer cells^41,42^, highlighting the role of heritable fluctuations in cellular states. A key challenge for the field will be to define the molecular factors driving these fluctuations and to determine how they shape intra- and inter-gastruloid fate biases. We envision that future studies could leverage “rewind experiments”^44^ or “decorated lineage trees”^45^ to dissect this phenomenon further. The concurrent measurement of cell lineage and the spatial positions of cells would be particularly valuable for obtaining a comprehensive view of monoclonal gastruloid development.

In stem cell biology, shifting from polyclonal to monoclonal stem cell aggregates as the starting point for organoid differentiation protocols may offer distinct advantages for certain applications. For example, in a companion study^26^, we show that cellular bottlenecks in polyclonal embryoid bodies (EBs) drastically reduce clonal complexity, confounding mosaic genetic screens, whereas monoclonal EBs offer the possibility of generating genetically homogenous “individuals”. The establishment of additional protocols for deriving monoclonal organoids of various kinds could help to mitigate inter-individual heterogeneity or, alternatively, to harness it for specific research goals^26^.

Our second goal was to establish the DNA Typewriter lineage tracing system in mammalian stem cells. An important milestone of our study is demonstrating that DNA Typewriter, previously validated in HEK293 cells, can reconstruct large-scale phylogenies from a single founder to tens of thousands of differentiated cells within a single experiment. This establishes DNA Typewriter as a powerful tool for tracking lineage relationships in stem cells over extended timescales. Future enhancements, such as ENGRAM-based decorations and improved cell recovery rates, will further refine lineage tracing, enhancing both resolution and interpretability. Additionally, computational tools must evolve to fully leverage large-scale time-calibrated phylogenies at the resolution of millions to billions of cells. Our implementation of DNA Typewriter in monoclonal gastruloids also lays the foundation for extending these approaches to *in vivo* models, such as the mouse, a system wherein each individual also begins from a single stem cell.

## Lead contact

Correspondence and requests for materials should be addressed to Jay Shendure.

## Materials availability

Reagents generated in this study are available from the lead contact.

## Data & code availability

The data generated in this study can be downloaded in raw and processed forms from the NCBI Gene Expression Omnibus under accession number GSE291244. The code used here is available at: https://github.com/shendurelab/DTT.

## Acknowledgments

We thank the members of the Shendure lab for helpful discussions. This work was supported by the Brotman Baty Institute for Precision Medicine and grants from the Paul G. Allen Frontiers Group (Allen Discovery Center for Cell Lineage Tracing to J.S.), Seattle Hub for Synthetic Biology, a collaboration between the Allen Institute, Chan Zuckerberg Initiative and University of Washington (award number CZIF2023-008738 to J.S.), Damon Runyon Cancer Research Foundation (DRG-2507-23 to W.C.; DRG-2435-21 to J.-B.L.; DFS-64-24 to J.C.), Alex’s Lemonade Stand Foundation (Grant 19-15730), Crazy 8 Initiative (to J.S.), and the National Institutes of Health (1UM1HG011586 to J.S.; R01HG010632 to J.S.; R00HG012973 to J.C.; P30CA008748 partially supporting J.C.; F31HG011576 to S.G.R.). H.L. is supported by the NSF Graduate Research Fellowship Program (DGE-2140004). H.K. is a Washington Research Foundation Postdoctoral Fellow. S.D. was supported by the European Molecular Biology Organisation (ALTF 732-2017) and the Human Frontier Science Program (LT 000213/2018). J.S. is an Investigator of the Howard Hughes Medical Institute.

## Author Contributions

S.R., J.C., J.S. designed the research.

S.R., J.C. performed experiments with assistance from B.K.M., W.C., H.L., H.K., X.L., J.-B.L, N.H., S.D.

C.Q., J.C., S.K., S.R. performed computational analyses.

J.C., C.Q., S.R., J.S. collaboratively explored and annotated the data.

J.C., C.Q., J.S. wrote the manuscript, with assistance from S.R.

J.C., J.S. supervised the project.

## Methods

### Culturing mouse embryonic stem cells (mESCs)

We adopted a standard protocol for culturing mESCs, similar to the protocol described in ref ^46^. Our mESCs (E14TG2a) were a gift from C. Schröter. During regular passaging and expansion, mESCs were initially cultured on a gelatin-coated plate using 2i-LIF medium, which is NDiff227 medium (Takara) supplemented with 3 µM CHIR99021 (CHIR, Selleck, S2924), 1 µM PD0325901 (Selleck, S1036), and 1,000 units of ESGRO recombinant mouse LIF protein (Sigma-Aldrich, ESG1107). Gelatin-coated plates were prepared by making 0.2% (w/v) gelatin solution (Sigma, G1393) and autoclaving the bottle, which was applied to each culturing well (1 mL per well within a 6-well culturing plate) and incubated in a tissue culture incubator set to 37 °C with 5% CO2 for at least 30 minutes. After the coating, the gelatin solution was aspirated from the plate immediately before depositing mESCs. Cells are grown in the stem-cell-designated incubator (set to 37 °C with 5% CO2) and biosafety cabinet to avoid cross-contamination. After the initial expansion, mESCs were further adapted to the Serum-based medium, which is GMEM (Gibco) supplemented with 8% Fetal Bovine Serum (FBS, Biowest), 8% KnockOut Serum Replacement (KSR, Gibco), 1X Glutamax (Gibco), 1X MEM non-essential amino acids (Gibco), 1 mM Sodium Pyruvate (Gibco), and 0.1 mM beta-mercaptoethanol (BME, Gibco).

### Monoclonal gastruloid induction protocol

To induce monoclonal gastruloids, we perform a two-stage protocol, where the first stage generates a monoclonal mESC cell aggregate (also referred to as spheroid culture), and the second stage closely mirrors the standard gastruloid induction protocol with a 24-hr CHIR pulsing.

The protocol starts by culturing Mito-C inactivated MEFs (CF-1 MEFs, Applied StemCell) in 6-well plates with DMEM medium with high glucose (Gibco), supplemented with 15% FBS (Biowest), 1X Glutamax (Gibco), and 1X MEM non-essential amino acids (Gibco). On the next day (Day 1), we flow-sort mESCs to ensure single-cell resuspension and add them to each 6-well plate at a low cell concentration of around 600 cells per 6-well. Cells are grown in the standard mESC media for an additional five days, where we expect to observe cell colonies of 100- to 150-µm diameter that are isolated from the other mESC colonies. On Day 6, the MEF layer is selectively dissociated while keeping mESC colonies intact by washing each 6-well with 2 mL of PBS without calcium or magnesium twice, and the addition of 2 mL of Collagenase IV (1 mg/mL, StemCell Technologies) and incubation at 37 °C for 20-30 minutes. Once colonies are successfully lifted off of the MEF layer, they are carefully pooled into a 15-mL tube and allowed to settle by gravity. The remaining Collagenase IV solutions are carefully aspirated, and monoclonal colonies are washed with serum-containing mESC media (to inactivate any remaining Collagenase IV and to wash remaining MEFs away). Colonies are then transferred to a non-adherent 10-cm plate in the differentiation medium (NDiff227, Takara). After 24 hours, 3D spheroids or aggregates are manually picked into 96-well plates (non-adherent, round bottom) and are subsequently used for downstream gastruloid induction protocol.

For the second stage, after each aggregate is placed into a well in a 96-well plate, they are treated as gastruloids of the 24-hour-after-aggregation time point in the conventional mouse 3D gastruloid induction protocol^46^ (**Figure 1A**). One day after culturing in the 96-well plate, culturing media was replaced by NDiff227 supplemented with 3 µM CHIR for the next 24 hours, matching the CHIR treatment between the 48- and 72-hour timepoints in the conventional gastruloid induction protocol. After the 24-hour CHIR pulsing, the culturing medium is replaced with the original NDiff227 medium until the harvesting of gastruloids for single-cell profiling.

### Transduction of mESCs

For the experiment reported in **Figure 1**, we used lentiviral transduction to stably integrate a degenerate 8-mer barcode (NNNNNNNN) to mESCs. We used ViraPower Lentiviral Expression System (ThermoFisher) to generate lentivirus packaged with our construct encoding GFP-P2A-Puromycin resistance gene (PuroR) with the random 8-mer DNA barcodes positioned in the 3’-UTR of GFP transcript. The resulting lentivirus was concentrated overnight using Peg-it Virus Precipitation Solution (SBI) then aliquoted and flash-frozen using liquid nitrogen. Viral titers were generated using established protocols, where an aliquot of frozen virus was thawed and applied to mESCs at different dilutions (200 µL of diluted lentivirus solution to roughly 500,000 cells per well within a 6-well culturing plate). GFP fluorescence was used as a proxy for MOI efficiency, which was measured with FACS analysis. After the determination of low MOI conditions, cells were selected on puromycin to retain a polyclonal mESC population with cells harboring a unique static 8-mer barcode.

### Generation of polyclonal DNA Typewriter lineage tracing cell line

For experiments reported in **Figure 2** onwards, we generated DNA Typewriter lineage-tracing mESC lines via sequential integrations of piggyBac transposons, with each integration followed by selection steps. First, we generated PB-TAPE (**Figure S1B**), which includes two gene expression cassettes: 1) TAPE-targeting epegRNAs (random NNNGGA insertions) expressed from U6 promoter, and 2) GFP encoding DNA Tape in its 3’-UTR. DNA Tape is structured to include the T7 promoter sequence, a 12-bp Tape-specific degenerate barcode (TapeBC, NNNNNAANNNNN), 6xTAPE capable of six sequential insertions, and 10X Genomics Capture Sequence 1 (CS1) to aid recovery of DNA Tape during scRNA-seq. We also cloned PB-iPEmax-HygroR (**Figure S1B**), which encodes Doxycycline-inducible PEmax-P2A-mCherry and rtTA3-P2A-HygroR (reverse-tetracycline transactivator 3 and Hygromycin-resistance gene linked by P2A ribosomal skipping peptide sequence).

Next, we transfected PB-TAPE with the Super piggyBac transposase expression vector (SBI) for stable integration. We mixed PB-TAPE, PB-PuroR, and piggyBac transposase plasmids with the 90:5:5 mass ratio, then used Lipofectamine 2000 (ThermoFisher) protocol to transfect mESCs (4 µg per 1 million cells in 6-well plate). Transfected cells were cultured for another 7 days, and placed under puromycin selection (1 µg/mL) for two days. The resulting cells were mostly GFP-positive when imaged under the fluorescent microscope, possibly due to the high copy number of PB-TAPE integrants in each cell.

Finally, we performed another transfection of PB-TAPE, PB-iPEmax-HygroR, and piggyBac transposase, mixed at the 85:10:5 mass ratio. The resulting cells were cultured for another 7 days, selected using hygromycin (150 µg/mL) for four days, and then frozen.

### sci-RNA-seq3 monoclonal gastruloid induction protocol

Polyclonal DNA Typewriter lineage tracing cells were thawed and passaged for at least a week. Recovered cells were FACS-sorted for GFP-positivity, with the upper 50% retained. Doxycycline (100 ng/mL) was added to the cell culture media a day before the sorting, and omitted in the cell culture after the seeding. Sorted cells were resuspended as single cells, diluted to 600 cells per 2 mL in Serum-medium, and plated on three 6-well plates containing the MEF cells, following the monoclonal gastruloid induction protocol described above. After the monoclonal colony separation step and formation of a spherical cell aggregate each from monoclonal mESC colony, we picked 144 cell aggregates into one-and-a-half 96-well plates. Gastruloids were grown for another three days after CHIR treatment. At the time of gastruloid harvesting, the culturing media from each well containing a gastruloid was collected for the Debris-seq protocol described below.

### Data generation and processing for the sci-RNA-seq3 experiment

For the application of sci-RNA-seq3 to mouse gastruloids, we used an adaptation of the optimized sci-RNA-seq3 protocol that includes a T7 transcription step^27,28,47^. 144 monoclonal gastruloids collected in three pools (each containing 48 gastruloids) were gently resuspended in 2 mL lysis buffer to free the nuclei and fixed with methanol (80% methanol in a total of 10ml) for storage at -80 °C with at least 2 million nuclei per pool. Samples were thawed on ice and 10 mL of SPBSTM (sucrose, PBS, triton, magnesium chloride) buffer plus 100 µL DEPC (Millipore Sigma) was added to rehydrate the nuclei. Nuclei were centrifuged at 500 xg for 3 minutes at 4 °C and resuspended in 60 µL SPBSTM for the T7 reaction (NEB HiScribe kit #E2050S), adding 75 µL of the NTP/buffer mix and 15 µL of the enzyme mix. The nuclei were incubated at 37 °C for 30 minutes, then the volume was brought up to 1 mL with SPBSTM plus 50 µL 100 mM BS3 to fix the nuclei again, 10 minutes on ice, then centrifuged. Nuclei were resuspended at 2 million per 500 µL, and 56 µL of dNTPs were added and then distributed to a 96-well plate for reverse transcription. A separate sci-RNA-seq3 experiment was conducted for each of the three pools. Two types of indexed reverse transcription primers were used in each well: an oligodT primer for the transcriptome, and another specific for the DTT construct. After reverse transcription, nuclei were pooled and redistributed to a second plate to attach a second index by ligation. Then nuclei were pooled and redistributed again into their final plates (six plates in total, where two plates were prepared from each pool containing 48 gastruloids) for second-strand synthesis. The nuclei were dissolved with protease and then split: 5 µL of 10 mM Tris was added to each well and then immediately removed to a second plate to divide each well into two. One half was taken through the remaining sci-RNA-seq3 protocol with tagmentation and PCR for the transcriptome. The other half was taken directly to a PCR specific to the DTT construct, using the normal sci-RNA-seq3 P5 indexed primer on one side and a custom P7 primer (CAAGCAGAAGACGGCATACGAGATNNNNNNNNNNGGTACCATAGCAGATGATCCATGGTC) on the other side, to specifically enrich for the DTTs. PCRs for the transcriptome and enrichment were sequenced independently on an Illumina Nextseq 2000.

Transcriptome data from each of the six individual plates were processed independently. For each plate, read alignment and gene count matrix generation were performed using the pipeline that we developed for sci-RNA-seq3 (https://github.com/JunyueC/sci-RNA-seq3_pipeline). Briefly, base calls were converted to fastq format using Illumina’s *bcl2fastq*/v2.20 and demultiplexed based on PCR i5 and i7 barcodes using maximum likelihood demultiplexing package *deML*^48^ with default settings. Demultiplexed reads were filtered based on the reverse transcription (RT) index and hairpin ligation adaptor index (Levenshtein edit distance (ED) < 2, including insertions and deletions) and adaptor-clipped using *trim_galore*/v0.6.5 (https://github.com/FelixKrueger/TrimGalore) with default settings. Trimmed reads were aligned to a custom mouse reference genome (GRCm39 with an additional DTT sequence) using *STAR*/v2.6.1d^49^ with default settings and gene annotations from GENCODE VM31. Mapped reads were extracted, and duplicates were removed using the unique molecular identifier (UMI) sequence, RT index, ligation index, and read 2 end-coordinate (that is, reads with identical UMI, RT index, ligation index, and tagmentation site were considered duplicates). Finally, mapped reads were split into constituent cellular indices by further demultiplexing reads using the RT index and ligation index. To generate digital expression matrices, we calculated the number of strand-specific UMIs for each cell mapping to the exonic and intronic regions of each gene with the *python*/v2.7.13 *HTseq* package^50^. For multi-mapping reads (*i.e.* those mapping to multiple genes), the read were assigned to the gene for which the distance between the mapped location and the 3’ end of that gene was smallest, except in cases where the read mapped to within 100 bp of the 3’ end of more than one gene, in which case the read was discarded. For most analyses, we included both expected-strand intronic and exonic UMIs in per-gene single-cell expression matrices. After the single-cell gene count matrix was generated, cells with low quality (UMI < 200 or detected genes < 100 or unmatched_rate (proportion of reads not mapping to any exon or intron) ≥ 0.4) were filtered out.

We performed two steps to exhaustively detect and remove potential doublets. Of note, all these analyses were performed separately on data from each plate. First, we applied the *scrublet*/v0.1^51^ pipeline to each dataset with parameters (min_count = 3, min_cells = 3, vscore_percentile = 85, n_pc = 30, expected_doublet_rate = 0.06, sim_doublet_ratio = 2, n_neighbors = 30, scaling_method = ‘log’) for doublet score calculation. Cells with doublet scores over 0.2 were annotated as detected doublets. Second, we performed two rounds of clustering and used the doublet annotations to identify subclusters that are enriched in doublets. The clustering was performed based on *Scanpy*/v.1.6.0^52^. Briefly, gene counts mapping to sex chromosomes were removed, and genes with zero counts were filtered out. Each cell was normalized by the total UMI count per cell, and the top 3,000 genes with the highest variance were selected, followed by renormalizing the gene expression matrix. The data was log-transformed after adding a pseudocount and scaled to unit variance and zero mean. The dimensionality of the data was reduced by PCA (30 components), followed by Louvain clustering with default parameters (resolution = 1). For the Louvain clustering, we first computed a neighborhood graph using a local neighborhood number of 50 using *scanpy.pp.neighbors*. We then clustered the cells into sub-groups using the Louvain algorithm implemented by the *scanpy.tl.louvain* function. For each cell cluster, we applied the same strategies to identify subclusters, except that we set resolution = 3 for Louvain clustering. Subclusters with a detected doublet ratio (by *Scrublet*) over 15% were annotated as doublet-derived subclusters. We then removed cells which are either labeled as doublets by *Scrublet* or that were included in doublet-derived subclusters. Altogether, 7.4% to 11.1% of cells in each plate were removed by this procedure.

After removing the potential doublets detected by the above two steps, we further filtered out the potential low-quality cells with the doublet score (by *Scrublet*) greater than 0.15, the percentage of reads mapping to ribosomal chromosomes (Ribo%) > 5, the percentage of reads mapping to exonic regions (exon%) > 85, or the percentage of reads mapping to mitochondrial chromosomes (Mito%) > 10. Subsequently, we determined thresholds for UMI counts per cell used for quality filtering, set at the mean plus 2 standard deviations and mean minus 1.5 standard deviations of log2-scaled values. Cells with fewer than 250 detected UMIs were further excluded. Following this, we merged cells from individual plates to generate the final dataset, composed of 247,064 cells. The median UMI count per cell was 609, with a median gene count detection per cell of 512.

We then applied *Seurat/v3*^53^ to this final dataset, performing conventional single-cell RNA-seq data processing: 1) retaining protein-coding genes, lincRNA, and pseudogenes for each cell and removing gene counts mapping to sex chromosomes; 2) normalizing the UMI counts by the total count per cell followed by log-transformation; 3) selecting the 2,500 most highly variable genes and scaling the expression of each to zero mean and unit variance; 4) applying PCA and then using the top 30 PCs to calculate a neighborhood graph, followed by louvain clustering (resolution = 1); 4) performing UMAP visualization in 2D space (min.dist = 0.3). For cell clustering, we manually adjusted the resolution parameter towards modest over-clustering, and then manually merged adjacent clusters if they had a limited number of differentially expressed genes (DEGs) relative to one another or if they both highly expressed the same literature-nominated marker genes. Subsequently, we annotated individual cell clusters using at least two literature-nominated marker genes per cell type label (**Table S1**).

DTT data from six individual plates (two plates per a pool of 48 gastruloids) were processed separately using the same pipeline: base calls were converted to FASTQ, demultiplexed, filtered by RT and hairpin ligation indices, trimmed, and aligned to a custom mouse genome (GRCm39 + DTT). Mapped reads were extracted for analysis.

### Debris-seq data generation and processing

Cell culture medium from each well containing a monoclonal cell aggregate was collected to purify genomic DNA from cell debris (Debris-seq). Each well was carefully pipet-mixed to resuspend any settled cell debris without disturbing the cell aggregates before removing the culturing medium. The culturing medium containing cell debris was transferred to a PCR tube strip containing 8 tubes and spun down using the table-top microfuge for 20 minutes at around 2000 xg (MyFuge, Benchmark). The culturing medium was carefully removed without disturbing the cell-debris pellet. The genomic DNA from each cell-debris pellet was purified using the DNA extraction kit (ARCTURUS PicoPure DNA Extraction Kit, ThermoFisher), where each pellet was resuspended in 10 to 15 µL of DNA extraction buffer. The DNA Tape sequences within the resulting genomic DNA were PCR-amplified, as described previously^2,54^. Briefly, about 2 µL of the genomic DNA solution was added to 50 µL of the PCR reaction (KAPA2G Robust HotStart ReadyMix), where PCR primers (0.4 µM final concentrations) added Illumina sequencing primer binding sequences. PCR reactions were performed for 3 minutes at 95 °C; 15 seconds at 95 °C, 10 seconds at 65 °C and 40 seconds at 72 °C for 30 cycles; and 1 minute at 72 °C. A portion of the resulting PCR reactions (1 µL) is then used for the subsequent PCR reaction (25 µL), which uses primers that add Illumina sequencing adaptors and dual-index sequences for the future demultiplexing of samples from the sequencing run. The same PCR reaction parameters were used, except for the reduction of cycling numbers to 6 cycles in total. The resulting libraries were pooled, cleaned up using magnetic beads (AMPure XP, Beckman Coulter), and sequenced on Illumina Nextseq2000.

The resulting sequencing runs were demultiplexed by their indices, generating a single fastq file per aggregate-bearing well. Fastq files were analyzed using a custom Python script that performed the following three steps: 1) For each sequencing read, it extracted TapeBC and DNA Typewriter Tape Site1-6 insertions (NNNGGA) based on their amplicon positions. The script goes through Site1 to Site6 sequentially, checking for insertions from the first site (“InsertBC(1)”) to the last (sixth) site (“InsertBC(6)”) on Tape. 2) Reads with the same TapeBC and editing patterns on InsertBC(1-6) are tabulated as a unique TapeBC-edit pattern and its frequency. 3) TapeBC and the InsertBC(1) are combined (e.g., TTTTTAAGCAAT-CCTGGA, where the first 12-bp is the NNNNNAANNNNN TapeBC and the second 6-bp is the insertion detected in the first site), and only the most frequent insertion pattern (single Site1-6 insertion pattern per a unique TapeBC-InsertBC(1) detected) is kept in the final output, with an additional threshold of read frequency to remove any combinations that are observed less than 0.2% of the total read. The resulting file includes the well position in the first column, TapeBC-InsertBC(1) in the second column, Site1-Site6 in the following six columns, and the read number in the last column.

### Mapping cells acquired from sci-RNA-seq3 to their gastruloid origins

We assigned each single-cell transcriptional profile to one of the 144 wells in which individual gastruloids were grown, as a proxy for their gastruloid-of-origin. Our heuristic for grouping/assigning cells had the following steps:

1. From the Debris-seq output, we defined unique DTTs using “TapeBC + InsertBC(1)” and retained only the most abundant combination of InsertBC(2)-(6) for each DTT within each well. We focused on 4,995 unique DTTs observed in Debris-seq data as confirmed candidates (*i.e.* “white list”), each exclusively appearing in one of 138 wells. The remaining 6 wells were excluded because they had no unique TapeBCs assigned.
2. In the sci-RNA-seq3 dataset, we excluded cells with fewer than three detected DTTs and then aligned the DTTs to the confirmed list from Debris-seq, allowing for a 1 bp mismatch on the 12-bp TapeBC while requiring exact matches on the first 4 editing sites. To account for potential differences in induction times between Debris-seq and scRNA-seq, we allowed ‘none’ editing on the 2nd, 3rd, and 4th editing sites of DTTs in the scRNA-seq to match any editing pattern in the DTTs from Debris-seq. This resulted in a matrix of DTTs by cells with read numbers as values. We excluded cells with fewer than three distinct DTTs detected, resulting in a final count of 154,988 cells and 4,609 DTTs.
3. We quantified the overlap between DTTs observed in cells vs. wells, and assigned 129,853 cells in cases where the largest overlap exceeded the second-largest overlap by at least 50%.
4. To make additional assignments, we performed principal components analysis (PCA)-based dimensionality reduction of scRNA-seq profiles based solely on DTTs matrix rather than transcriptional profiles. DTTs with less than 10 reads across all cells were filtered out. We applied *Seurat/v5*^55^ to normalize the read counts by the total count per cell, followed by log-transformation and scaling each DTT to have a zero mean and unit variance. Next, PCA was applied, and the top 100 PCs were selected to calculate a neighborhood graph. This was followed by UMAP visualization in a 2D space (min.dist = 0.1). For unassigned cells, we identified the wells of their top 10 nearest already-assigned neighbor cells in PCA space (n = 100), and assigned a further 19,363 cells in cases where the largest overlap exceeded the second largest overlap by at least 50%.
5. After filtering out gastruloids with fewer than 100 cells, a total of 148,724 cells (96% of 154,988 cells) were allocated to 121 gastruloids. On average, each gastruloid contained 1,229 cells, with a median of 1,066 cells. The maximum cell count reached 5,883.
6. We identified the DTTs detected in at least 5% of the assigned cells, and then evaluated Jaccard similarities between the DTTs of cells assigned to each gastruloid and those detected from Debris-seq for each gastruloid.

### Lineage tracing monoclonal gastruloid induction protocol (8 gastruloids)

We generated a highly efficient DNA Typewriter lineage recording cell line by characterizing the recording efficiency of each monoclonal cell line. We started from the ongoing culture of the polyclonal DNA Typewriter lineage tracing mESC line (described above in the “Generation of polyclonal DNA Typewriter lineage tracing cell line” section), then used FACS to sort a high GFP positive population (1% GFP brightness). The resulting single-cell sorted cells were plated on the layer of MEF to grow monoclonal colonies, which were picked, individually dissociated, and plated into 36 wells in a 96-well plate coated with 0.1% Gelatin. They were expanded until they reached a confluent well in a 6-well, where each culture was split into three parts: 1) frozen to preserve, 2) cultured for another 2 days in a 96-well, and 3) cultured for 2 days in a 96-well in the presence of 100 ng/mL Doxycycline to induce genomic recording. Cultured cells were harvested after 2 days and lysed to collect genomic DNAs. DNA Tapes from genomic DNA samples were PCR-amplified and sequenced to quantify the editing with or without Doxycycline induction. Among 36 cell lines, we identified three (named Clone-05, Clone-25, and Clone-32) that had the highest editing efficiency with Doxycycline induction and minimal editing without the induction.

We used these three cell lines, mixed at equal numbers, to induce monoclonal cell lines for high-resolution lineage reconstruction. We added 100 ng/mL Doxycycline into the culturing medium throughout the monoclonal gastruloid induction protocol, starting a day before the single-cell sorting step. We diluted 3600 sorted cells to 12 mL of Serum-based medium, then distributed them across wells of a single 6-well plate containing MEF to support monoclonal colony formation. 32 monoclonal cell aggregates were picked into a 96-well plate, and treated with CHIR for 24 hours according to the monoclonal gastruloid induction protocol. A day after the CHIR-treatment step (equivalent to the 96-hr after-aggregation timepoint in the conventional gastruloid induction protocol), we collected cell culture medium from each well and performed Debris-seq to access the editing efficiencies within each cell aggregate. All aggregates showed relatively high editing efficiencies (34 ± 12 % of the third sites within the DNA Typewriter Tapes edited). Debris-seq also provided cell line information of each monoclonal gastruloid, by matching the recovered TapeBCs with previously profiled TapeBCs from each of the three cell lines. We used this information to select 8 gastruloids to profile using 2 lanes of 10X Genomics scRNA-seq kit (Well14, Well17, Well25 into the first reaction lane containing cells from Clone 05, and Well01, Well03, Well21, Well16, and Well28 into the second reaction lane containing cells from Clone 25 and 32), dissociated and processed following the conventional protocols (Cell Preparation guide, CG00053 Rev C, and User Guide for Chromium Next GEM Single Cell 3’ HT Reagent Kits v3.1, Rev D).

### Data generation and processing for the “eight monoclonal gastruloids” experiment

The single-cell transcriptome data from each 10X Genomics scRNA-seq lane were processed separately using Cell Ranger 7.2.0^56^ with default settings (e.g., --include-introns true) and refdata-cellranger-mm10-3.0.0 as the reference. We extracted the gene-by-cell matrix from the ‘raw_feature_bc_matrix’ folder, filtered out cells with UMI counts below 500 or fewer than 250 detected genes, and retained genes from chromosomes 1-19, X, Y, and MT. We then detected doublets using the Scrublet/v0.1^51^ pipeline and calculated the percentage of reads mapping to either mitochondrial (i.e. MT%) or ribosomal chromosomes (i.e. Ribo%) for individual cells. After manually examining the distribution of UMIs and MT% across cells, we applied the following criteria to filter out potentially low-quality cells:

- Log2 UMI Counts: Excluded cells with counts below 12.5 in lane 1 or below 12 in lane 2, as well as those exceeding the top 0.5% of total cells.
- Doublet Scores: Removed cells with doublet scores above 0.2, as calculated by Scrublet.
- MT%: Excluded cells with MT% over 10% or below 1%.
- Ribo%: Removed cells with Ribo% over 40%.

After combining cells from two lanes, we applied Seurat/v5^55^ to this final dataset, performing conventional single-cell RNA-seq data processing: 1) retaining protein-coding genes and lincRNA for each cell and removing gene counts mapping to sex chromosomes; 2) normalizing the UMI counts by the total count per cell followed by log-transformation; 3) selecting the 2,500 most highly variable genes and scaling the expression of each to zero mean and unit variance; 4) applying PCA and then using the top 30 PCs to calculate a neighborhood graph, followed by louvain clustering; 5) performing UMAP visualization in 2D space (min.dist = 0.3). For cell clustering, we manually adjusted the resolution parameter towards modest over-clustering, and then manually merged adjacent clusters if they had a limited number of differentially expressed genes (DEGs) relative to one another or if they both highly expressed the same literature-nominated marker genes (**Table S1**). Subsequently, we annotated individual cell clusters using at least two literature-nominated marker genes per cell type label.

Next, we processed the sequencing reads of DTT captured in the same scRNA-seq experiment. We extracted cell-specific barcode (CellBC), UMI added during cDNA synthesis, TapeBC and 6 InsertBC from each DTT sequencing reads, removed reads with less than 2 UMI associated with particular CellBC-TapeBC-6xInsertBC combinations, removed reads without CellBC matching to ones recovered from the single-cell transcriptomic library, removed reads without TapeBC matching to previously characterized ones from three clonal cell lines after correcting a single base-pair mismatch. In our data set, we often observed recovery of multiple 6xInsertBC combinations per CellBC-TapeBC combination, which suggested the existence of multiple DTT integrants with the same TapeBC. This may have occurred due to insufficient complexity of TapeBC in the DTT library, transposase-mediated repeated integration during DNA replication, or duplication of ploidy within the mESC. To allow discernment of unique integration within the genome during “cell-to-well” assignments, we used the first InsertBC (InsertBC(1)) as part of the TapeBC that identified a unique integration. Insertions are identified by their expected position within the amplicon sequencing read and additional 1-nt in its 3’-end, and searching for GGAT sequences in the last three position of NNNGGA insertions and the next base T. In case there are multiple 6xInsertBC combinations assigned to each CellBC-TapeBC-InsertBC(1) combination, we chose the InsertBC(2-6) combination with most UMI and sequencing read counts.

From either Debris-seq or scRNA-seq data, we defined unique DTTs with “TapeBC + InsertBC(1)” and retained only the most abundant InsertBC(2)-(6) edit combination for each DTT within each well or cell. We assigned cells to specific wells/gastruloids by matching the DTTs detected in individual cells with those identified in individual wells using Debris-seq. The assignment strategy was generally consistent with our approach for mapping cells obtained from sci-RNA-seq3 to their gastruloid origins, with adjustments made to account for differences in cell number and sequence coverage:

1. We combined DTTs detected from eight wells across three clones in Debris-seq, retaining those that were either unique to a well or overlapped across multiple wells.
2. In the scRNA-seq dataset, we excluded cells with fewer than two detected DTTs or with a total DTT UMI count of less than 10.
3. For additional assignments, we performed dimensionality reduction on the cells using only the DTT matrix. For unassigned cells, we identified the wells of their top 10 nearest assigned neighbor cells in PCA space (n = 30) and assigned them if the largest overlap exceeded the second largest overlap by at least 50%.
4. A total of 8,545 cells were allocated to 8 gastruloids. On average, each gastruloid contained 1,068 cells, with a median of 809 cells, ranging from 312 to 2,696.

### Inferring DTT integration events and edit patterns

A major challenge in analyzing DTT data is that TapeBCs do not uniquely define genomic loci, as most DTTs are duplicated—likely due to the high MOI transposase system. This duplication complicates lineage reconstruction, making it difficult to compare edit patterns across cells without knowing the originating DTT locus. To address this, we developed a method to infer the most likely number of integration events per TapeBC into the genome as well as the edit patterns that were generated from each integration locus, capturing the true sequence of events that occur from the starting cell at each locus independently. The key principle on which this method was based was the idea that in a given cell, for a given TapeBC, there should exist only one observed pattern of edits. If multiple patterns of edits with the same TapeBC in the same cell are observed, this is evidence for multiple genomic integration sites for that TapeBC and associated DTT.

From the scRNA-seq DTT data, we generated a table where each row represents a unique sequence defined by the CellBC, TapeBC, six editing sites (InsertBC(1) to InsertBC(6)), and the UMI count for that exact sequence. For each TapeBC across all cells, we computed a pairwise Hamming distance matrix for unique edit site sequences and grouped identical sequences into clusters if they contained more than 100 UMIs. UMIs not assigned to any cluster were discarded as likely noise from PCR or sequencing errors. The consensus sequence for each cluster was then refined to include only edited regions by identifying the edit key “GGAT” every 7 bases.

After identifying consensus edit patterns, we generated a cell-by-pattern matrix with UMI counts, row-normalized, and log-transformed to highlight differences between cells. A binarization threshold was set at the lowest point between distribution modes, producing a more interpretable matrix indicating whether a pattern is present in each cell for a given TapeBC. We then clustered the binary matrix using DBSCAN, grouping cells with the same sets of expressed patterns. Cells not assigned to a cluster, typically due to dropout, were classified using KNN in normalized log-UMI space. Each cluster’s binary expression vector was then used to compute a mutual exclusivity matrix via cosine distance with itself, forming a graph where nodes represent edit patterns and edges indicate mutual exclusivity (meaning the patterns are not co-expressed in the same cells). We intersected these graphs across all clusters to create a final graph representing relationships between edit patterns across all cells, assuming that in a monoclonal experiment, co-expressed patterns in any cluster of cells must have originated from distinct loci.

To group edit patterns into shared loci of origin, we used the mutual exclusivity graph to identify complete subgraphs (cliques) as putative integration events. We searched for a clique partition that: 1) minimized the number of inferred integration events; and 2) maximized the sum of edge weights, which reflect shared consecutive edits between patterns- in other words, the most biologically parsimonious solution. To achieve this, we applied graph coloring to the complement graph, using a greedy algorithm to assign nodes to cliques. By running this algorithm multiple times with randomized node orders, we selected the solution with the fewest cliques and the highest sum of edge weights. Each inferred clique represents a distinct integration locus for a given TapeBC, grouping edit patterns accordingly. This resulted in a final matrix where rows represent cells and columns correspond to inferred loci, serving as the basis for calculating pairwise lineage distances. More details are available in the **Supplementary Note**.

### Generating phylogenetic trees from DTT data

To reconstruct the phylogenetic tree from genomic recordings, we first filtered out DTTs found in fewer than 10 cells for each clone (Clone-05, Clone-25, and Clone-32). We then excluded cells with DTT counts outside the range of mean - 1.5 × SD to mean + 2 × SD. To calculate cell-cell distances within each clone, we retained DTTs observed in both cells and computed an edit distance for each DTT (each containing six edit sites) between the two cells. This distance represents the number of steps required to transform the DTT in one cell to match the other, including reversing edits back to unedited and then applying forward edits. For example, the distance between (A C D - - -) and (B - - - - -) is 4. The possible distances range from 0 to 12.

The average edit distance across all shared DTTs between pairs of cells was calculated, followed by the generation of a phylogenetic tree using the UPGMA method, implemented in the ape package^57^ in R.

### Bootstrap analysis

We conducted a bootstrapping analysis to assess the robustness of the trees for individual clones or gastruloids. For each of the 100 bootstraps, we resampled an equal number of DTTs (each containing six edit sites) with replacements to reconstruct a new tree. Clade concordance was evaluated by comparing each bootstrap tree to the original and calculating the transfer bootstrap expectation (TBE)^35^. Specifically, the TBE for each clade in the original tree was determined by finding the minimal transfer distance across all clades in the bootstrap tree, followed by computing the mean values across the 100 bootstraps. A TBE cutoff of 0.7 was used, with values above this threshold indicating moderate to strong support.

### PATH analysis

PATH (phylogenetic analysis of trait heritability) adapts Moran’s I to quantify either the autocorrelation (within cell types) or the cross-correlations (between cell types) of cell phenotypes across the phylogenetic tree ^37^. If closely related cells exhibit greater phenotypic resemblance (i.e. from the same cell types) than random cells, the phenotype is considered autocorrelated. To perform the PATH analysis, we created two data matrices: one representing cell-cell phenotypic distances from the phylogenetic tree and the other as a cell-by-cell type matrix with binarized values indicating whether each cell belongs to a specific cell type. Each cell type was then scaled to have a zero mean and unit variance.

### “Tree-of-trees” experiment

We performed a two-stage “tree-of-trees” experiment by inducing multiple monoclonal gastruloids from a single starting mESC. From the mESC culture (Clone-05), we used FACS to sort around 3,600 cells, which were plated across a 6-well plate containing MEFs. The serum-based medium contained Doxycycline (100 ng/mL) throughout the experiment, starting from a day before the cell sorting (D0). Five days after the cell plating (D6), nine monoclonal colonies were isolated from MEFs, and cultured in a non-adherent, round-bottom 96-well for each to form a spherical aggregate. On next day (D7), the culturing medium from each well was collected for Debris-seq, and each cell aggregate was single-cell dissociated using Trypsin and plated into a culturing well with MEFs within a 6-well plate, totaling 9 wells across one-and-a-half 6-well plates. Based on the editing efficiency measured from the Debris-seq result, we selected one well for monoclonal colony isolations (D13). After culturing isolated monoclonal cell aggregates in a non-adherent 10-cm culturing plate to form spherical cell aggregates, we transferred 108 aggregates into two 96-well plates the next day (D14), followed by the CHIR treatment and the rest of the monoclonal gastruloid induction protocols. During the CHIR treatment, we noticed that we lost two gastruloids (second 96-well plate, well B10 and D6, or P2-B10 and P2-D6), possibly removed along with the culturing medium during the medium replacement. We collected culturing medium from each well for Debris-seq on D17 (including the missing well, which may contain cell debris), and harvested gastruloids on D18 (equivalent to a 144-hour after-aggregation time-point in the conventional gastruloid induction protocol) for the 10X Genomics scRNA-seq protocol.

### Data generation and processing for the “tree-of-trees” experiment

We dissociated all 106 gastruloids and profiled them across eight 10X Genomics lanes (v3.1-HT). The single-cell transcriptome data from each lane were processed separately using Cell Ranger 7.2.0^56^ with default settings (e.g., --include-introns true) and refdata-cellranger-mm10-3.0.0 as the reference. We extracted the gene-by-cell matrix from the ‘raw_feature_bc_matrix’ folder, filtered out cells with UMI counts below 500 or fewer than 250 detected genes, and retained genes from chromosomes 1-19, X, Y, and MT. We then detected doublets using the Scrublet/v0.1^51^ pipeline and calculated the percentage of reads mapping to either mitochondrial (i.e. MT%) or ribosomal chromosomes (i.e. Ribo%) for individual cells. After manually examining the distribution of UMIs and MT% across cells, we applied the following criteria to filter out potentially low-quality cells:

- Log2 UMI Counts: Excluded cells with counts below 10.5 (or 11 in some lanes) or exceeding the top 0.5% of total cells.
- Doublet Scores: Removed cells with doublet scores above 0.2, as calculated by Scrublet.
- MT%: Excluded cells with MT% over 10% or below 1%.
- Ribo%: Removed cells with Ribo% over 40%.

After combining cells from eight lanes, we applied Seurat/v5^55^ to this final dataset, performing conventional single-cell RNA-seq data processing: 1) retaining protein-coding genes and lincRNA for each cell and removing gene counts mapping to sex chromosomes; 2) normalizing the UMI counts using the SCTransform function^58^ implemented in Seurat; 3) applying PCA and then using the top 30 PCs to calculate a neighborhood graph, followed by Louvain clustering; and 4) performing UMAP visualization in 2D space (min.dist = 0.3). For cell clustering, we manually adjusted the resolution parameter towards modest over-clustering, and then manually merged adjacent clusters if they had a limited number of differentially expressed genes (DEGs) relative to one another or if they both highly expressed the same literature-nominated marker genes (**Table S1**). Subsequently, we annotated individual cell clusters using at least two literature-nominated marker genes per cell type label.

### Generating the “tree of trees”

The phylogenetic tree of 108 gastruloids was constructed based on gastruloid-gastruloid distances derived from monoclonal gastruloid-specific edits of DTTs captured by Debris-seq. To calculate the distance between two gastruloids, the following steps were performed: For each gastruloid, at each of the six sites across all 76 DTTs, the percentage of reads for each specific edit was determined, and combinations with at least 20% of reads were identified as dominant “TapeBC-Site-Edit” combinations. The Jaccard similarity between the two gastruloids’ sets of dominant combinations was calculated, with (1 - similarity) representing their distance. The phylogenetic tree was generated using the UPGMA method in the ape package^57^ in R.

### Assigning “founders” to pseudo-ancestors

For the DTT data extracted from scRNA-seq, as described above, we split some TapeBCs to account for piggyBac excision and re-integration events, resulting in a matrix of 65,465 cells × 231 DTTs (74 distinct TapeBCs). We then filtered out DTTs observed in fewer than 10 cells and excluded cells with a number of observed DTTs outside the range of mean - 1.5 × SD to mean + 2 × SD. Cell-cell distances were calculated as described previously by retaining DTTs observed in both cells and computing the edit distance for each DTT (containing six edit sites). This distance reflects the number of steps required to transform one DTT into another, including reversing edits and applying forward edits. The average edit distance across all shared DTTs between cell pairs was then calculated, followed by phylogenetic tree generation using the UPGMA method in the ape package^57^ in R.

To assign each of the seven major clades from the “tree of trees” to a pseudo-ancestor in the single-cell phylogenetic tree, we constructed a cell × “TapeBC-Site-Edit” combination matrix (where 1 indicates the combination was detected in the cell and 0 was not) and normalized it by the sum of each column. For each major clade in the “tree of trees”, we identified its exclusively dominant “TapeBC-Site-Edit” combinations. For each pseudo-ancestor in the single-cell phylogenetic tree, we selected the corresponding subset of cells and calculated the fold change between the mean value of dominant combinations for a given major clade and the mean value of dominant combinations across the other six major clades. This step generated a matrix where each row represents a pseudo-ancestor, each column corresponds to a major clade, and the values indicate fold-change. Next, for each major clade, we selected all pseudo-ancestors where the ratio of log2(fold-change + 1) between the given clade and the highest among the remaining six clades was greater than 2, and the one with the highest number of cells was designated as the representative of the major clade.

For each of the seven major clades, we subsetted the cells from its assigned pseudo-ancestor and the gastruloids within that clade, then repeated the assignment. For each gastruloid, we identified its exclusively dominant “TapeBC-Site-Edit” combinations. Then, for each pseudo-ancestor, we calculated the fold change between the mean value of dominant combinations for a given gastruloid and the mean across other gastruloids. This step generated a matrix where each row represents a pseudo-ancestor, each column corresponds to a gastruloid, and the values indicate fold-change. Next, for each gastruloid, we selected all pseudo-ancestors where the ratio of log2(fold-change + 1) between the given clade and the highest among the remaining six clades was greater than 2, and the one with the highest number of cells was designated as the representative of the gastruloids. Through this heuristic, a total of 96 gastruloids were assigned distinct pseudo-ancestors that we believe to correspond to their founding mESC, with these clades collectively comprising 51,963 cells. One gastruloid (P2-B10) was excluded due to the absence of exclusive dominant combinations, and 11 (P1-B10, P2-C9, P1-A5, P1-A4, P2-C3, P2-D12, P1-D9, P1-D8, P1-D6, P2-A12, and P2-D8) were excluded because they could not be assigned to a distinct pseudo-ancestor.

## Supplementary Tables

<u>Table S1.</u> Marker genes used for mouse 0gastruloids cell type annotation.

## Supplementary Note

### Inferring tape integration events and patterns of edits

To achieve high resolution lineage trees, we developed an mESC line with a high DNA Typewriter Tape copy number via high MOI piggyBac integration. While such a system offers ample recording media necessary for diverse edit accumulation without exhaustion of recording capacity (*i.e.* saturation of editing sites on DNA Typewriter Tape), a major hurdle in the analysis of the DNA Typewriter data is that the Tape barcodes (TapeBCs) are not sufficient to define a unique genomic locus at which lineage information is recorded. This is due to the observation in our study that many or even most piggyBac integrated Tapes are duplicated in the genome, presumably due to excision/reintegration events before the transposase has diluted out.. The presence of duplicated Tapes presents a challenge for reconstructing lineages, as it becomes difficult to compare the patterns of edits observed for a given TapeBC across cells without knowing the underlying Tape locus from which they arose.

However, given that we have simultaneously recorded cell barcode information along with the expressed tape sequences, we can detect these duplication events as coexpressed tapes of differing patterns of edits. In other words, if a given cell has a sequential pattern of edits for a particular TapeBC of A-B-C but also expresses the same TapeBC with a pattern of D-E-F, this TapeBC is associated with at least two genomic integrations of DNA Typewriter Tape. Thus, we set out to devise a method that could infer the most likely structure of integration events per TapeBC into the genome as well as the patterns of edits that were generated from each integration locus. In this way, we can capture the true sequence of events that occur from the starting cell at each locus independently.

### Determining consensus patterns of edits

We start from a table that is populated at each row by a unique sequence consisting of the CellBC, TapeBC, editing sites 1-6 (InsertBC(1)-(6)), and the number of UMIs that contain this exact sequence. Next, we filter this table for a given TapeBC in order to track its patterns of edits across all cells. We then compute a pairwise hamming distance matrix between each of the unique sequences at the edit sites. This distance matrix is then “clustered” by grouping all the unique sequences by exact matches and assigning them to a cluster if the cluster size would be greater than 100 UMIs. The remaining UMIs that had not been assigned to any cluster would be discarded as noise; this is analogous to a DBSCAN approach with a radius of 0. The consensus sequence for each cluster is refined into only the edited portions by searching for the presence of the edit key “GGAT” every 7 bases. This approach offers a few advantages: by pooling information across all cells, we are looking for patterns of edits that occur throughout the dataset. Additionally, the sequences that are not assigned to any cluster are likely generated from PCR or sequencing errors, and can be easily identified and thrown out.

### Generating graph of edit pattern mutual exclusivity

Once the consensus patterns of edits are identified, we can then generate a cell-by-pattern matrix, with the values being the number of UMIs associated with each cell and pattern. This matrix is then row-normalized and log-transformed in order to more clearly accentuate differences between cells. Next, a binarization threshold is determined as the lowest point between the modes of the distribution of normalized values in the cell-by-pattern matrix. Binarizing this matrix allows us to generate a more interpretable matrix which represents whether or not a given pattern of edits is present in that cell for that TapeBC. This binarized matrix represents an intuitive way to think about multiple integrations; if a cell has a row sum of more than 1 (i.e. more than one pattern of edits), there are multiple integrations. However, the goal here is not to infer per-cell integrations, but rather to infer the tape integration structure across all cells. For example, if pattern 1 and pattern 2 are both co-expressed in the same cells, they must have been generated from different loci.

To learn the structure of Tape integrations across all cells for a given TapeBC and assign patterns of edits to specific loci of integration, we performed the following analysis: We clustered this binary matrix using DBSCAN to group cells with the exact same groups of expressed patterns. Cells that were not assigned to any given cluster, usually due to dropout, were assigned to a cluster using a KNN classifier in normalized log-UMI space. Each cluster thus represents a group of cells for which a given pattern is expressed. For each cluster, given its representative binary vector of whether each pattern of edits is expressed, we can then generate a matrix of mutual exclusivity by computing the pairwise cosine distance between this vector and itself. This generates an adjacency matrix which in turn implies a graph whose nodes represent patterns of edits and whose edges represent mutual exclusivity between that pair of patterns in the same cells. The cosine metric was chosen because it only draws an edge between patterns if a 1 is present for both patterns, rather than a 0 in both. We compute such graphs for each cluster of cells and then intersect them all to generate a single representative graph across all cells. The assumption here is that if two patterns are co-expressed in at least one cluster, they must be mutually exclusive with respect to tape locus in all cells. That is to say, those patterns of edits must have been generated at separate loci by definition.

### Grouping patterns into loci

Given this graph, how can we determine how to group patterns into shared loci? If such a graph were complete, that is to say fully connected, it would imply that each pattern of edits is never co-expressed and therefore there is only one locus from which all patterns were generated. We can extend this idea to our observed graphs: a complete subgraph (or clique) within the graph represents a putative locus (or Tape integration event). Therefore, the problem can be formulated in terms of finding a node partition of the graph such that each induced subgraph is a clique. However, we must constrain this problem with several biological considerations. The most likely explanation is one that minimizes the number of inferred integration events; thus, we pick the solution that minimizes the number of cliques. Given that there may be multiple equivalent solutions under this metric, we introduce edge weights as the number of shared consecutive edits between patterns. In this way, cliques with higher sums of edge weights have more related patterns and are therefore more likely to represent patterns that were generated from the same locus. To summarize, the problem becomes finding a clique partition of the graph that: 1) minimizes the number of cliques, and 2) maximizes the sum of edge weights across all cliques.

The problem of finding minimal clique partitions cannot be realistically achieved through exhaustive search on large datasets. However, the problem can be flipped on its head. The complement of the graph, generated by inverting all the edges, can be “colored” with a minimal number of colors using greedy graph coloring approaches. Graph coloring refers to a problem where nodes are assigned colors such that no two neighboring nodes can share the same color. Thus, the color assignments of the graph complement correspond to clique memberships in the original graph. However, since we need to also optimize for the sum of edge weights in the original graph, we run the graph coloring algorithm a large number of times with randomized node orderings and select the solution which produces the fewest number of cliques and the highest sum of edge weights retained.

Each of these inferred cliques then represents a locus of integration for that specific TapeBC, as well as grouping the exact patterns that belong at that locus. From this, we can then generate a final matrix with cells as the rows and inferred loci as the columns. The detected patterns per cell are then stored in the columns according to the clique groupings. This final matrix is then used as the basis for calculating pairwise lineage distances between cells and ultimately for generating phylogenies with UPGMA.

## SUPPLEMENTARY FIGURES

**Figure S1.**
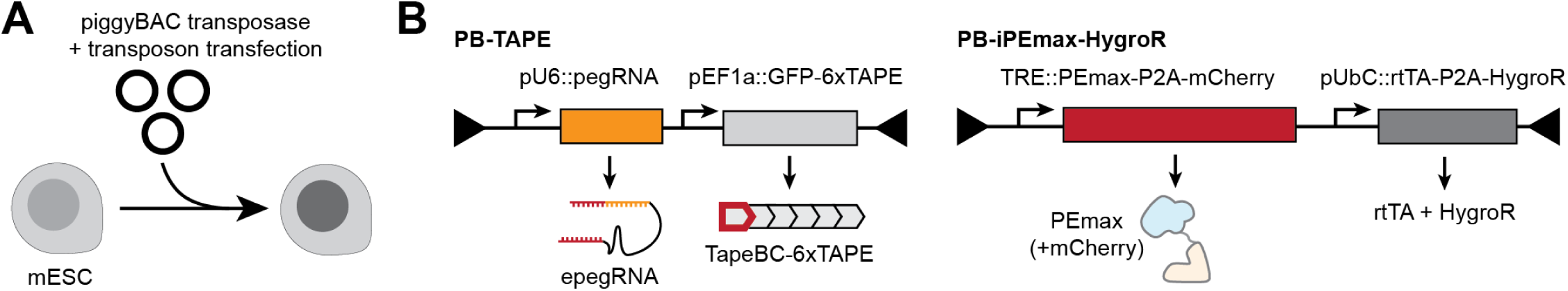
Generation of mESCs with DNA Typewriter Tape. **(A)** Schematic of transfecting mESC with three different kinds of DNA plasmids: transposon bearing DNA Typewriter Tape (PB-TAPE), transposon bearing Dox-inducible Prime Editor expression cassette (PB-iPEmax-HygroR), and plasmid expressing piggyBac transposase. **(B)** Architecture of two transposons: PB-TAPE contains a human U6 promoter driving epegRNA expression (pU6::epegRNA, where epegRNA inserts one of 64 possible NNNGGA sequences to DNA Typewriter Tape targets) and human EF1a promoter driving GFP-6xTAPE expression (pEF1a::GFP-6xTAPE), where 12-bp (NNNNNAANNNNN) TapeBC and 6xTAPE (DNA Typewriter Tape with 6 sequential editing sites) embedded in 3’-UTR of eGFP-encoding transcript. PB-iPEmax-HygroR contains a Dox-inducible promoter driving expression of PEmax-P2A-mCherry (TRE::PEmax-P2A-mCherry) and a human ubiquitin C promoter driving rtTA-P2A-HygroR expression. Dox-induction of modified cells result in expression of PEmax that binds to epegRNA and targets 6xTAPE for random sequential editing to generate a heritable DNA barcode used for lineage tracing and lineage tree reconstruction.

**Figure S2.**
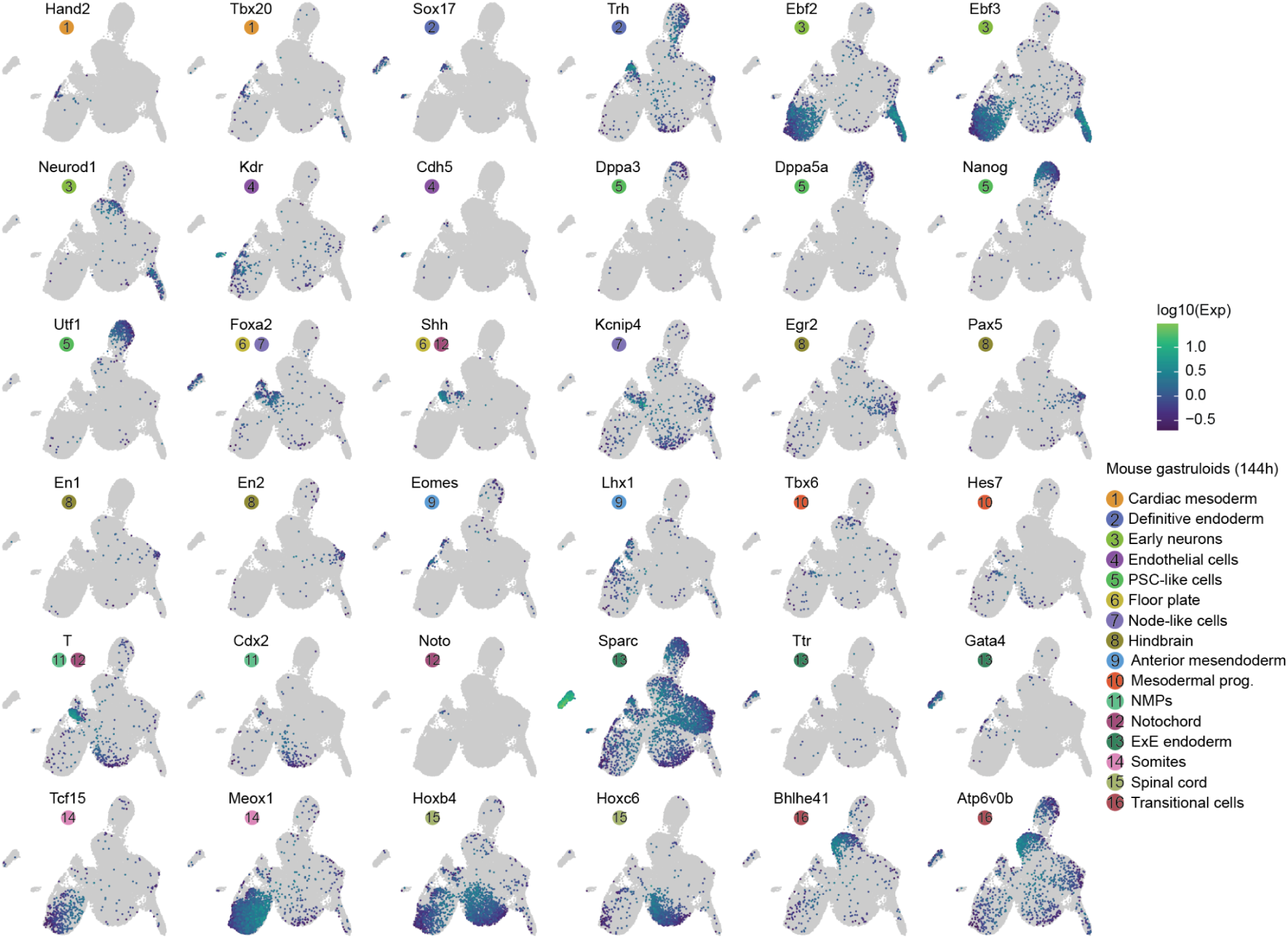
Transcriptional heterogeneity in mouse gastruloids. The same UMAP as in Figure 2B, colored by gene expression of marker genes for each cell cluster. References for marker genes are provided in **Table S1**.

**Figure S3.**
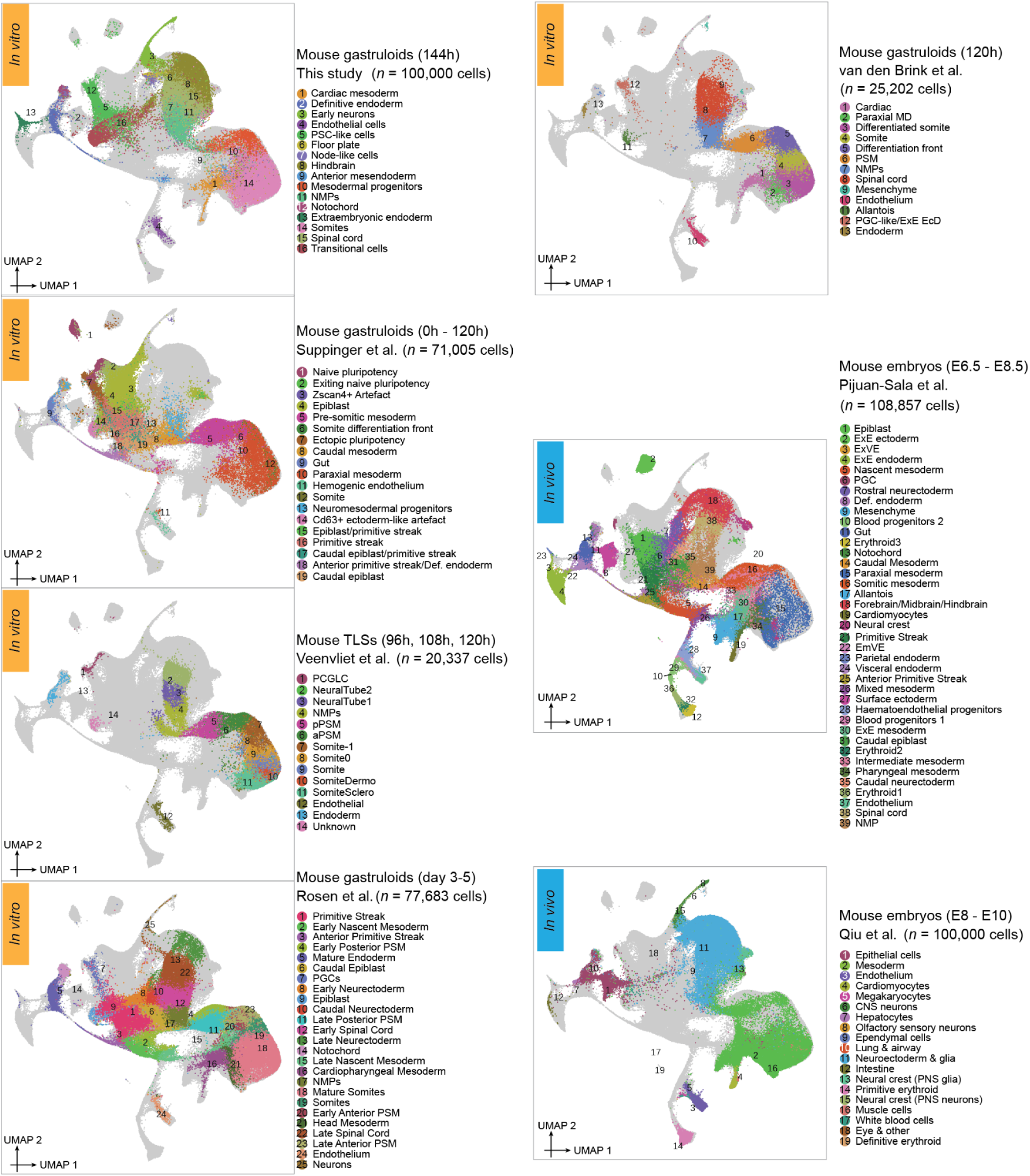
Integrating & co-embedding cells from seven studies of mouse gastruloids or embryos. UMAP visualization of 503,084 co-embedded cells from seven datasets, including mouse gastruloids from this study, four published mouse gastruloid studies^25,29,32,33^, and two studies of mouse embryos during gastrulation (E6.5-E8.5)^31^ or early somitogenesis (E8-E10)^30^, after batch correction^53^. The same UMAP is shown seven times, with colors highlighting cells from each dataset. To align with the cell counts in the other datasets, sci-RNA-seq3 profiles from this study (top left) and mouse embryos early somitogenesis^30^ (bottom right) were randomly subsampled to 100,000 cells each.

**Figure S4.**
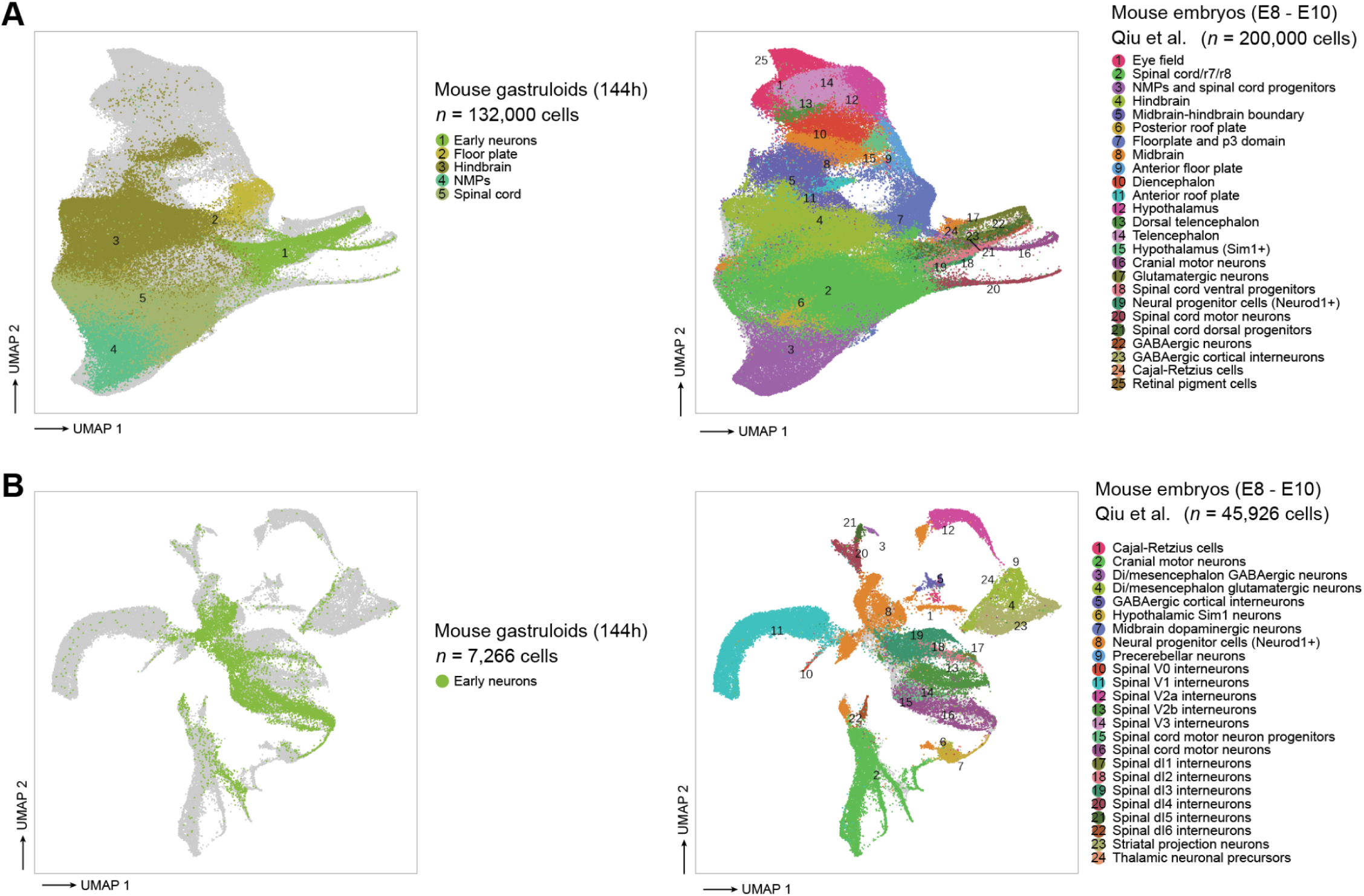
Integrating & co-embedding patterned neuroectodermal and neuronal cells from mouse gastruloids and embryos. **(A)** UMAP visualization of single cell transcriptional profiles of 332,000 nuclei annotated as patterned neuroectodermal cell types from either 144 hr mouse gastruloids (left; this study) or mouse embryos during early somitogenesis (E8-E10)^30^ (right), after batch correction^53^. The same UMAP is shown twice, with colors highlighting cells from each dataset. Cells from the mouse embryo study were randomly subsampled to 200,000. **(B)** Same as panel **A**, but for 53,192 nuclei annotated as neuronal cell types in the same two studies.

**Figure S5.**
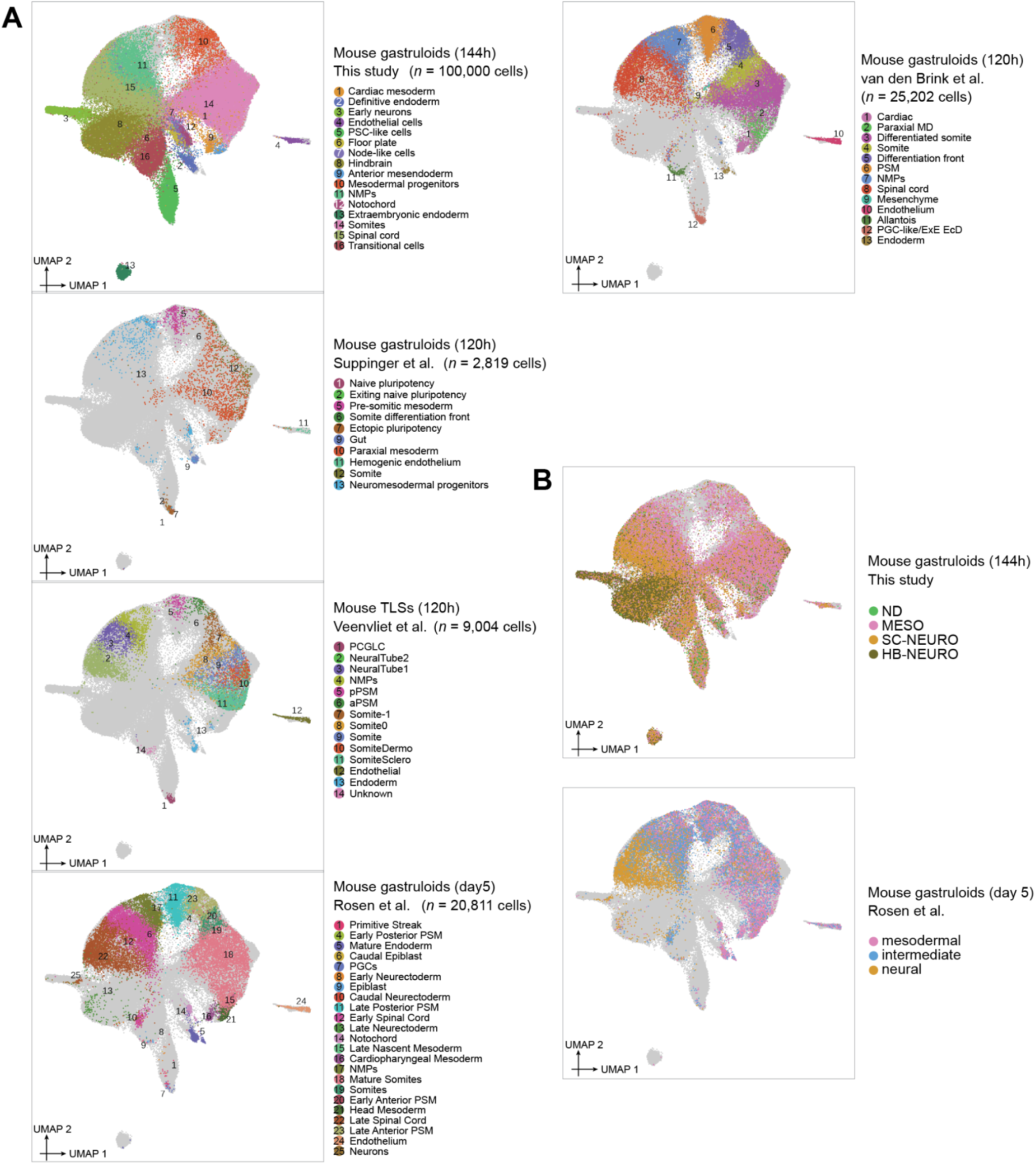
Integrating & co-embedding cells from mouse gastruloids from this study (144 hrs) and four other studies (120 hr time point only). **(A)** UMAP visualization of 157,836 co-embedded cells from five datasets, including mouse gastruloids from this study (144 hours), and four other published mouse gastruloid studies (120 hours)^25,29,32,33^, after batch correction^53^. The same UMAP is shown five times, with colors highlighting cells from each dataset. To align with the cell counts in the other datasets, cells from the current study (top left) were randomly subsampled to 100,000 cells each. **(B)** The same UMAP as shown in panel **A** is displayed twice, with colors highlighting cells from the four gastruloid groups delineated in this study (top), or from the clusters delineated by Rosen et al. (2022)^32^.

**Figure S6.**
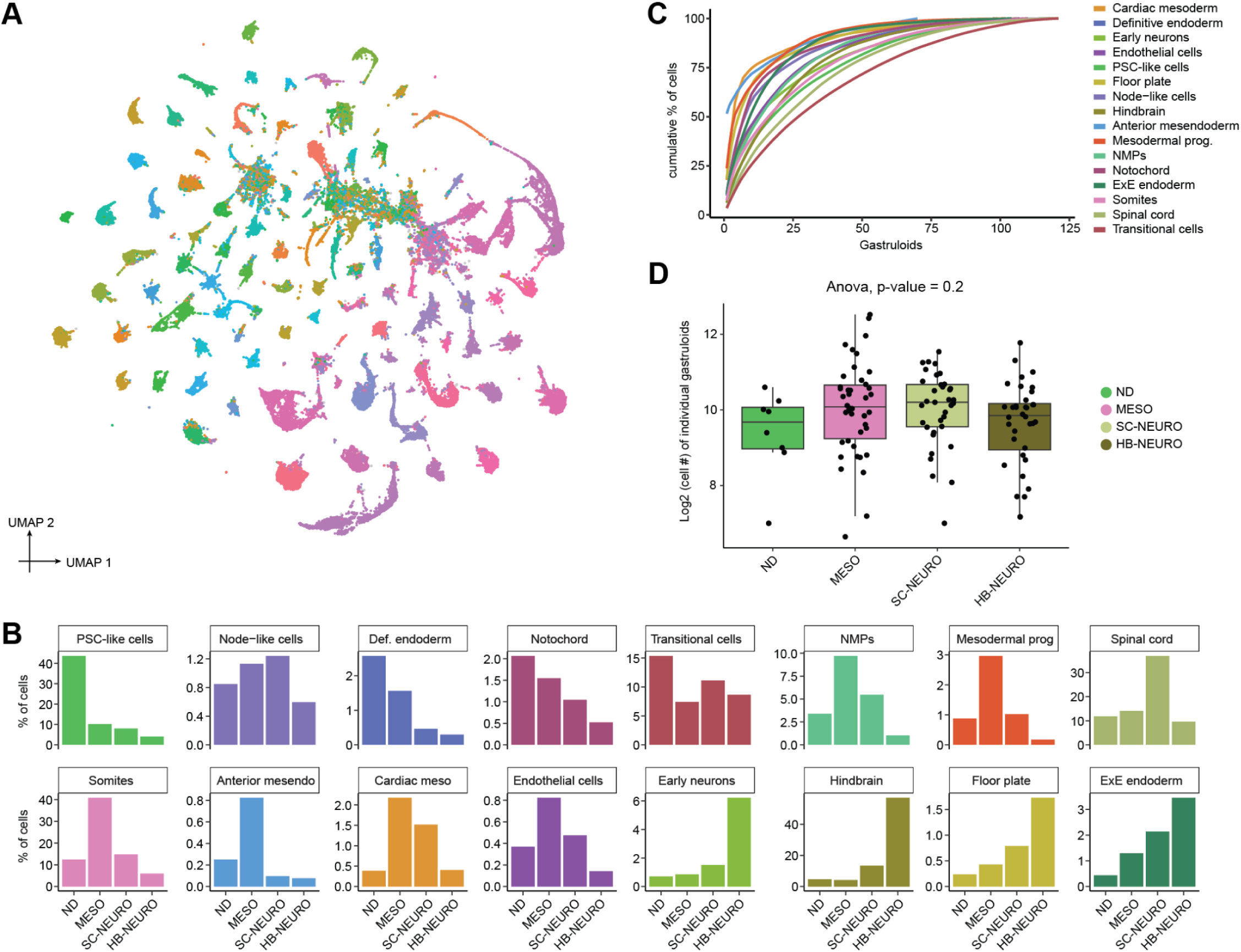
Assigning cells & assessing heterogeneity across 121 monoclonal mouse gastruloids. **(A)** 2D UMAP visualization of 154,988 cells (with ≥3 TapeBCs detected) derived from 121 monoclonal mouse gastruloids. Dimensionality reduction was performed on a TapeBC × cell matrix with read counts as values, using PCA followed by UMAP purely for visualization purposes. Cells are colored according to the gastruloids to which they were assigned. Unassigned cells (4%) are depicted in gray. **(B)** The proportions of sixteen cell types for individual gastruloids were calculated, and the average values within each of the four groups (ND, MESO, SC-NEURO, HB-NEURO) are presented. **(C)** For each cell type, the cumulative percentage of cells was plotted across 121 gastruloids, ranked by the abundance of that cell type from highest to lowest. **(D)** Log2 number of cells assigned to individual gastruloids (n = 121) for each of the four groups. Boxplots represent IQR (25th, 50th, 75th percentile) with whiskers representing 1.5× IQR. ND: non-differentiating, MESO: somite-like, SC-NEURO: spinal cord-like, or HB-NEURO: hindbrain-like.

**Figure S7.**
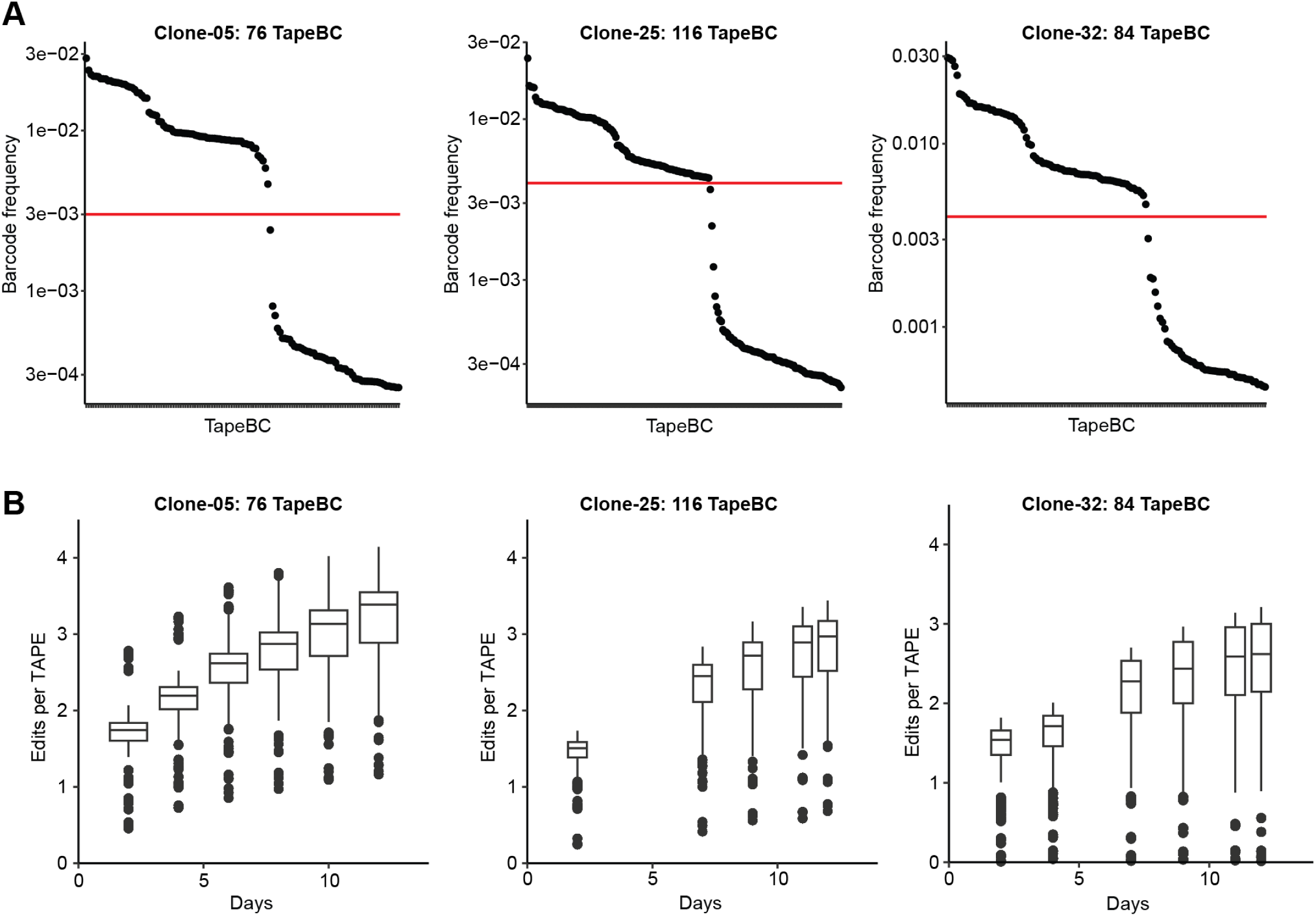
Characterization of three monoclonal mESC cell lines for recording capacity for DNA Typewriter-based lineage recording. **(A)** TapeBC barcodes from each of three monoclonal mESC cell lines were PCR amplified and sequenced from genomic DNA. Frequency thresholds (red line) to define bona fide TapeBC barcodes were set as shown, resulting in estimates of 76, 116, and 84 DTT/epegRNA integrations for Clone-05, Clone-25, and Clone-32, respectively. Note that these are lower bounds, as some integrants are duplicated by piggyBac excision and re-integration events^28,34^**. (B)** The accumulation of editing at DTT in each monoclonal mESC cell line with continuous doxycycline (Dox) treatment to induce PEmax expression. 100 ng/mL Dox was added to cells at Day 0, and cell lines were collected at multiple time points of 2-3 days in interval, depending on the availability of cells at the time of the passaging and collection. The experiment was terminated at Day 12, which is equivalent to the last time point in the monoclonal gastruloid induction protocol. Boxplot represents IQR (25th, 50th, 75th percentile) with whiskers representing 1.5× IQR.

**Figure S8.**
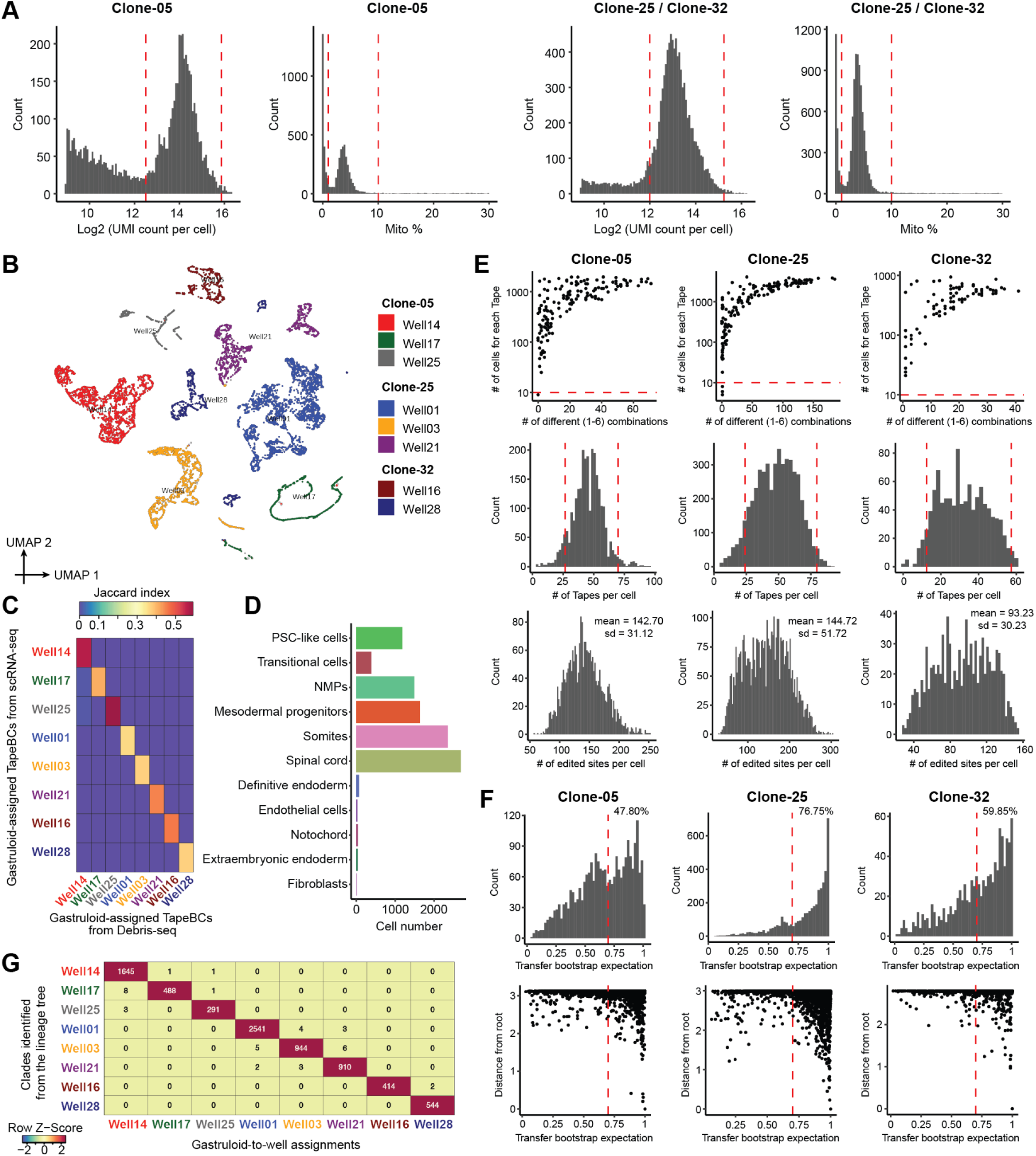
Quality control for scRNA-seq and cell-to-gastruloid assignments. **(A)** Histograms of log2(UMI count) per cell and the proportion of reads mapping to mitochondrial chromosomes (Mito%) per cell, derived from scRNA-seq data (10X Genomics) from Lane 1 (left, Clone-05) and Lane 2 (right, Clones-25 & 32). Cells with log2(UMI counts) <12.5 in Lane 1 or <12 in Lane 2 were excluded, as were those in the top 0.5% of total UMI counts (red vertical lines) and those with Mito% >10% or <1% (red vertical lines). **(B)** 2D UMAP visualization of 8,547 cells, with ≥2 TapeBCs detected and TapeBC UMIs ≥10. Dimensionality reduction was performed with PCA on a TapeBC × cell matrix with read counts as values, followed by UMAP solely for visualization. Cells are colored by their assignment to one of eight monoclonal gastruloids. Unassigned cells (2%) are depicted in light gray. **(C)** Jaccard similarities between the TapeBCs detected in cells assigned to each gastruloid (row; detected in ≥5% of the cells within that gastruloid) vs. TapeBCs detected by Debris-seq of each well (column). Only TapeBCs detected in both scRNA-seq and Debris-seq were included in these calculations. **(D)** Frequency of each cell type across 9,929 cells. **(E)** Filtering based on DTTs detected from Clone-05 (left), Clone-25 (middle), and Clone-32 (right): Top: The number of cells detected for each DTT (y-axis) is plotted against the number of different InsertBC(1-6) combinations (x-axis). DTTs with a cell count >10 (red horizontal lines) were retained. Middle: Histograms of the numbers of post-filtering DTTs detected per cell. Cells outside the range of mean - 1.5 × SD to mean + 2 × SD were excluded (red vertical lines). Bottom: Post-filtering histograms of the total number of edited sites per cell, with the mean and standard deviation indicated. **(F)** We performed a bootstrapping analysis to evaluate the robustness of the trees shown in Figure 3D, composed of 2,438 tips & 2,437 clades in Clone-05, 4,418 tips & 4,417 clades in Clone-25, and 960 tips & 959 clades in Clone-32. For each of 100 bootstraps, an equal number of DTTs (each containing six edit sites) was resampled with replacement to reconstruct a new tree. Clade concordance was determined by comparing each bootstrap tree to the original and calculating transfer bootstrap expectation (TBE)^35^. Briefly, the TBE of each clade in the original tree was calculated by determining the minimal transfer distance across all clades in the bootstrap tree, followed by computing the mean values over the 100 bootstraps. Here we show a histogram of the TBEs of clades (top), and a scatter plot of TBE against the distance from the root for each clade (bottom). Dashed red vertical lines indicate a TBE cutoff of 0.7, with values greater than that corresponding to moderate to strong support^35^. **(G)** The numbers of cells in each of ten major clades (with two clades combined for Well17 and Well01, respectively) of the cell lineage tree shown in Figure 3D plotted against the well assignments of those same cells.

**Figure S9.**
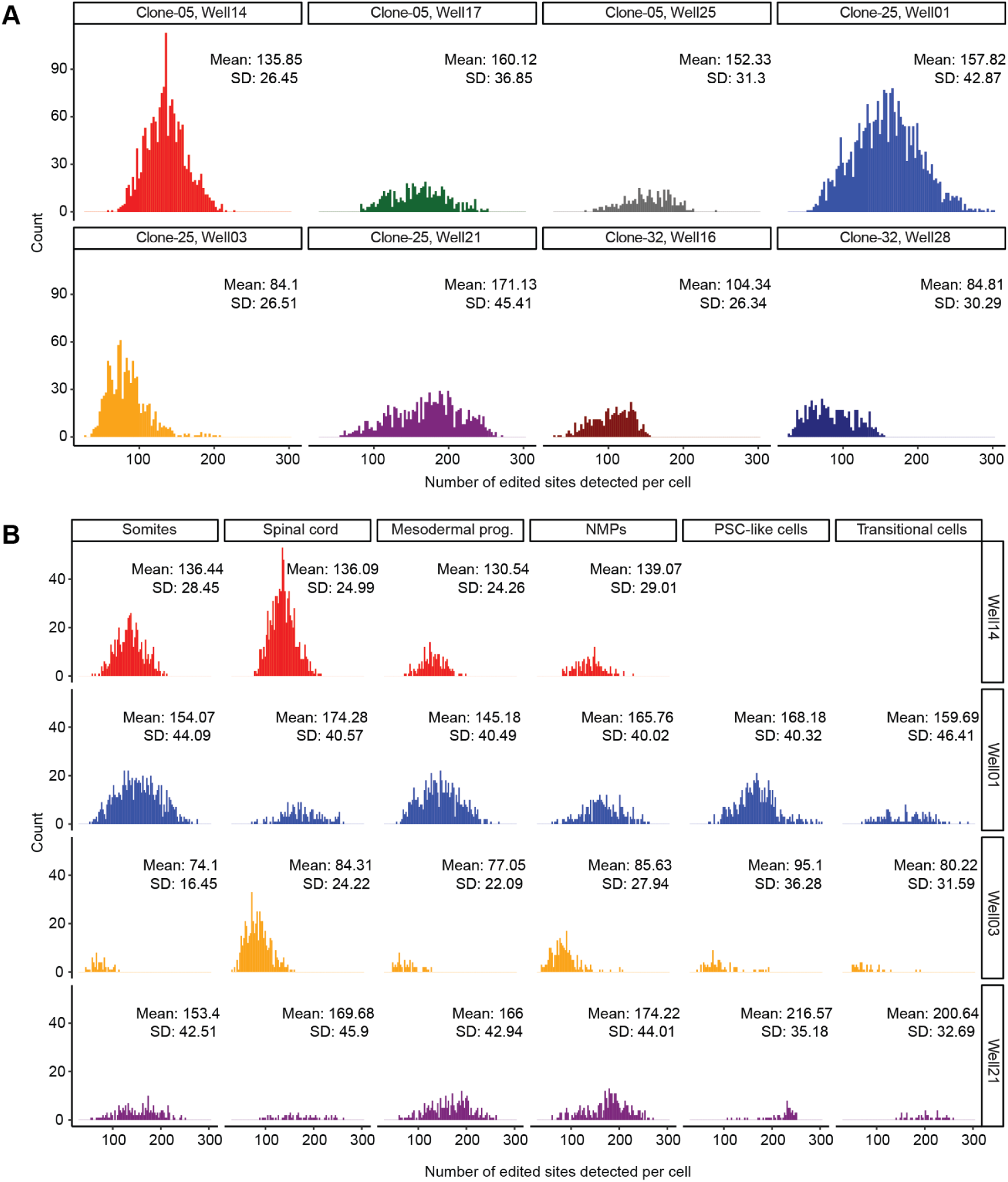
Numbers of edited sites in DTT detected per cell, across gastruloids and cell types. **(A)** To systematically compare the overall activity of PEmax between gastruloids and cell lines, for each DTT observed in each of the 7,816 cells used to reconstruct the lineage tree, we counted the total number of edited sites observed in each cell. Histograms of these counts are shown, together with the mean and standard deviation, for each of the eight gastruloids. **(B)** Same as panel **A**, but for the four gastruloids with the most sampled cells, and further splitting these out by the six most abundant cell types.

**Figure S10.**
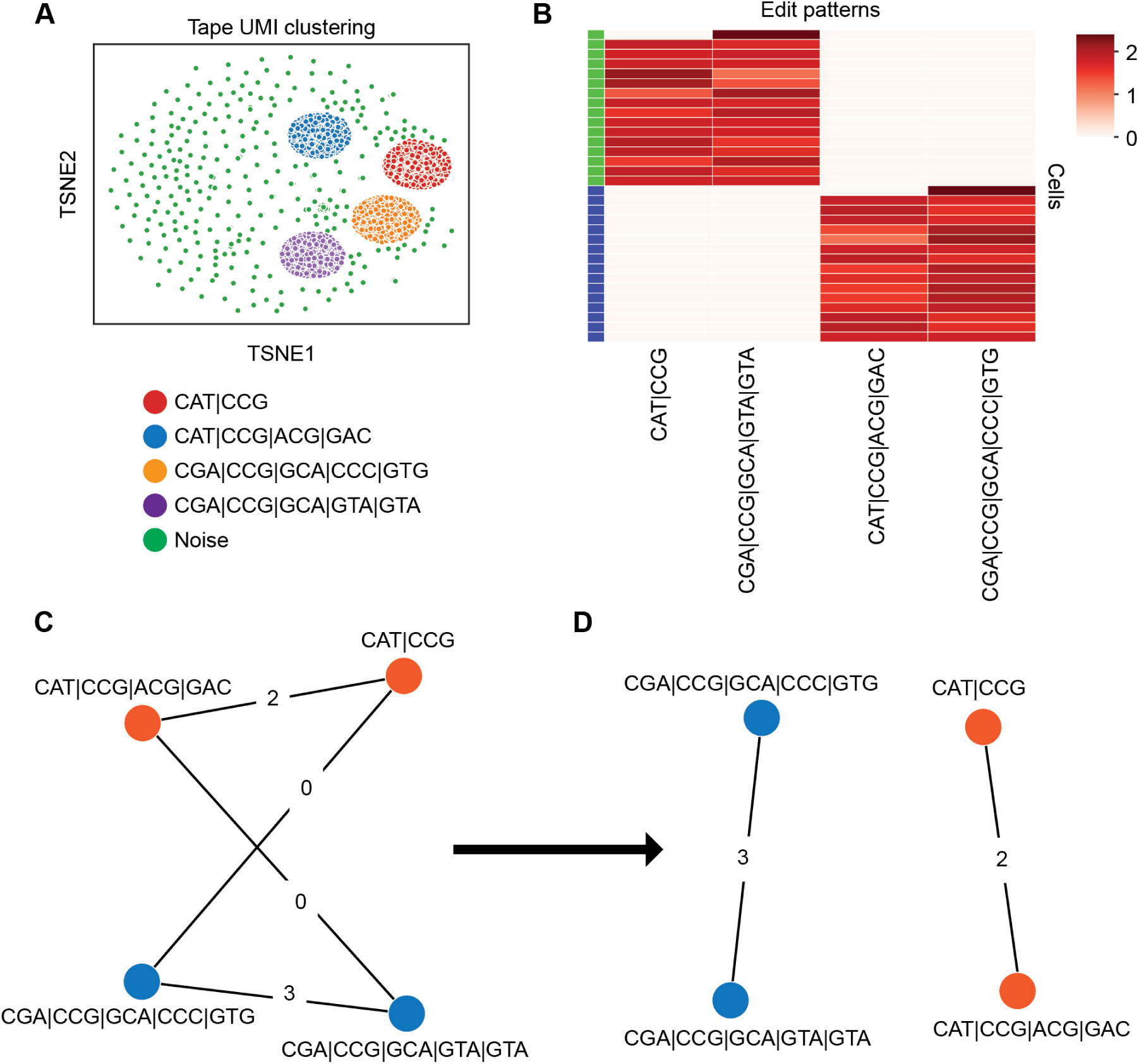
Disentangling TapeBCs that are integrated at multiple genomic locations. We relied on piggyBac transposition at a high MOI in order to obtain a large number of DTTs in each monoclonal cell line. Each DTT is associated with a degenerate TapeBC that facilitates the assignment of DTT-derived sequencing reads to specific integrations (**Figure S1B**). However, a major challenge is that due to piggyBac excision and re-integration events^28,34^ during the establishment of the monoclonal cell line, a given TapeBC might be associated with two or more independent integrations, confounding lineage reconstruction. Here we present an example vignette of our strategy for disentangling such duplicated TapeBCs by relying on the patterns in which their associated edits either <u>do</u> (consistent with deriving from separate integrations) or <u>don’t</u> (consistent with deriving from the same integration) appear in the same cells. Further details on our algorithmic strategy are presented in a **Supplementary Note**. This example vignette is for a specific TapeBC from a subset of 32 cells with one major bifurcation into two clades. **(A)** t-SNE depicting clusters of DTT UMIs across all 32 cells. The labels for each cluster represent a unique pattern of edits. Each 3N edit is separated by a “|”. UMIs not assigned to any cluster were discarded as likely noise from PCR or sequencing errors. **(B)** After identifying consensus edit patterns, we generated a cell-by-pattern matrix with UMI counts, row-normalized and log-transformed to highlight differences between cells. This matrix was then binarized (not shown) and the rows were clustered into two groups of cells. **(C)** The representative mutual exclusivity graph for this dataset of patterns is shown. An edge is drawn between two patterns of edits if and only if those two patterns of edits are never co-expressed in any cluster of cells in the previous matrix. Edge weights indicate the number of shared sequential edits between mutually exclusive patterns. **(D)** Graph coloring on the complement of the mutual exclusivity graph reveals the two most likely groups of mutually exclusive patterns. These groupings correspond to unique DTT integration loci of the same TapeBC.

**Figure S11.**
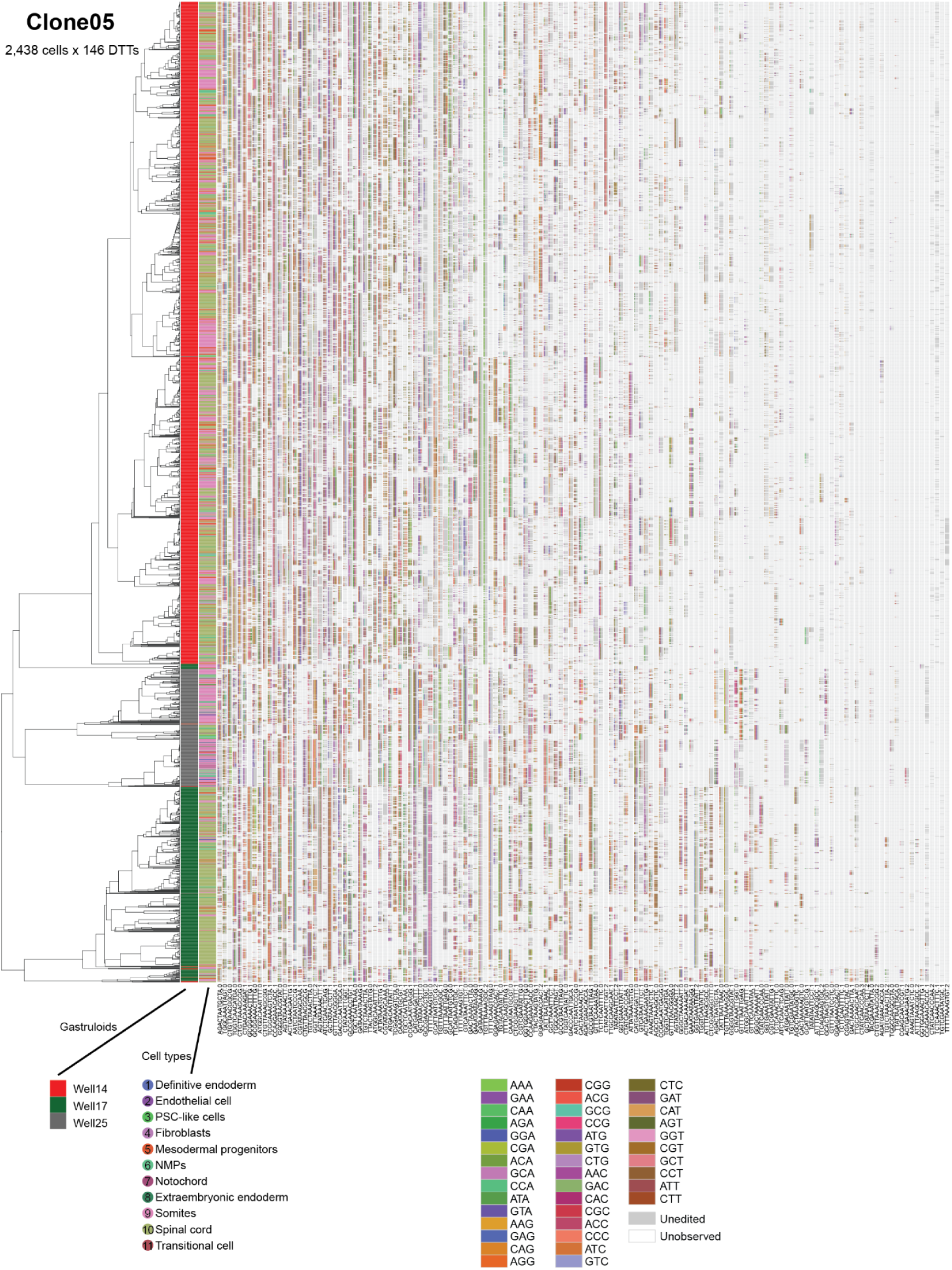
A cell-by-DTT matrix of Clone-05. Left: The left dendrogram shows the phylogenetic tree of 2,438 cells from Clone-05 based on cell-cell distances, the colors of the middle bar correspond to 3 gastruloids to which each cell was assigned, and the colors of the right bar correspond to the 11 annotated cell types. **Right:** Edits at each site across 146 DTTs, ordered by total number of edited sites from high (left) to low (right). Observed, unedited sites are colored grey, while unobserved DTTs are colored white.

**Figure S12.**
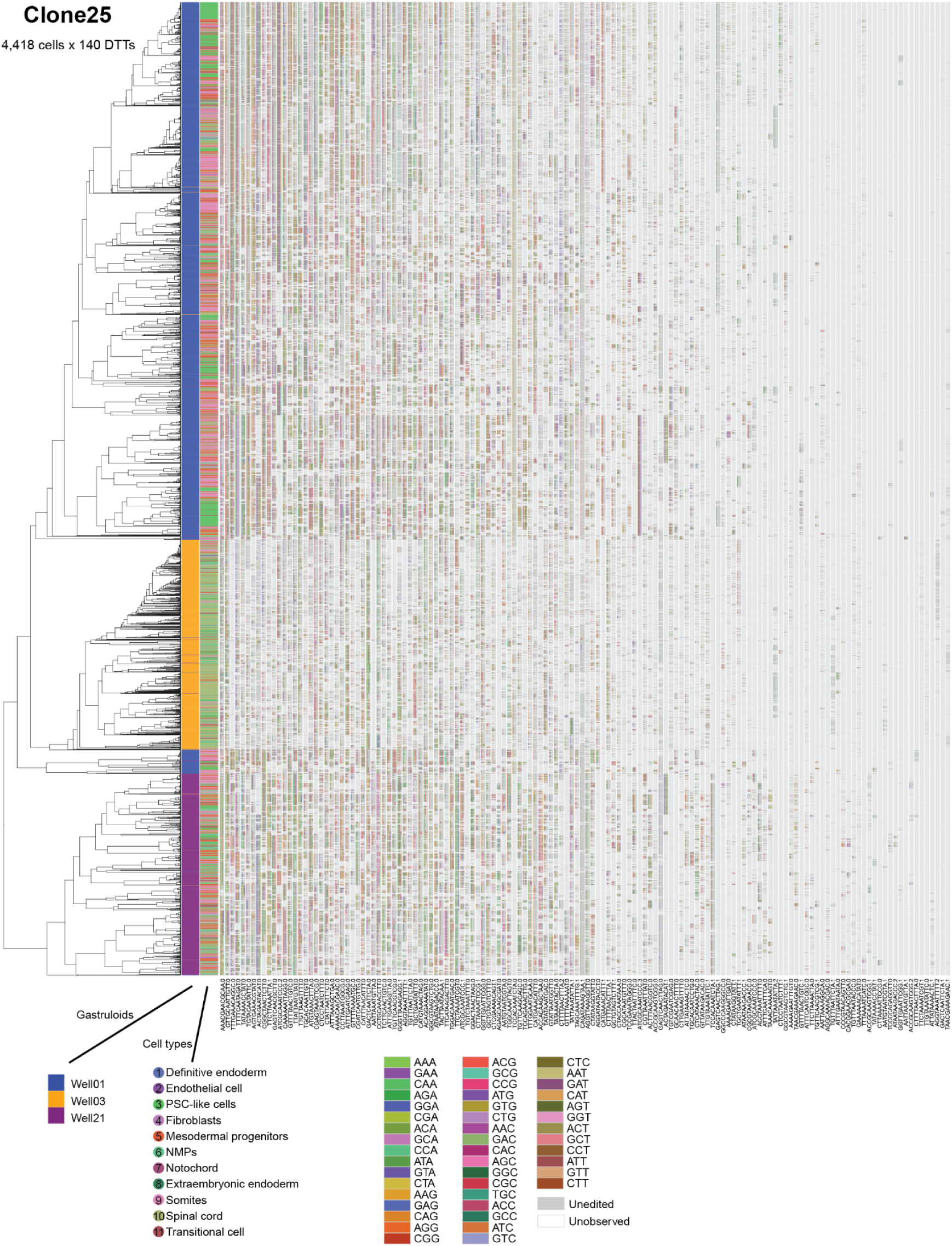
A cell-by-DTT matrix of Clone-25. Left: The left dendrogram shows the phylogenetic tree of 4,418 cells from Clone-25 based on cell-cell distances, the colors of the middle bar correspond to 3 gastruloids to which each cell was assigned, and the colors of the right bar correspond to the 11 annotated cell types. **Right:** Edits at each site across 140 DTTs, ordered by total number of edited sites from high (left) to low (right). Observed, unedited sites are colored grey, while unobserved DTTs are colored white.

**Figure S13.**
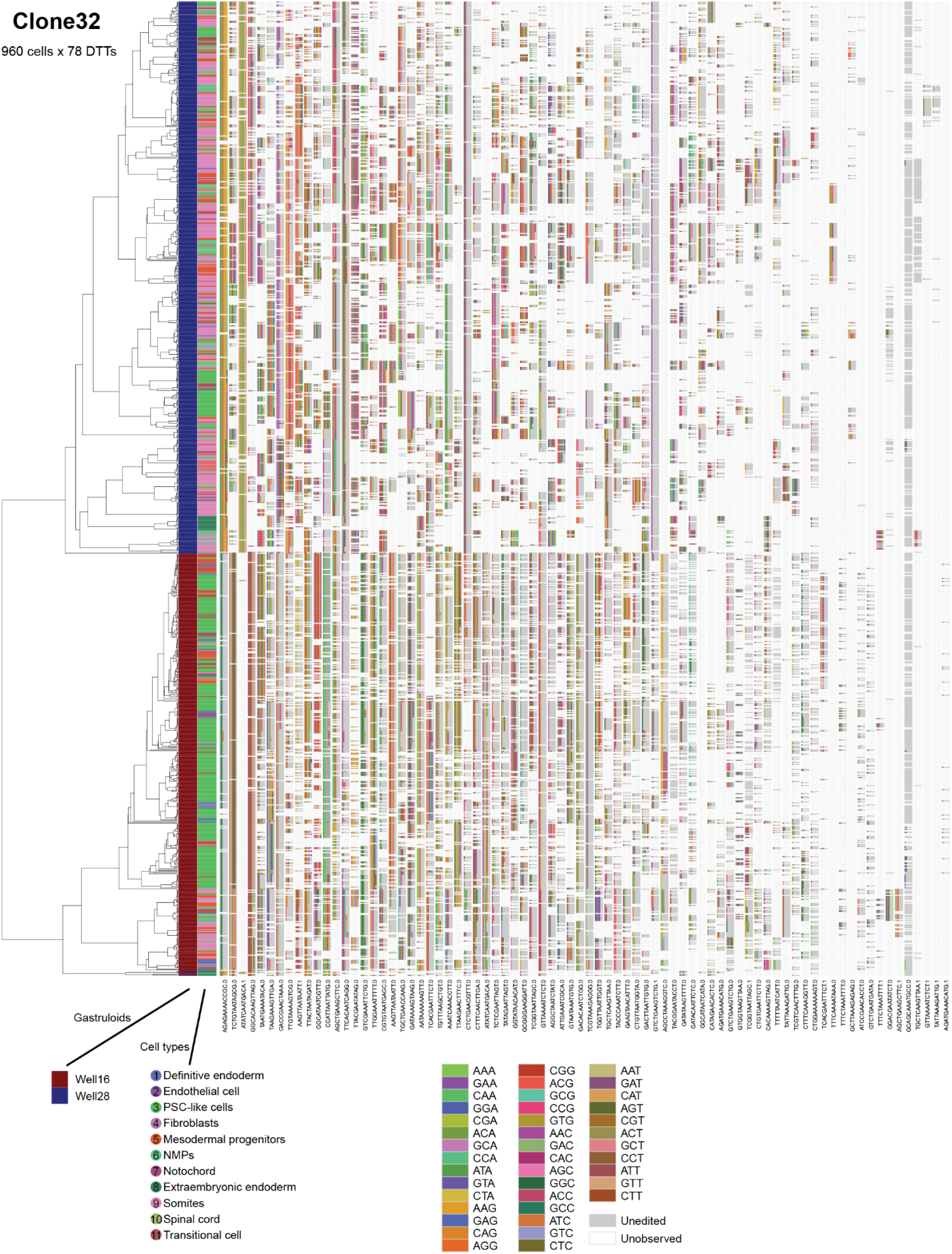
A cell-by-DTT matrix of Clone-32. Left: The left dendrogram shows the phylogenetic tree of 960 cells from Clone-32 based on cell-cell distances, the colors of the middle bar correspond to two gastruloids to which each cell was assigned, and the colors of the right bar correspond to the 11 annotated cell types. **Right:** Edits at each site across 78 DTTs, ordered by total number of edited sites from high (left) to low (right). Observed, unedited sites are colored grey, while unobserved DTTs are colored white.

**Figure S14.**
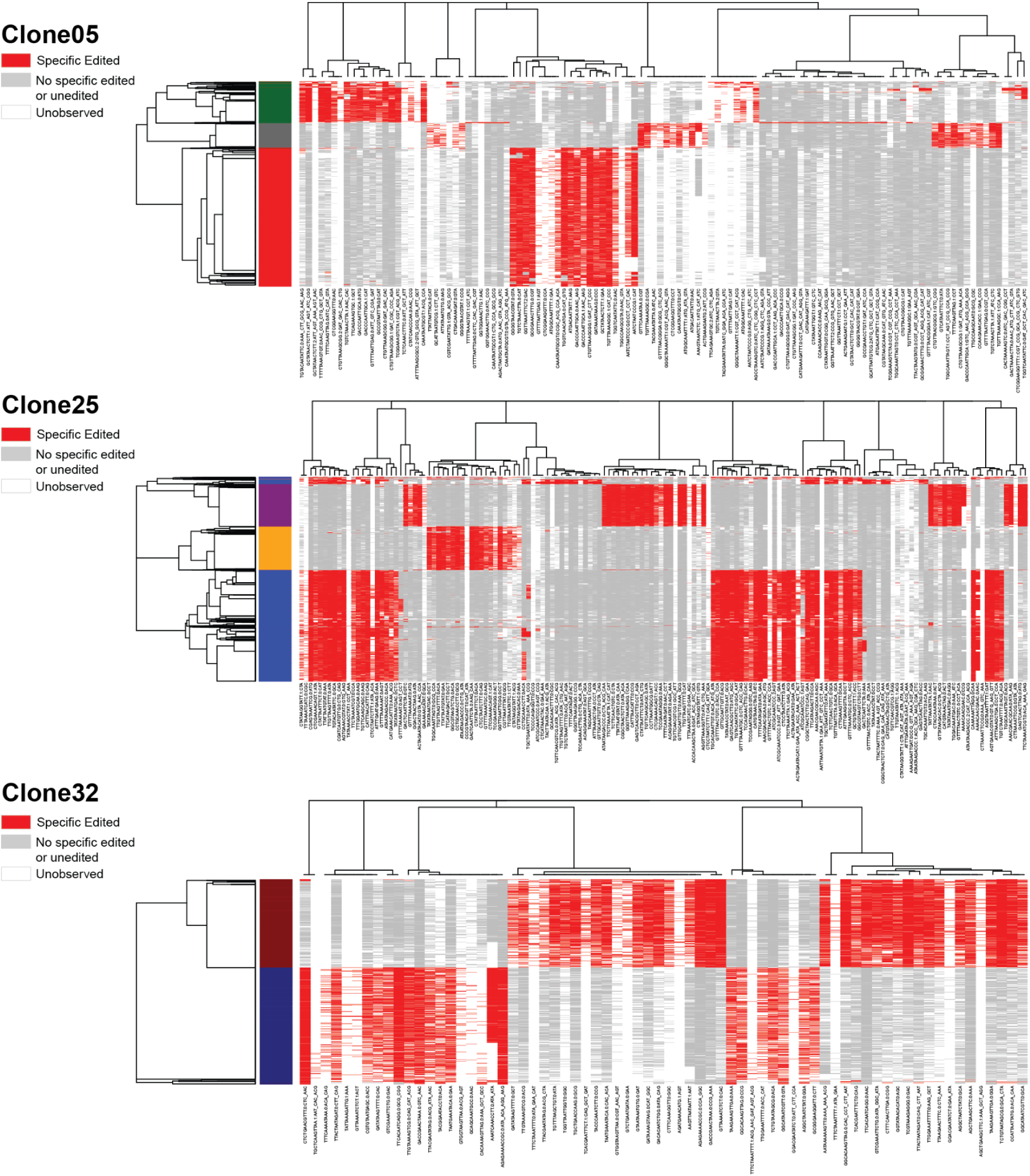
Each monoclonal gastruloid is defined by edits that likely occurred prior to the seeding of its founding mESC. As shown in Figure 3A, we induced PEmax by applying 100 ng/mL doxycycline beginning 24 hrs prior to sorting and seeding of single cells to individual wells. Here we show a biclustered heatmap of 114 (Clone-05), 154 (Clone-25), and 70 (Clone-32) TapeBC-Site-SpecificEdit combinations (columns), after excluding unobserved DTTs, that were detected in >95% of cells assigned to one of the ten major clades shown in Figure 3D, and <5% of cells of clades dominated by a different gastruloid from the same clone (rows). We detect 20 to 57 such TapeBC-Site-SpecificEdit combinations per major clade, which we infer to be editing events that occurred during the 24 hr window during which PEmax was induced but prior to seeding of single cells to individual wells.

**Figure S15.**
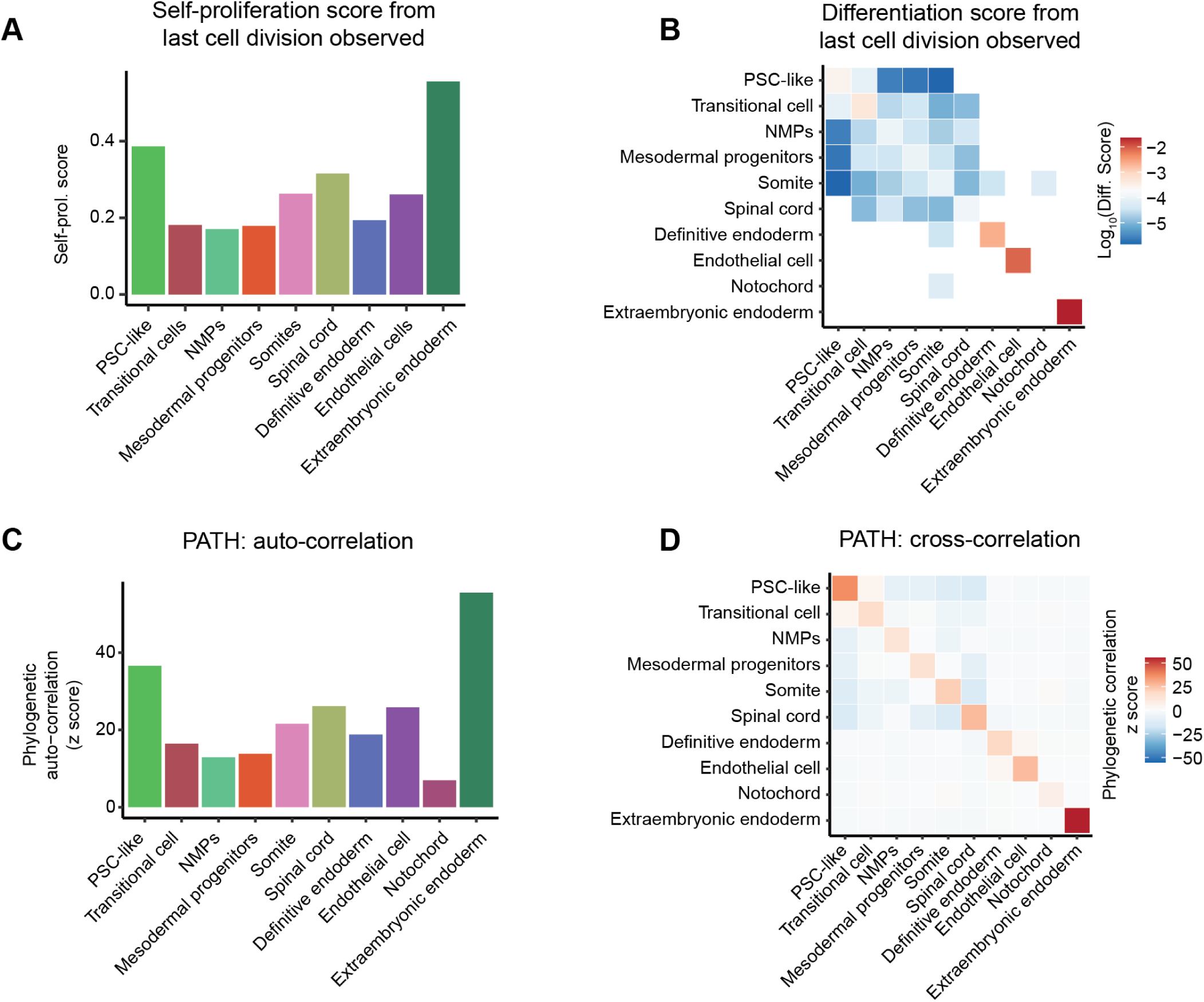
Quantitative assessment of cell-type self-proliferation, differentiation, and correlation in developmental lineages. The cell-cell distance matrices for the three clones were first combined by manually assigning the maximum distance between cells from different clones. **(A)** The self-proliferation score for each cell type was calculated as the ratio of the frequency of both cells from the last observed cell division event belonging to that cell type to the total number of cells of that type. **(B)** The differentiation score for each possible cell-type-pair was calculated as the ratio of the frequency of the pair detected from the last observed cell division event to the total frequency of all possible cell-cell combinations in the phylogenetic tree. **(C)** The auto-correlation z-score of each cell-type was calculated using PATH^37^. **(D)** The cross-correlation z-score of each possible cell-type-pair was calculated using PATH^37^.

**Figure S16.**
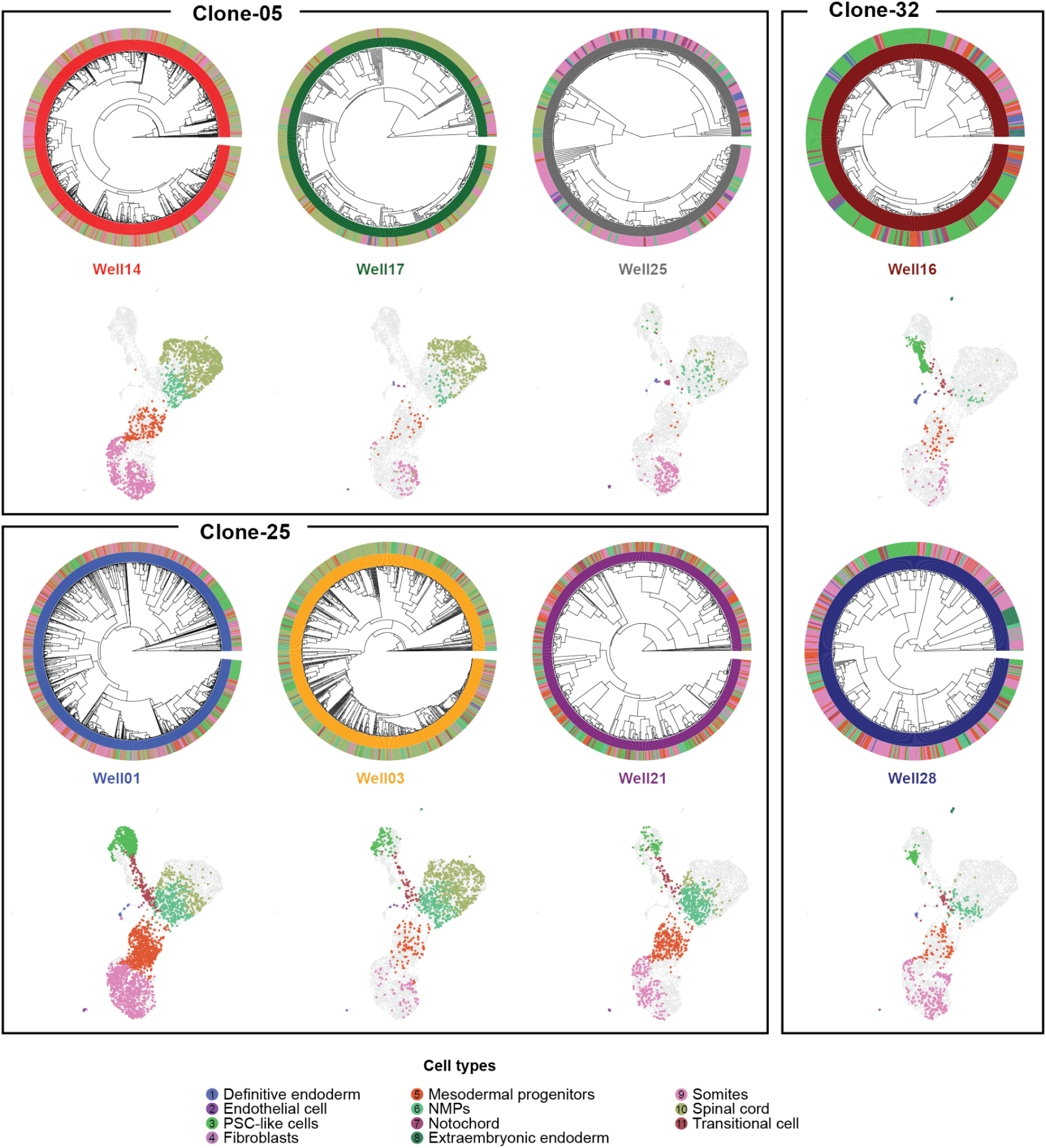
Cell lineage reconstructions of individual monoclonal gastruloids. For each monoclonal gastruloid: Top: Phylogenetic reconstruction of cell lineage relationships among higher-quality, gastruloid-assigned cells for each gastruloid. The inner circle shows the phylogenetic tree based on cell-cell edit distances, the colors of the middle circle correspond to 8 gastruloids to which each cell was assigned, and the colors of the outer circle correspond to the 11 annotated cell types. Bottom: 2D UMAP, reproduced from Figure 3B, showing cells assigned to individual gastruloids, colored by cell types. Gastruloids are grouped by their originating clonal cell line (*i.e.* Clone-05, Clone-25, Clone-32).

**Figure S17.**
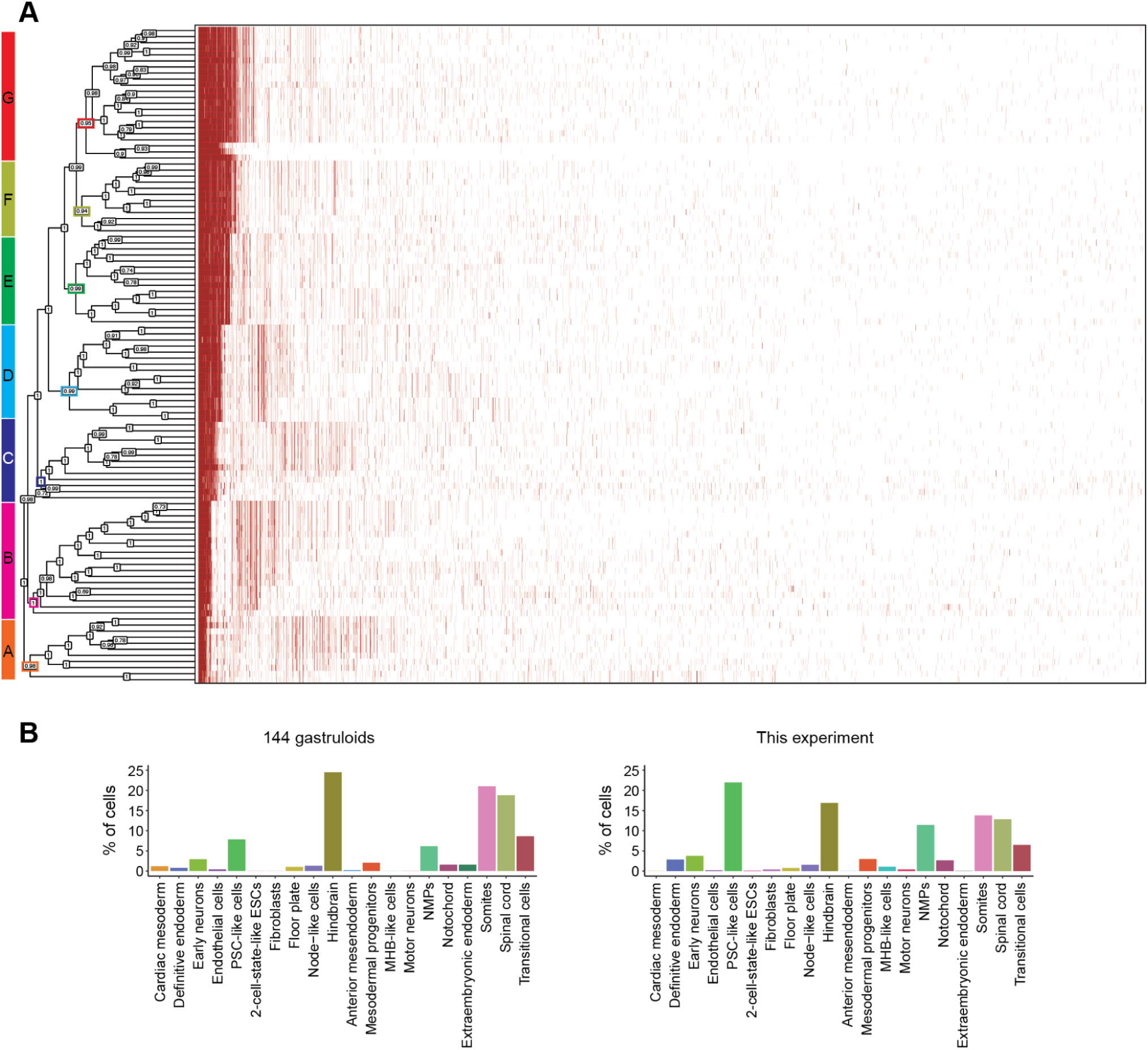
Phylogenetic tree of 108 gastruloids constructed using Debris-seq data. **(A)** For each gastruloid, the percentage of reads corresponding to specific edits at six sites across 76 DTTs was determined. Combinations with at least 20% of reads were identified as dominant "TapeBC-Site-Edit" combinations. The Jaccard similarity between the sets of dominant combinations was calculated, and (1 - similarity) was used as the distance metric across gastruloids. The left phylogenetic tree displays the relationships among the 108 gastruloids, based on 7,146 "TapeBC-Site-Edit" combinations shown on the right (red indicates detection in the corresponding gastruloid). The tree also includes the TBE for each ancestral node, calculated from 100 bootstraps. 106 of the 107 ancestor nodes (99%) showed moderate to strong support, with a TBE greater than 70%. The seven major clades (A-G) are highlighted on the left. **(B)** Comparison of the cell type compositions from our earlier experiment involving 144 gastruloids (247,064 cells; Figure 2B) and this experiment with 108 gastruloids (66,978 cells; Figure 5C).

**Figure S18.**
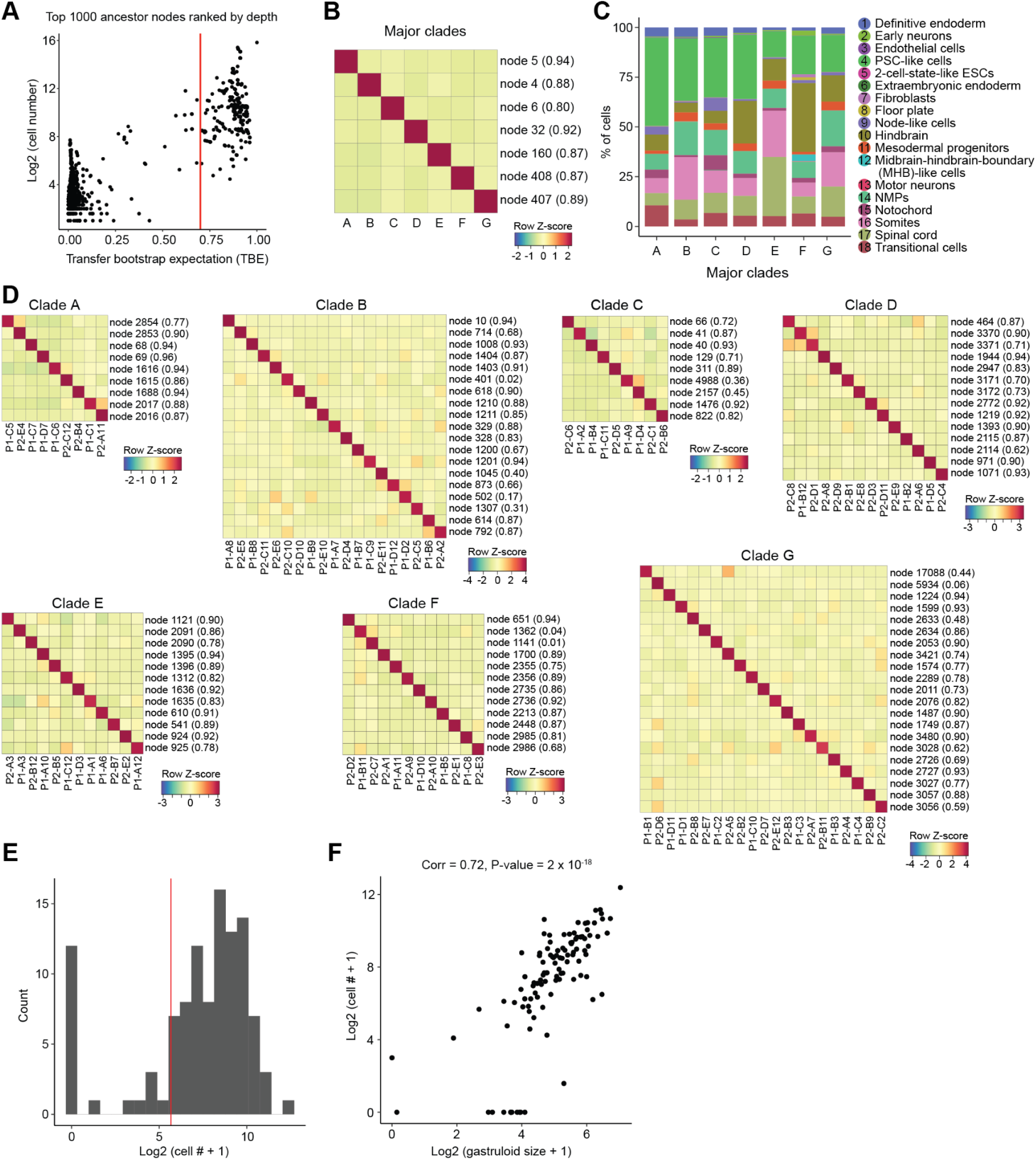
Identifying gastruloid founders among pseudo-ancestors. **(A)** We performed a bootstrapping analysis to evaluate the robustness of the top 1,000 clades (ranked by their distance to the root) in the tree shown in Figure 5D. For each of 100 bootstraps, an equal number of DTTs (each containing six edit sites) was resampled with replacement to reconstruct a new tree. Clade concordance was determined by comparing each bootstrap tree to the original and calculating transfer bootstrap expectation (TBE)^35^. Here we show a scatter plot of TBE against the number of cells for each clade. Dashed red vertical line indicates a TBE cutoff of 0.7, with 45% of clades having a TBE > 0.7. **(B)** To assign each of the seven major clades from the “tree of trees” (Figure 5B) to a pseudo-ancestor in the single-cell phylogenetic tree (Figure 5D), we constructed a cell × “TapeBC-Site-Edit” combination matrix (where 1 indicates the combination was detected in the cell and 0 was not) and normalized it by the sum of each column. For each major clade in the “tree of trees”, we identified its exclusively dominant “TapeBC-Site-Edit” combinations. For each pseudo-ancestor in the single-cell phylogenetic tree, we selected the corresponding subset of cells and calculated the fold-change between the mean value of dominant combinations for a given major clade and the mean value of dominant combinations across the other six major clades. For each major clade, we selected all pseudo-ancestors with a ratio of log2(fold-change + 1) between the given clade and the top one among the left six clades >2, and the one with the highest number of cells was designated as the representative of the major clade. The heatmap represents the log2(fold-change + 1) for each of the seven major clades and their assigned pseudo-ancestors, with the TBE for each node based on 100 bootstraps indicated in the brackets. **(C)** After assignment, cell type composition for each major clade. **(D)** For each of the seven major clades, we subset the cells from its assigned pseudo-ancestor and the gastruloids within that clade, then repeated the assignment. For each gastruloid, we identified its exclusively dominant “TapeBC-Site-Edit” combinations. Then, for each pseudo-ancestor, we calculated the fold-change between the mean value of dominant combinations for a given gastruloid and the mean across other gastruloids. For each gastruloid, we selected all pseudo-ancestors with a ratio of log2(fold-change + 1) between the given gastruloid and the top one among the left gastruloids >2, and the one with the highest number of cells was designated as the representative of the gastruloids. During this process, a total of 96 gastruloids were assigned distinct pseudo-ancestors, comprising 51,963 cells. One gastruloid was excluded due to the absence of exclusive dominant combinations, and 11 were excluded because they could not be assigned to a distinct pseudo-ancestor. The heatmap represents the log2 (fold-change + 1) for each gastruloid and their assigned pseudo-ancestors, with the TBE for each node based on 100 bootstraps indicated in the brackets. **(E)** A total of 51,963 cells were assigned to 108 wells/gastruloids, with 88 of the 108 wells/gastruloids having at least 50 assigned cells (indicated by the red vertical line). **(F)** The correlation between the number of cells assigned to individual wells/gastruloids (y-axis) and gastruloid size (x-axis) across all 108 wells/gastruloids. Pearson correlation coefficients and p-values are indicated above the plot. Sizes of individual gastruloids were measured in units of pixels^2 from their brightfield images. A threshold was applied to each image to distinguish the gastruloid from the background.

**Figure S19.**
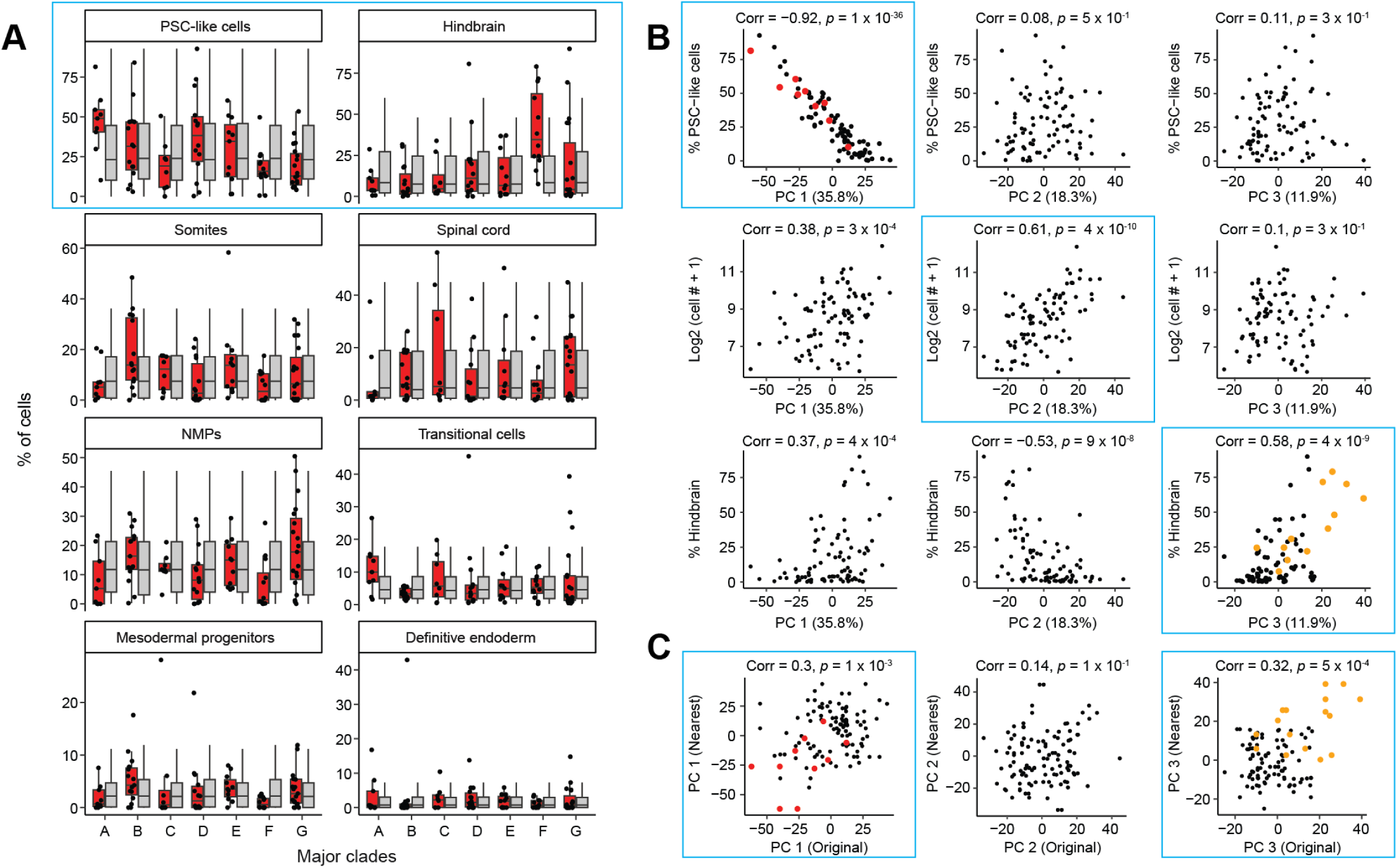
Cell type proportions and PCA analysis of monoclonal gastruloids. **(A)** The proportions of the eight most frequent cell types for 88 individual gastruloids (≥50 cells assigned) were calculated, categorized into seven groups corresponding to the major clades, as shown in Figure 5B. The group labels for each gastruloid were permuted 5,000 times, and cell-type proportions were recalculated each time. Boxplots, showing 88 replicates from real observations (red box) and 5,000 × 88 replicates from permutation (grey box) in each subpanel, represent the interquartile range (25th, 50th, 75th percentiles), with whiskers extending to 1.5× the IQR. Black dots represent replicates from real observations. **(B)** Embeddings of pseudo-bulk RNA-seq profiles of 88 monoclonal gastruloids (≥50 cells assigned) were generated by aggregating single-nucleus data and performing PCA for dimensionality reduction. The percentage of cells corresponding to selected cell types (1st and 3rd rows), or the cell number (2nd row) is plotted along PC1 (left column), PC2 (middle column), and PC3 (right column). In the top left and bottom right subpanels, gastruloids from major clade A (red) and clade F (orange) were highlighted. **(C)** For each of the 88 gastruloids, their nearest neighbors were identified from the phylogenetic lineage tree, resulting in 112 pairs. Of note, some gastruloids had more than one nearest neighbor, and some pairs were redundant (both A vs. B and B vs. A were retained). For each of the top three PCs, the values for each gastruloid (x-axis) are plotted against those of their nearest neighbors (y-axis). Pearson correlation coefficients and p-values are indicated above each plot. Gastruloids from major clade A (red) and clade F (orange) are colored. Of note, subpanels highlighted with a light blue rectangle are shown in Figure 5E-G as well.

**Figure S20.**
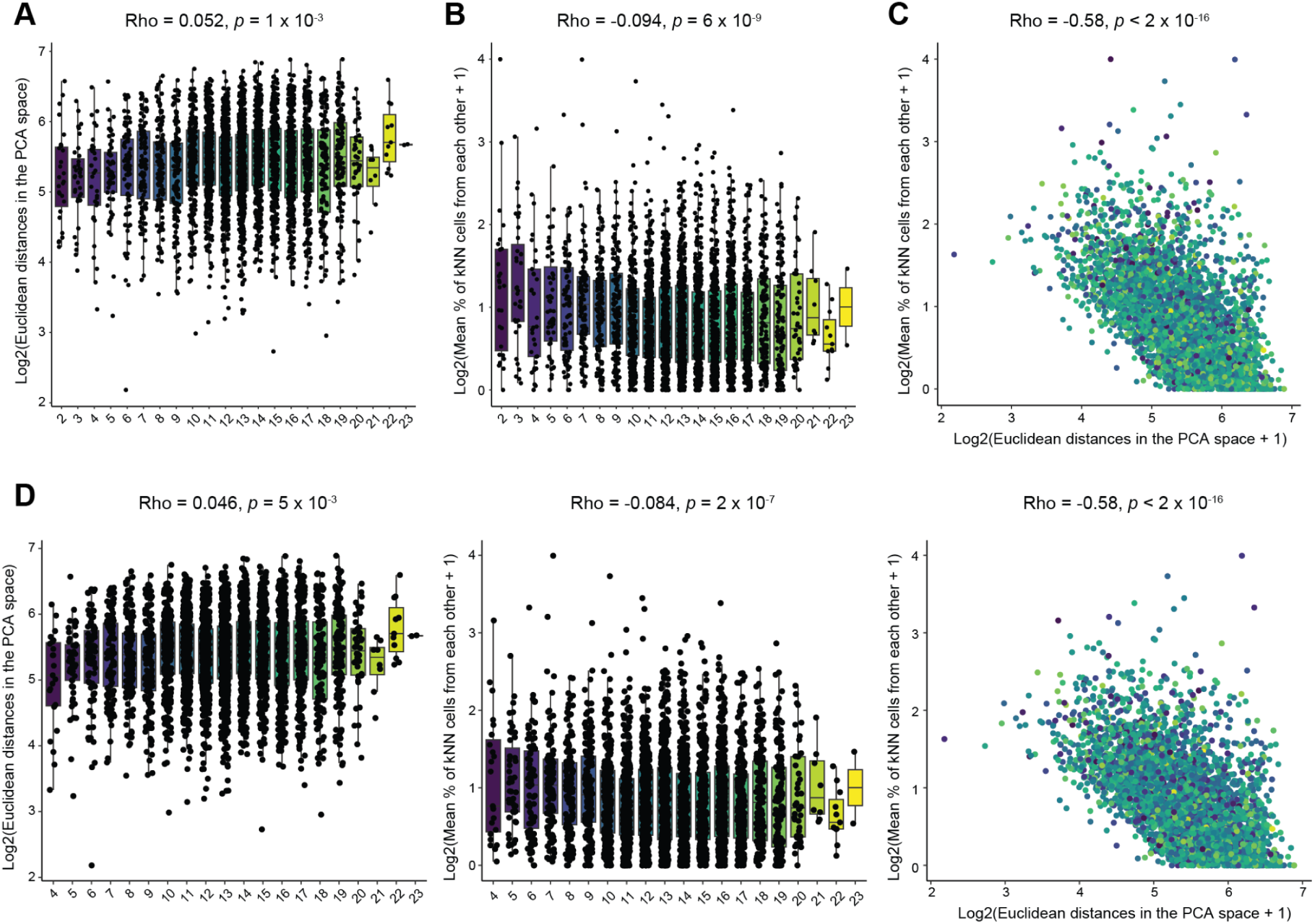
Pairwise distance and neighboring cell relationships among monoclonal gastruloids. **(A)** Pairwise Euclidean distances among 88 monoclonal gastruloids (≥50 cells assigned), calculated in 10-dimensional PCA space, are plotted against the “first epoch” lineage distances of their founder mESCs. Boxplots, encompassing a total of 3,828 pairs, represent IQR (25th, 50th, 75th percentile) with whiskers representing 1.5× IQR. Spearman correlation coefficient and p-value are indicated above. **(B)** For each gastruloid, the top 15 nearest neighboring cells for its cells were identified based on the 30-dimensional PCA space of the single-cell transcriptome. These neighboring cells were categorized by their gastruloid of origin, normalized by the total number of neighboring cells from that gastruloid, across all gastruloids. For each pair of gastruloids (e.g., A and B), the percentages of neighboring cells of A originating from B and those of B originating from A were averaged. Pairwise percentages of neighboring cells from each other among 88 gastruloids, are plotted against their lineage distances. Boxplots, encompassing a total of 3,828 pairs, represent IQR (25th, 50th, 75th percentile) with whiskers representing 1.5× IQR. Spearman correlation coefficient and p-value are indicated above. **(C)** The log2 pairwise Euclidean distances are plotted against the log2 pairwise percentages of neighboring cells for all 3,828 pairs. Spearman correlation coefficient and p-value are indicated above. **(D)** For each of the 88 gastruloids, the nearest neighbors were identified from the phylogenetic lineage tree, yielding 112 pairs. After removing redundancies (e.g., retaining only one of A vs. B and B vs. A), 76 pairs remained. These 76 pairs were excluded, and the analyses shown and described in panels **A–C** were repeated with the remaining 3,752 pairs.

